# Targeting Epsins by nanotherapy regulates lipid metabolism and promotes ABCG1-mediated cholesterol efflux to fortify atheroma regression

**DOI:** 10.1101/2022.08.09.503334

**Authors:** Kui Cui, Xinlei Gao, Beibei Wang, Hao Wu, Yunzhou Dong, Yuling Xiao, Xingya Jiang, Marina V. Malovichko, Kathryn Li, Qianman Peng, Yaowei Lu, Bo Zhu, Rongbin Zheng, Scott Wong, Douglas B. Cowan, MacRae Linton, Sanjay Srivastava, Jinjun Shi, Kaifu Chen, Hong Chen

## Abstract

**BACKGROUND:** Excess cholesterol accumulation in lesional macrophages elicits complex responses in atherosclerosis. Epsins, a family of endocytic adaptors, fuel the progression of atherosclerosis; however, the underlying mechanism and therapeutic potential of targeting Epsins remains unknown. In this study, we determined the role of Epsins in macrophage-mediated metabolic regulation. We then developed an innovative method to therapeutically-target macrophage Epsins with specially-designed S2P-conjugated lipid nanoparticles (NPs), which encapsulate small interfering RNAs to suppress Epsins.

**METHODS:** We used single cell RNA sequencing (scRNA-seq) with our newly developed algorithm MEBOCOST to study cell-cell communications mediated by metabolites from sender cells and sensor proteins on receiver cells. Biomedical, cellular and molecular approaches were utilized to investigate the role of macrophage Epsins in regulating lipid metabolism and transport. We performed this study using myeloid-specific Epsin double knockout (LysM-DKO) mice and mice with a genetic reduction of ABCG1 (LysM-DKO-ABCG1^fl/+^). The NPs targeting lesional macrophages were developed to encapsulate interfering RNAs to treat atherosclerosis.

**RESULTS:** We revealed that Epsins regulate lipid metabolism and transport in atherosclerotic macrophages. Inhibiting Epsins by nanotherapy halts inflammation and accelerates atheroma resolution. Harnessing lesional macrophage-specific NP delivery of Epsin siRNAs, we showed that silencing of macrophage Epsins markedly diminished atherosclerotic plaque size and promoted plaque regression. Mechanistically, we demonstrated that Epsins bound to CD36 to facilitate lipid uptake by enhancing CD36 endocytosis and recycling. Conversely, Epsins promoted ABCG1 degradation via lysosomes and hampered ABCG1-mediated cholesterol efflux and reverse cholesterol transport. In a LysM-DKO-ABCG1^fl/+^ mouse model, enhanced cholesterol efflux and reverse transport due to Epsin deficiency was suppressed by the reduction of ABCG1.

**CONCLUSIONS:** Our findings suggest that targeting Epsins in lesional macrophages may offer therapeutic benefits for advanced atherosclerosis by reducing CD36-mediated lipid uptake and increasing ABCG1-mediated cholesterol efflux.

**Novelty and Significance:** *WHAT IS KNOWN?:*

- Epsin endocytic adaptor proteins are upregulated in human and mouse atherosclerotic lesions
- Lesional macrophages internalize lipids primarily through scavenger receptor-mediated endocytosis such as CD36 and SR-A
- Macrophage-mediated cholesterol efflux and reverse cholesterol transport is crucial to atheroma resolution

*WHAT NEW INFORMATION DOES THIS ARTICLE CONTRIBUTE?:*

- ScRNA-seq combined with the newly-developed algorithm MEBOCOST reveals that Epsins are involved in macrophage-mediated lipid metabolic regulation
- Macrophage epsins promote lipid uptake by targeting CD36 endocytosis and membrane recycling via the Epsin ENTH domain
- Epsins bind ubiquitinated ABCG1—resulting in the endocytosis and lysosomal degradation of this cholesterol transporter, which reduces cholesterol efflux.
- Macrophage-specific, nanoparticle-mediated RNAi delivery exhibits a therapeutic benefit for the treatment of atherosclerosis
- Atherosclerotic plaque regression using this nanoparticle delivery platform represents a clinically-relevant approach for the treatment of advance atherosclerosis

*BRIEF SUMMARY:* Despite the development of new cholesterol-lowering therapies, including the recently approved PCSK9 small interfering RNA (siRNA) antagonists, patients still face a tremendous risk of developing major acute cardiovascular events resulting from chronic inflammation in the plaque. We employed a novel nanomedicine platform containing a stabilin-2 targeting peptide (S2P) to deliver Epsin-specific siRNAs to lesional macrophages. We discovered that inhibition of these adaptor proteins in lesional macrophages significantly diminished plaque size and necrotic core area, increased fibrous cap thickness, and promoted plaque regression.

## INTRODUCTION

Atherosclerosis is a complex and chronic condition that results in a buildup of arterial fatty deposits, called plaques that are the root cause of coronary heart disease, ischemic stroke, and peripheral artery disease ^1, 2^. Despite effective lipid-lowering therapies including new drugs of antibody and siRNA drugs targeting PCSK9, atherosclerosis is still the leading cause of mortality worldwide. Atherosclerotic lesion formation involves interactions between oxidized low-density lipoprotein (oxLDL) and prominent constituents of the arterial wall including monocyte-derived macrophages, immune cells and vascular smooth muscle cells ^3, 4^. Mounting evidence indicates that factors with the potential to promote macrophage egress from plaques (*e.g.,* regulatory T cells) play a critical role in regulating macrophage pro-resolving functions during atherosclerosis regression ^5^. Furthermore, the pro-inflammatory effects of oxLDL lead to endothelial activation and macrophage recruitment ^6^. Invading macrophages in the sub-endothelium ingest modified LDL and are transformed into lipid-laden foam cells ^1, 3^. This escalates arterial inflammation—a prime contributor to the transition from a stable to vulnerable atheroma ^7–10^. Accordingly, patients that receive a therapeutic cocktail aimed at alleviating dyslipidemia and hypertension still develop deleterious cardiovascular diseases ^2, 11^. This accentuates the urgent need for next-generation therapies geared toward eradicating inflammation in the arterial wall.

Epsins are a family of highly-conserved endocytic adaptor proteins that associate with the plasma membrane through interactions between their N-terminal homology (ENTH) domain and membrane PI (4,5) P2 ^12–14^, where it recognizes and recruits ubiquitinated cell surface receptors to clathrin-coated pits for internalization via its ubiquitin-interacting motif (UIM) and clathrin/AP-2 binding sites, respectively ^15–18^. Epsins 1 and 2 are ubiquitously expressed, redundant in function, and essential for embryonic survival ^19^. The embryonic lethality of Epsin 1 and 2 double knockout mice (DKO) prompted us to generate conditional Epsin1^fl/fl^; Epsin2^-/-^ mice. Combining DKO strains with specific Cre deleter mouse strains creates tissue and cell type-specific DKO mice ^19, 20^. Despite their multifaceted functions, our previous work demonstrates that Epsins possess a vast degree of specificity and selectivity in choosing their binding partners. Therefore, the actions of Epsins are cell context-dependent ^19–26^. Notably, we demonstrate that myeloid-specific Epsin loss downregulates inflammation and impedes atheroma formation in an atherogenic mouse model by stabilizing macrophage LRP-1 ^26^. However, this model failed to address the role of epsins in hindering atheroma regression and promoting plaque rapture and consequent debilitating episodes.

Lesional macrophages internalize lipids primarily through scavenger receptor-mediated endocytosis ^27^, which is also responsible for foam cell formation ^28–32^. Mice lacking both Scavenger Receptor-A (SR-A) and CD36 revealed that CD36 is the major oxLDL receptor required for foam cell formation ^29–32^. Our previous studies revealed that Epsins 1 and 2 are upregulated in lesional macrophages ^26^. Epsin-deficient (DKO) macrophages exhibited a striking reduction in foam cell formation ^26^. As robust lipid uptake by macrophages is one of major responsible factors that drive foam cell formation, we posit a central role for Epsins in regulating lipid uptake in macrophages; however, how Epsins facilitate lipid uptake and whether this occurs through scavenger receptor-mediated uptake of modified LDL during foam cell formation remains unknown.

Reverse cholesterol transport (RCT) is responsible for removing excess cholesterol from peripheral tissues to the liver, where it is reused or removed from the body ^33^. A critical part of RCT is cholesterol efflux, where accumulated cholesterol removed from macrophages in the subintima is mediated by ATP-binding membrane cassette transporters (ABCA1 and ABCG1) and other molecules ^34–36^. Among the regulators of RCT, ABCG1 plays a prominent role in macrophage cholesterol and phospholipid transport to maintain cellular lipid homeostasis by using high density lipoprotein (HDL) as a cholesterol acceptor ^37^. HDL is thought to facilitate macrophage re-programming to be less inflammatory through an ATF-3-dependent pathway ^38^. Several clinical trials and more recent work suggest that increased HDL-cholesterol concentrations fail to improve cardiovascular disease outcomes ^39–41^; yet, new data indicate that rather than the steady-state HDL-cholesterol (HDL-C) levels ^42^, the ability of HDL particles to transport cholesterol from atherosclerotic plaques to the liver is more important ^43, 44^. Consequently, a shift in focus from increasing HDL-cholesterol concentrations to raising RCT function by increasing cholesterol efflux capacity (CEC), the HDL-ABCG1-mediated cholesterol efflux from macrophages, has become a prospective therapy for coronary artery disease (CAD), especially in severe acute myocardial infarction (AMI) patients ^45–48^. As lack of effective cholesterol efflux in macrophages hinders the reversal of foam cells to healthy macrophages, we envisioned a major role for macrophage Epsins in regulating cholesterol efflux. Given the importance of cholesterol efflux and RCT in atheroma resolution and the aforementioned risk reduction in AMI patients ^34–36^, it is crucial to determine whether Epsins inhibit cholesterol efflux and RCT to promote atherosclerotic plaque progression and impede its regression.

In this study, we harnessed innovative single-cell bioinformatics technology and therapeutic nanotechnology to better understand atherosclerosis. This allowed us to determine the detrimental role of epsins in prohibiting atheroma regression and promoting plaque rupture using a preclinical atherosclerosis regression model. Mechanistically, we utilized scRNA-seq analysis of macrophages and newly-developed bioinformatic techniques to identify new targets involved in lipid metabolism during atherosclerosis. Based on these new targets, we generated a myeloid-specific-deficient ABCG1 murine model to uncover how epsins regulate lipid metabolism to propel atherosclerosis. Specifically, we show that Epsins facilitate CD36-mediated lipid uptake and inhibit ABCG1-mediated cholesterol efflux, which is an entirely new phenomenon in the field of lipid metabolism. We also employed a novel macrophage-specific nanoparticle delivery system to investigate the therapeutic treatment of atherosclerosis. Lastly, using an early-stage progression model, an advanced-stage progression model, and a regression model of atherosclerosis we established the therapeutic benefit of treating this disease with a nanoparticle delivery system. This rigorous and comprehensive mechanistic study on lipid metabolism, and clinically-relevant nanoparticle-mediated RNAi therapies represents a potential breakthrough for treating this devastating disease.

## METHODS

Detailed Methods and materials are available in the Supplemental Materials. Antibodies, primers, and reagents are listed in Supplemental Table S1.

### Animal models

In this study, all animal procedures were performed in compliance with institutional guidelines and mouse protocols were approved by the Institutional Animal Care and Use Committee (IACUC) of Boston Children’s Hospital, MA, USA. Both male and female mice were used. C57BL/6 mice (stock #00664), ApoE^-/-^ mice (stock #008525), LysM-Cre deleter mice (stock #004781), and ABCG1^flox^ mice (C57BL/6-Abcg1^tm1Ched^/J, stock #027954) were purchased from Jackson Research Laboratory. As double knockout of Epsins 1 and 2 in mice lead to embryonic lethality, we generated Epsin1^fl/fl^;Epsin2^-/-^ mice. These mice were subsequently combined with LysM-Cre deleter mouse strains to create myeloid-specific Epsins deficient (LysM-DKO) mice and bred them to the C57BL/6 background ^26^. As proper controls, we also generated wild type C57BL/6 mice bearing LysM Cre. These mice only have one copy of LysM Cre as homozygous LysM Cre mice are susceptible to atherosclerosis. Furthermore, these mice were crossed to the ApoE^-/-^ (C57BL/6) background to obtain ApoE^-/-^/WT and ApoE^-/-^/LysM-DKO mice ^26^. We used macrophages from WT and LysM-DKO on either normal background or ApoE^-/-^ background. We have not seen significant differences in the results with these backgrounds. ***For simplicity, we referred to the macrophages from WT or ApoE^-/-^/WT as WT macrophages and macrophage from LysM-DKO and AopE^-/-^/LysM-DKO as DKO macrophages.*** We used these mice (male and female) and primary macrophages (peritoneal and bone marrow derived) isolated from them for this study.

In addition, to generate myeloid-specific ABCG1-KO mice, we crossed the LysM-cre^+^ mice with ABCG1^fl/fl^ mice. We then bred the obtained LysM-cre^+^; ABCG1^fl/fl^ mice with ApoE^-/-^ mice and backcrossed them on to the C57BL/6 background.

To induce atherosclerosis, mice were fed Western diet (WD, Protein 17% kcal, Fat 40% kcal, Carbohydrate 43% kcal; D12079B, Research Diets, New Brunswick, USA) starting at the age of 6-8 weeks for 8-20 weeks. Mice were sacrificed at different time points based on the experimental design and peritoneal macrophages, blood, heart, aorta and bone marrow monocytes were harvested. For control mice, in addition to ApoE^-/-^;Epsin1^+/+^;Epsin2^+/+^ mice, we also used ApoE^-/-^;Epsin1^+/+^;Epsin2^+/+^ mice with a single copy of LysM-cre, and ApoE^-/-^;Epsin1^fl/fl^;Epsin2^-/-^ littermates lacking the single copy of LysM-cre. To simplify the terminology, we refer to these control mice as ApoE^-/-^, as results were not different in any of the analyses we performed. For the study of atheroma resolution, WT mice (both male and female) at the age of 8 weeks were intravenously injected with 2×10^11^ genomes of PCSK9 adeno-associated virus (rAAV8-D377Y- mPCSK9 purchased from Boston Children’s Hospital Viral Core Facility) followed by 17 weeks of WD feeding. For each experimental model and time point, 6-10 mice were analyzed and both male and female mice were used in separate groups.

### Aortic single-cell preparation and single-cell RNA (scRNA) sequencing

WT and DKO mice were euthanized by CO2 inhalation. The aortas were isolated after perfusion with 30 mL of PBS through left ventricular and quickly transferred to cold DMEM medium. Aortas from the two groups were cut into about small pieces and digested with an enzyme solution (5mg/mL collagenase type I, 5mg/mL collagenase type IV, and 5mg/mL liberase) for 90min at 37 °C on a shaker. The digested cell suspension was filtered through a 40 μm strainer and washed twice with PBS. The cells were resuspended and ready for sequencing in PBS with 0.04% bovine serum albumin, and their viability was over 90%.

Single-cell RNA-Seq library construction was performed using the protocol provided by 10X Genomics. In brief, the single-cell suspensions from both groups, reagents, gel beads and partitioning oil were loaded to 10X Chromium Chip G to generate single-cell Gel Beads-in-emulsion (GEMs, Single cell 3’ Reagent Kits v3.1, 10X Genomics). scRNA was barcoded through reverse transcription in individual GEMs followed by a post GEM-RT cleanup and cDNA amplification. Then, a 3’-gene expression library construction was performed. Finally, the library was sent for sequencing.

### ScRNA-seq data analysis and metabolite-sensor communication inference

The raw scRNA-seq data were processed using Cell Ranger (version 6.1.2) (10x Genomics). The reads were mapped to the prebuilt mouse mm10 genome. The resulted gene expression matrix in individual single cells was processed by the R package Seurat (version 4.1.0) ^49^. Low-quality cells with number of expressed genes less than 200 or larger than 5000 or with percentage of mitochondria reads greater than 10% were dropped out. Rarely expressed genes which were detected in less than 3 cells were removed. Mitochondria genes and ribosomal protein coding genes were removed from the expression matrix before normalization. The high-quality data after these filtering steps was then used at additional processing steps including log normalization, data scaling, principal component analysis (PCA), cell clustering, and UMAP visualization. The cell clusters were visualized by the DimPlot function. Differentially expressed genes among cell clusters were identified by FindAllMarkers function using the default Wilcoxon test method, with minimal percentage of expressed cells as 25% and minimal log2 fold change as 0.25. Next, marker genes in each cell clusters were used to annotate cell types based on known marker genes in PanglaoDB database ^50^ and literatures. The marker gene expressions were visualized by DotPlot and VlnPlot function. Trajectory analysis was conducted by Monocle3 ^51, 52^. The metabolite-sensor cell-cell communication was analyzed by MEBOCOST ^53^. The data was analyzed following the tutorial on the MEBOCOST website (https://github.com/zhengrongbin/MEBOCOST). scRNA- seq expression data was used to estimate metabolite abundance and calculate communication score for each condition. Next, results of two conditions were combined to compare the communications. The differences in communication scores between two conditions were calculated. The prediction of sender-metabolite-sensor-receiver communication events were visualized by barplot, flow plot and circle plot. The metabolite abundance and sensor expression levels were exhibited by violin plot. Index of dispersion (IOD) was calculated using communication scores across conditions as described in the MEBOCOST paper^53^. The top 100 most variable communications were selected for further investigation. All the data for scRNA-seq are available in Data files (S1-S2).

### Human samples

Human healthy control (n=3) and diseased aortic arch samples (n=3) from atherosclerosis patients were purchased from Maine Medical Center Biobank. The medical information of the atherosclerotic patient samples is in Data file S9. The paraffin sections were de-paraffinized and performed antigen retrieval to unmask the antigenic epitope with 10mM Sodium Citrate, pH 6.0, with 0.5% Tween 20 at 90°C for 10 minutes. Immunofluorescence staining of the slides was performed with the standard protocol described below.

### Synthesis of DSPE-PEG-S2P, preparation and characterization of S2PNP-siRNA

To construct the lesional macrophage-targeted siRNA NPs, S2P peptide-conjugated DSPE-PEG (DSPE-PEG-S2P) was first synthesized via a thiol-maleimide Michael addition click reaction between S2P peptide (CRTLTVRKC, GLS Biochem Systems Inc.) and DSPE-PEG-Mal [PEG molecular weight, 3.4 kDa; Nanocs Inc.], as reported previously ^54^. Then, a robust self-assembly method was used to prepare the targeted polymer-lipid hybrid NPs for siRNA delivery ^54, 55^. In brief, G0-C14 and PLGA were dissolved separately in anhydrous dimethylformamide (DMF) to form a homogeneous solution at the concentration of 2.5 mg/mL and 5 mg/ml, respectively. DSPE-PEG- OCH3 (DSPE-mPEG) and DSPE-PEG-S2P were dissolved in HyPure water (GE Healthcare Life Sciences, catalog no. SH30538) at the concentration of 0.1 mg/mL. 1 nmol Epsin1 siRNA and 1 nmol Epsin2 siRNA were gently mixed with 100 μL of the G0-C14 solution. The mixture of siRNA and G0-C14 was incubated at room temperature for 15 min to ensure the full electrostatic complexation. Next, 500 μL of PLGA polymers were added and mixed gently. The resultant solution was subsequently added dropwise into 10 mL of HyPure water containing 1 mg lipid- PEGs (i.e., 50% DSPE-PEG-S2P and 50% DSPE-mPEG hybrids for the S2P-targeted siRNA NPs, or 100% DSPE-mPEG for the non-targeted siRNA NPs) under magnetic stirring (1,000 rpm) for 30 min. The siRNA NPs were purified by an ultrafiltration device (EMD Millipore, MWCO 100 kDa) to remove the organic solvent and free excess compounds via centrifugation at 4 °C. After washing 3 times with HyPure water, the siRNA NPs were collected and finally resuspended in pH 7.4 PBS buffer. The NPs were used freshly or stored at -80 °C for further use. The physicochemical properties (particle size and surface charge) of S2PNP-siEpsin1/2 were characterized by dynamic light scattering (DLS, Brookhaven Instruments Corporation). The S2PNP-siEpsin1/2 was ∼89 nm in size as measured by DLS, and their surface charge was determined to be ∼ -5.3 mV.

### Statistical analysis

Statistical analysis was processed using GraphPad Prism 8.4. Data are shown as mean ± standard deviation (mean ± SD). Normality was determined using D’Agostino-Pearson normality testing for experiments with n ≥ 8 and Kolmogorov-Smirnov normality testing for experiments with n < 8. Data were analyzed by the unpaired Student’s t-test or Mann-Whitney U test where appropriate for two groups. Data for multiple groups were analyzed with ANOVA with Bonferroni test or Dunn’s multiple comparison test where appropriate. P value less than 0.05 was considered significant.

## RESULTS

### Single-cell RNA-seq revealed that macrophage Epsins regulate cholesterol metabolism and efflux pathways

For an unbiased analysis of all pathways regulated by Epsins in macrophages, we performed single-cell RNA sequencing (scRNA-seq) analysis of aortic tissue isolated from wild type (WT) and macrophage-specific DKO mice. We identified a diverse range of cell types, including vascular smooth muscle cells (VSMCs), fibroblasts, endothelial cells, macrophages, and other cells (Figure 1A). Known cell markers facilitated the annotation of cell types, such as the Tagln and Acta2 for VSMCs, Pecam1 and Cdh5 for endothelial cells, Cd14 and Cd68 for macrophages^50, 56, 57^ (Figure S1A). Macrophage populations were further subclustered to investigate their heterogeneity among subpopulations (Figure 1B). Multiple well-known macrophage subpopulations were recaptured, including M1 and M2 macrophages. Markers reported in the literature were used to annotate the subtypes(*e.g.,* Mrc1 (Cd206) and Cd163 for M2 macrophages)^58^ (Figure S1B and S1C). The composition of the aortic cell population showed striking differences between WT and DKO aortic cells (Figure S2).

**Figure 1.**
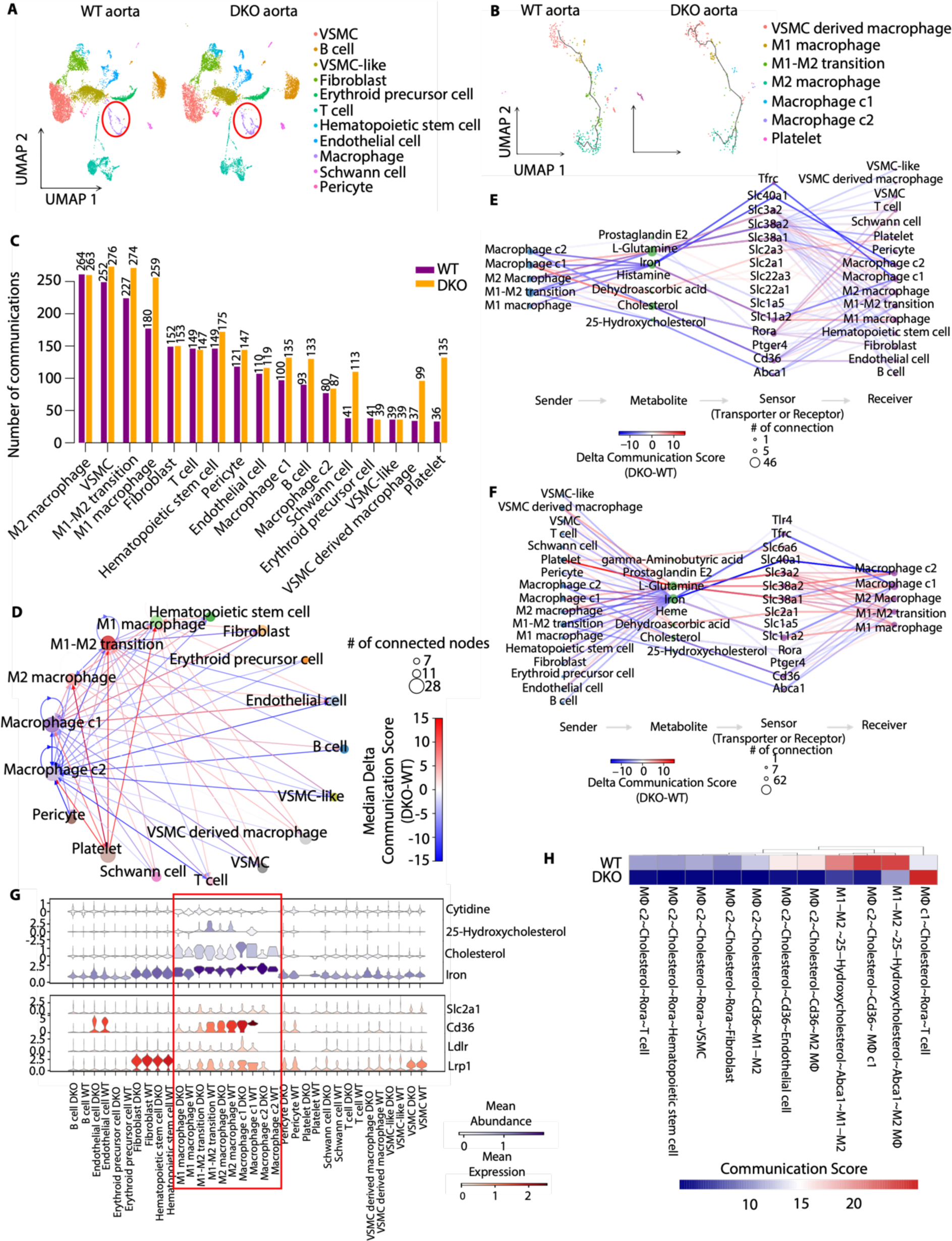
scRNA-seq data revealed down-regulation of cholesterol-related cell-cell communications mediated by metabolites and their sensor proteins in macrophage subpopulations in DKO aorta compared to WT. (**A**) UMAP plot of cell clusters in WT and DKO aorta. Macrophage populations are indicated in the red circle. (**B**) UMAP of subclusters of macrophage populations in (A). Trajectory inferred by Monocle3 was displayed. (**C**) A barplot showing the number of communication events in cell groups in WT and DKO. (**D**) A circle plot showing the differential communications between cell groups in DKO compared to WT. The arrows indicate directions of communication from sender cells to receiver cells. The size of nodes positively correlates with the number of connected nodes. Line colors indicate the difference in communication scores of DKO compared to WT. (**E**-**F**) A flow plot of communications comprising senders, metabolites, sensors, and receivers from macrophage subpopulations to all the cell types (E); and from all cell types to macrophages (F). (**G**) Violin plots of the abundance of representative metabolites and sensors across cell types in WT and DKO. Metabolite abundance (upper), sensor abundance (below). (**H**) A heatmap showing the most variable communications related to cholesterol metabolism in macrophages in the comparison of WT and DKO aorta.

The molecular mechanisms underlying Epsin-mediated regulation of foam cell formation and promotion of lipid uptake by macrophages are poorly understood. We therefore employed the algorithm MEBOCOST, which is the first algorithm developed to detect cell communications mediated by metabolites and their sensor proteins such as receptors and transporters (metabolite-sensor cell communications) ^53^. The results showed that macrophage subpopulations and VSMCs were the top cell types ranked by the number of metabolite-sensor cell communications from the largest to the smallest (Figure 1C). Furthermore, the numbers of metabolite-sensor cell communications in these cell populations increased in DKO compared to WT (Figure 1C). Macrophage subpopulations were found to not only send metabolites to other cell types, but also receive metabolites from other cell types. They could send and receive metabolites in an autocrine manner as well (Figure 1D). The communications between a pair of cell populations can be mediated by multiple metabolite-sensor pairs. The overall changes in communication were estimated by the median differences of communication scores between any two cell groups. We observed widespread changes of cell-cell communications in DKO when compared to WT, with both strengthened and weakened communications (Figure 1D). To gain a deeper insight into the changed metabolite-sensor cell communications related to macrophages, we inspected individual sender cell types of metabolites and receiver cell types of sensor proteins. First, we observed a decrease of cholesterol-Cd36 cell communications from macrophages, especially in macrophage cluster 2 (c2), to other cell types in DKO compared to WT (Figure 1E); second, cholesterol-Cd36 cell communications from macrophage c2 to macrophage c1 were reduced by the DKO (Figure 1F). As for the specificity of metabolite and sensor abundance in different cell types, we observed that cholesterol and its derivative, 25-hydroxycholesterol, were highly-enriched in macrophage populations; Cd36, the receptor of cholesterol, was also highly expressed in macrophage populations (Figure 1G). The cell communications through cholesterol and Cd36 in macrophages was supported by their abundance in these cells. Finally, the most variable communications in macrophages mediated by cholesterol appeared to be mostly decreased rather than increased (Figure 1H).

We subsequently performed bulk RNA-seq analysis of macrophages and revealed that loss of Epsins 1 and 2 significantly altered the expression of 2208 genes (Figure S3). These include 1138 downregulated and 1070 upregulated genes (Figure S3A). Notably, the differential genes are associated with cholesterol efflux and metabolism, suggesting that Epsins play a role in regulating lipid cholesterol metabolism and efflux pathways (Figure S3B). Consistently, Gene Set Enrichment Analysis (GSEA) showed that genes related to fatty acid metabolism tended to be markedly downregulated in DKO macrophages (Figure S3C). In contrast, genes associated with cholesterol transport and efflux (Figure S3D), and negative regulation of macrophage foam cell formation (Figure S3E), tended to be upregulated by Epsin deficiency.

We next compared our DKO macrophage RNA-seq data with a public RNA-seq data from wild type and CD36 knockout (CD36KO) macrophages in the public database ^59^. We observed a significant overlap of differential genes induced by the Epsin-DKO and CD36KO mouse macrophages. There were 169 genes significantly upregulated by both Epsin-DKO and CD36KO (4-fold enrichment compared to the 41 genes expected by random chance, Fisher’s exact test P value 9.73 x 10^-25^) (Figure S3F left panels); 29 genes (6-fold enrichment, Fisher’s exact test P value 6.15 x 10^-9^) were downregulated by both Epsin-DKO and CD36KO (Figure S3F right panels). GO analysis revealed multiple pathways regulated by both Epsin or CD36 deficiency. These include the downregulation of inflammation pathways, lipid biosynthetic processes, triglyceride metabolic processes, and upregulation of the small GTPase-mediated signal transduction pathway (Figure S3G). The results indicated that Epsins and CD36 may have superimposing roles in regulating molecular pathways involved in lipid metabolism and inflammation.

### Epsin promotes CD36-mediated lipid uptake by promoting CD36 endocytosis and recycling

Based on the bioinformatic results, we hypothesized that Epsins regulate CD36 internalization and thus lipid uptake. We performed qRT-PCR, western blot (WB) and flow cytometry to check the RNA, total protein, and surface protein levels of CD36. Interestingly, RNA (Figure 2A) and total protein (Figure 2B) expression of CD36 remained unchanged between WT and DKO macrophages; however, the surface protein level of CD36 significantly decreased with the treatment of oxLDL in clathrin-dependent manner in WT, but not in DKO macrophages (Figure 2C and S4A-C). Immunofluorescence staining (IF) revealed oxLDL-induced CD36 trafficking to early endosomes. We observed colocalization of CD36 with EEA1 (early endosome antigen 1, Figure 2D) and Rab11 (recycling endosome marker, Figure 2E) in WT macrophages treated with oxLDL, which was not observed in DKO macrophages.

**Figure 2.**
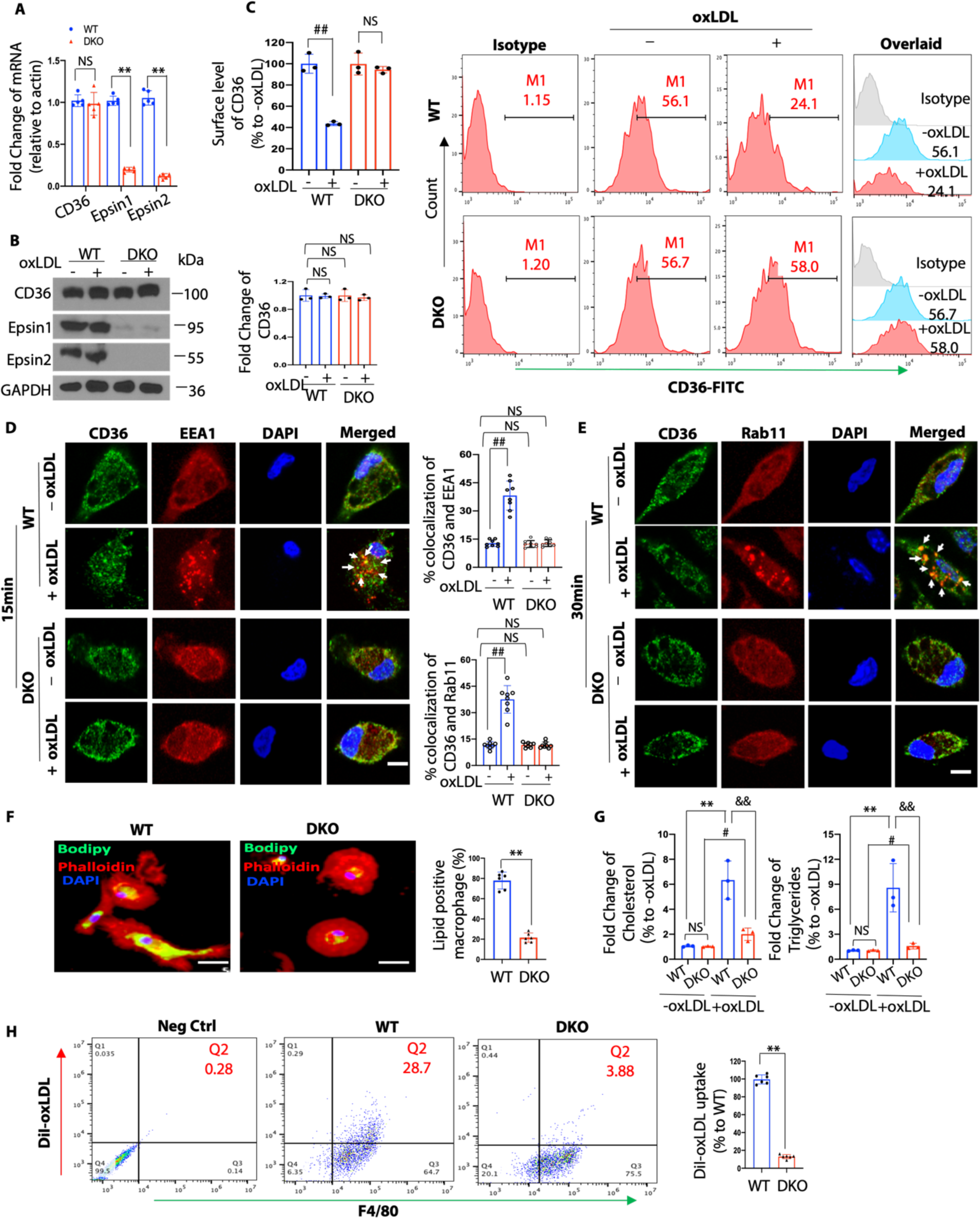
Epsin facilitates CD36-mediated lipid uptake by promoting CD36 endocytosis and recycling. (**A-C**) Bone marrow derived macrophages (BMDM) or thioglycolate induced peritoneal macrophages from WT (n=5) and DKO (n=5) mice were incubated in lipid-deficient medium for 24h and treated with or without 100μg/mL oxLDL for 1h at 37^0^C. qRT-PCR analysis of CD36 expression (A), western blot (WB) analysis for total protein level of CD36 (B) and flow cytometry for surface level of CD36 (C). NS, no significant difference; **WT vs DKO group, P<0.01; ^##^-oxLDL vs +oxLDL group, P<0.01. (**D-E**) Elicited WT and DKO macrophages were incubated in lipid-deficient medium for 24h and treated with or without 100μg/mL oxLDL for 15min (D) or 30min (E) at 37^0^C. Macrophages were co-stained with CD36 (green), the early endosome marker EEA1 (red) or the recycling endosome marker Rab11 (Red) and DAPI (bule), and imaged using confocal microscope. White arrows indicate the endocytic vesicles, scale bar=5μm, ^##^-oxLDL vs +oxLDL group (n=8), P<0.01. (**F**) BODIPY staining of peritoneal macrophages pre-incubated with 25μg/mL oxLDL for 24h in lipid-deficient medium, **WT vs DKO group (n=6), P<0.01, scale bar=10μm. (**G**) Cholesterol and triglycerides levels in WT and DKO macrophages treated with 25μg/mL oxLDL for 24h in lipid-deficient medium (n=3). **- oxLDL (WT) vs +oxLDL (WT), P<0.01; ^#^-oxLDL (DKO) vs +oxLDL (DKO), P<0.05; ^&&^ +oxLDL (WT) vs +oxLDL (DKO), P<0.01. (**H**) Macrophages isolated from WT and DKO mice were incubated in lipid-deficient medium for 24h followed by the treatment with DiI-oxLDL for 2h at 37^0^C to assess the lipoprotein uptake by flow cytometry, **WT vs DKO group (n=6), P<0.01. Data from A-H are presented as mean ± SD and were analyzed using either unpaired Student’s t- test (two groups) or one-way ANOVA (multiple comparisons).

To further study the role of Epsins in regulating macrophage lipid uptake, we stained oxLDL-treated WT and DKO macrophages with BODIPY and Oil Red O (ORO). Deficiency of Epsins impaired oxLDL uptake and foam cell formation (Figure 2F and S5A-B) and reduced cholesterol and triglyceride levels (Figure 2G) in DKO macrophages. Flow cytometry confirmed that macrophage lipid uptake was reduced in DKO macrophages using DiI-oxLDL treatment (Figure 2H and S4B). Together, these results indicate that Epsins are crucial for CD36-mediated lipid uptake and cholesterol metabolism.

### Epsin ENTH domain is required for CD36-mediated lipid uptake and foam cell formation

Next, we investigated how Epsins and CD36 interact to cause the disparity in CD36 surface and total protein levels using immunoprecipitation (IP) and WB. We determined that CD36 binds Epsin1 in WT, but not in DKO macrophages (Figure 3A). This indicated that Epsin1 and CD36 interact endogenously. We then determined which Epsin regions were responsible for this interaction. We created Flag-tagged mammalian Epsin1 wild type full length and ENTH or UIM deleted constructs (FLAG-Epsin1^WT^, FLAG-Epsin1^△ENTH^ or FLAG-Epsin1^△UIM^) (Figure 3B). We transfected these constructs to HEK 293T cells and performed co-IP with CD36. The interaction between Epsin1 and CD36 was dependent on the ENTH domain as the binding between Epsin1^△ENTH^ and CD36 was dramatically reduced in FLAG-Epsin1^△ENTH^-expressing cells (Figure 3B). As expected, we did not detect ubiquitinated CD36 in response to oxLDL treatment in the presence of the proteasome inhibitor MG132 (Figure 3C). In addition, we transfected the Epsin full length and deletion Epsin constructs into WT and DKO macrophages treated with oxLDL. The transfection of full-length Epsin restored lipid uptake in DKO macrophages, which was not observed in cells expressing FLAG-Epsin1^△ENTH^ (Figure 3D, 3E and S6).

**Figure 3.**
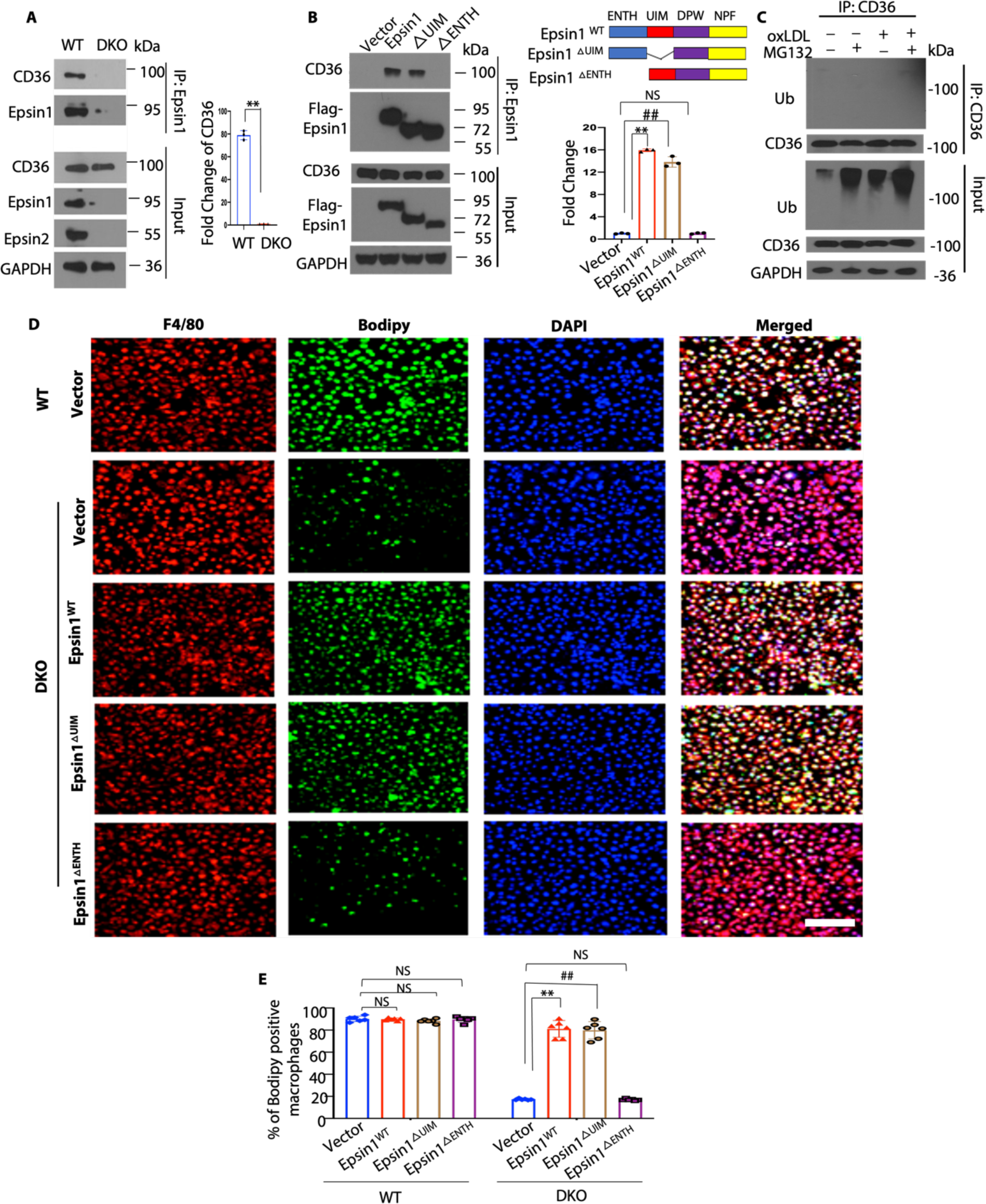
Epsin interacts with CD36 via Epsin-ENTH domain which is critical for CD36- mediated lipid uptake and foam cell formation. (**A**) BMDM or peritoneal macrophages isolated from WT and DKO mice were incubated in lipid-deficient medium for 24h followed by the treatment with 100μg/mL oxLDL for 1h, IP and WB analysis of Epsin1 and CD36 (n=3, **WT vs DKO, P<0.01). (**B**) CD36 plasmids and full length (FLAG-Epsin1^WT^) or domain-deletion constructs (FLAG-Epsin1^△ENTH^ or FLAG-Epsin1^△UIM^) in the pcDNA3 vector were transfected into HEK 293T cells for 48 h and then treated with 100μg/mL oxLDL for 1h, followed by immunoprecipitation (IP) and WB analysis using antibodies against FLAG tags and CD36 (n=3, **Epsin1^WT^ vs vector group, P<0.01; ^##^Epsin1^△UIM^ vs vector group, P<0.01). (**C**) Peritoneal macrophages isolated from WT mice were incubated in lipid-deficient medium for 24h followed by treatment with 5 μM MG132 for 3h. Cells were subsequently treated with 100 μg/mL oxLDL for 1h followed by IP and WB for ubiquitin and CD36. (**D**) FLAG-Epsin1^WT^, FLAG-Epsin1^△ENTH^, and FLAG-Epsin1^△UIM^ constructs were transfected into DKO macrophages for 48 h and treated with 100μg/mL oxLDL for 1 h, followed by staining with F4/80 (red), BODIPY (green) and DAPI (blue). Scale bar= 200μm. (**E**) Statistics for (D and Figure S6), n=6, **Epsin1 ^WT^ vs vector group, P<0.01; ^##^Epsin1^△UIM^ vs vector group, P<0.01. Data from A-D are presented as mean ± SD and were analyzed using either unpaired Student’s t-test (two groups) or one-way ANOVA (multiple comparisons).

### RNA-seq analyses of WT, Epsin-DKO and ABCG1-KO macrophages reveal that Epsins and ABCG1 reciprocally regulate inflammation and lipid metabolism pathways

Emerging evidence supports raising RCT function by increasing HDL-cholesterol efflux capacity (CEC) to treat recurrent heart attacks. We have generated myeloid-specific ABCG1- knock out (ABCG1-KO) mice. Given the intimate relationship of HDL with ABCG1 in regulating cholesterol efflux, and the involvement of Epsins in cholesterol efflux regulation, we performed RNA-seq analysis of WT, ABCG1-KO and Epsin-DKO macrophages. The loss of ABCG1 in macrophages significantly altered the expression of 1733 genes, including 986 downregulated and 747 upregulated genes (Figure S7A-S7C). GSEA indicated that genes associated with the inflammatory response (Figure 4A) and macrophage activation (Figure 4B) are upregulated by ABCG1-KO in macrophages. In contrast, genes negatively regulating macrophage-derived foam cell formation are downregulated by ABCG1-KO (Figure 4C). We then performed GO enrichment analysis for genes up- and down-regulated in ABCG1-KO relative to WT macrophages. Positive regulation of cytokine production, tumor necrosis factor production and acute inflammatory response were upregulated by ABCG1-KO in macrophages (Figure 4D). Negative regulation of lipid storage and cholesterol biosynthetic processes were down regulated in ABCG1-KO compared to WT macrophages (Figure 4D). Interestingly, GSEA analyses showed that genes associated with cell immune responses (Figure S7D), regulation of leukocyte migration (Figure S7E) and leukocyte proliferation (Figure S7F) are markedly upregulated by ABCG1 deficiency in macrophages. In contrast, these genes are downregulated in DKO macrophages (Figure S7D-S7F), These results suggest that Epsins and ABCG1 reciprocally regulate inflammation pathways. Examination of individual gene expression confirmed common target genes reversely regulated by Epsins and ABCG1. For instance, Itgb3 prevents the differentiation of macrophages and monocytes into foam cells ^60^ and it was up and down regulated by DKO and ABCG1-KO, respectively (Figure 4E). In addition, GO enrichment analysis revealed pathways conversely regulated by ABCG1-KO and DKO in macrophages (Figure 4F). Therefore, our RNA-seq analyses from ABCG1-KO and Epsin-DKO further supports our findings that Epsins and ABCG1 play opposing roles in regulating cholesterol efflux, inflammatory responses, and foam cell formation.

**Figure 4.**
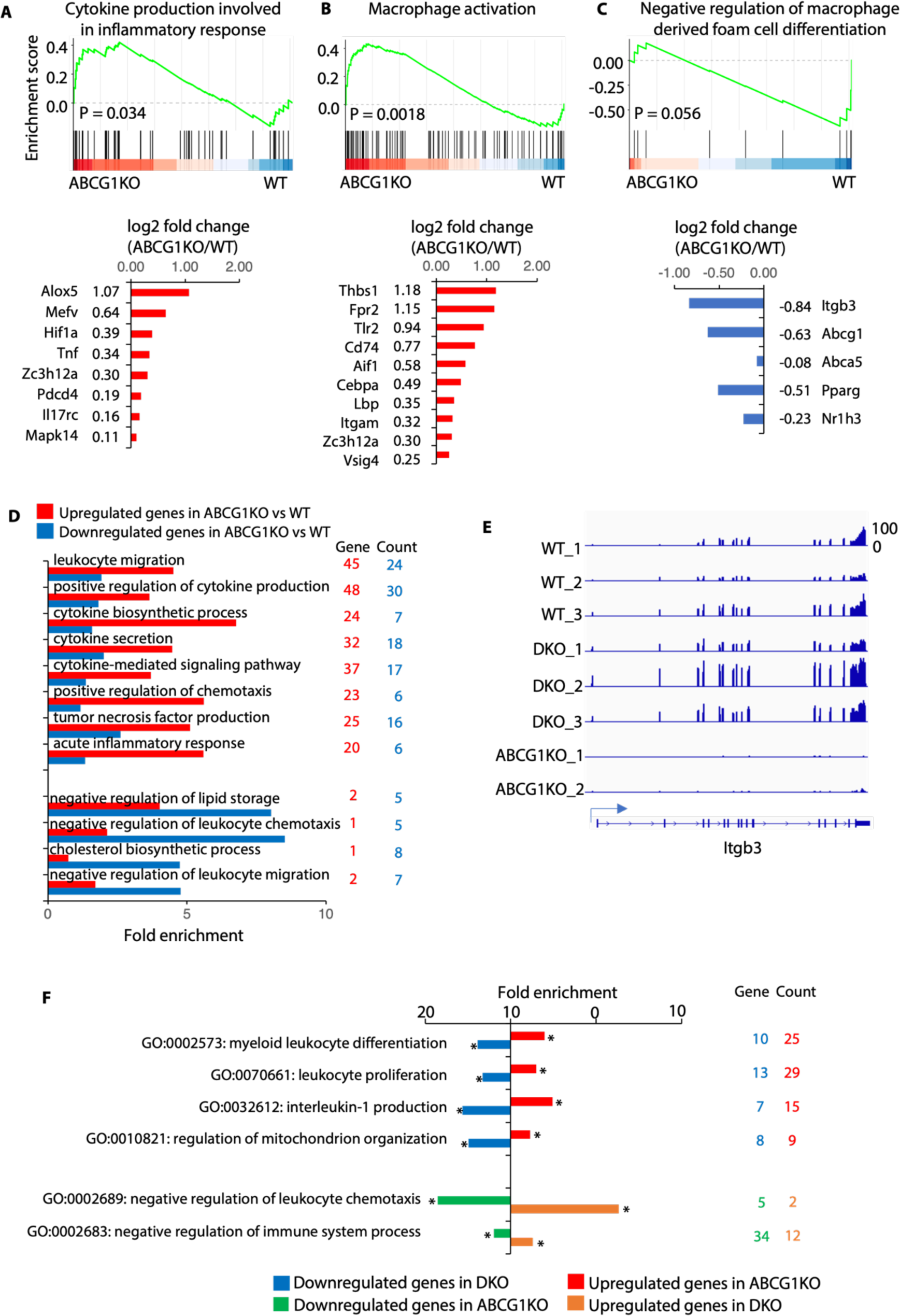
RNA-seq analyses of WT, DKO and ABCG1 knockout (ABCG1KO) macrophages indicate inverse regulation of lipid metabolism and inflammatory response. (**A-C**) The Gene Set Enrichment Analysis (GSEA) (top panels) indicates the tendency of individual pathways to be up or down regulated in ABCG1KO macrophages compared to wild type. Genes associated with cytokine production involved in the inflammatory response (A), macrophage activation (B), and negatively regulating macrophage derived foam cell differentiation (C) are analyzed. The Barplots (bottom panels) showing log2 fold changes of expression of altered genes. (**D**) GO enrichment analysis for up- and down-regulated genes in ABCG1KO relative to WT. (**E**) Genome browser tracks to show expression of Itgb3 in individual samples. (**F**) The GO enrichment analysis revealed pathways reversely regulated in ABCG1 knock out and Epsin deficient macrophages, compared to wild type. *P<0.05.

### Epsin loss augments macrophage cholesterol efflux *in vitro* and RCT *in vivo*

RNA-seq analysis of WT, DKO and ABCG1-KO macrophages indicated that the loss of Epsins affects cholesterol efflux. At the same time, we wanted to determine if Epsins inhibit cholesterol efflux and RCT to ameliorate atherosclerosis. WT and DKO macrophages were pre-treated with a liver X receptor (LXR) agonist (T0901317) and incubated with [^3^H]-cholesterol and ac-LDL. We found that *in vitro* cholesterol efflux to HDL, but not ApoA-1, was markedly enhanced in DKO versus WT macrophages (Figure 5A and 5B). Consistently, *in vivo* RCT (Figure 5C) was elevated in mice injected with [^3^H]-cholesterol-loaded DKO macrophages compared to WT macrophages. Our results showed that [^3^H]-radioactivity in liver, plasma, feces, and intestinal contents all showed significant increases in mice injected with DKO macrophages, but not those injected with WT macrophages (Figure 5D-5I and S8). These findings suggest that Epsin-deficient macrophages have higher cholesterol efflux and RCT.

**Figure 5.**
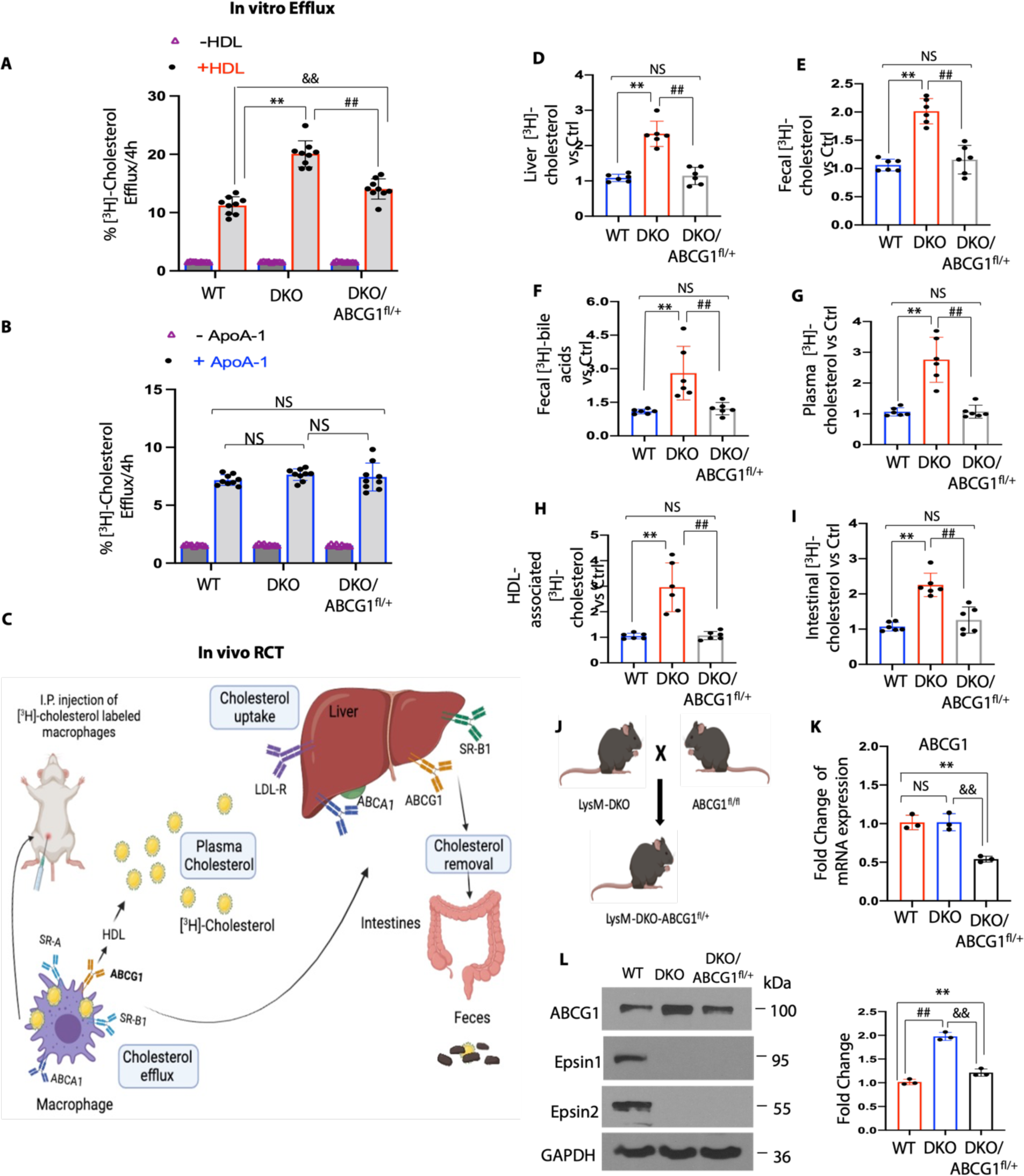
Epsins deficiency in macrophages shows increased cholesterol efflux *in vitro* and reverse cholesterol transport (RCT) *in vivo*. (**A-B**) Peritoneal macrophages were isolated from WT, DKO) and DKO/ABCG1^fl/+^ mice. *In vitro* [^3^H]-cholesterol labeled WT, DKO or DKO/ABCG1^fl/+^ macrophages were incubated in the presence or absence of HDL (25μg/mL) and ApoA-1 (10μg/mL) in the presence of 3μmol/L LXR agonist T0901317 (n=9). **WT vs DKO, P<0.01; ^##^DKO vs DKO/ABCG1^fl/+^, P<0.01; ^&&^WT vs DKO/ABCG1^fl/+^, P<0.01. (**C**) Schematic of [^3^H]-cholesterol loaded macrophage-RCT pathway. WT, DKO or DKO/ABCG1^fl/+^ macrophages were treated with 4μCi/mL [^3^H]-cholesterol and 50μg/mL ac-LDL for 48h followed by the injection of radiolabeled foam cells to C57BL/6/WT mice. After 48h, blood, liver and feces were collected and measured using a scintillation counter. (**D-I**) Distribution of [^3^H]-radioactivity counts in serum, HDL, liver feces and intestinal contents were determined by scintillation counter (n=6). **WT vs DKO, P<0.01; ^##^DKO vs DKO/ABCG1^fl/+^, P<0.01. (**J**) LysM-DKO mice were crossed with ABCG1^fl/fl^ mice to generate LysM-DKO-ABCG1^fl/+^ mice. (**K-L**) Peritoneal macrophages isolated from WT, LysM-DKO and LysM-DKO/ABCG1^fl/+^ mice (n=6) were analyzed by qRT- PCR (K) and WB (L) (n≥3). **WT vs DKO/ABCG1^fl/+^ group, P<0.01; ^##^WT vs DKO group, P<0.01; ^&&^DKO vs DKO/ABCG1^fl/+^ group, P<0.01. Data from A-L are presented as mean ± SD and were analyzed using one-way ANOVA (multiple comparisons).

### Epsin-UIM domain binds ABCG1 to promote degradation via the lysosome

To investigate the molecular mechanisms underlying Epsin-mediated ABCG1 regulation, we tested whether Epsins interact with ABCG1 and governed its turnover in macrophages. We first assessed protein levels of surface receptors such as ABCG1, ABCA1, SR-A1, and LDLR in macrophages involved in the progression of atherosclerosis. WB results showed that ABCG1, but not ABCA1, SR-A1 and LDLR, were upregulated in primary macrophages from DKO mice when compared to WT mice (Figure 6A). The mRNA levels of ABCG1 in both WT and DKO macrophages remain unchanged (Figure S9). Because ABCG1 is known to mediate cholesterol efflux to HDL ^48^, we hypothesized that Epsins bind ABCG1 under atherosclerotic conditions and promote ABCG1 ubiquitin-dependent degradation to inhibit cholesterol efflux in macrophages and facilitate the development of atherosclerosis. When the isolated peritoneal macrophages from DKO and WT mice were pretreated with a LXR activator followed by a cell surface biotinylation assay, we observed significantly elevated levels of biotinylated ABCG1 in DKO macrophages (Figure 6B). We also performed flow cytometry and observed that only WT macrophages showed a decreased surface level of ABCG1 with the oxLDL treatment in a clathrin-dependent way (Figure 6C and S10A-C). Therefore, decreased total and surface levels of ABCG1 in WT macrophages implies that Epsins promote ABCG1 degradation.

**Figure 6.**
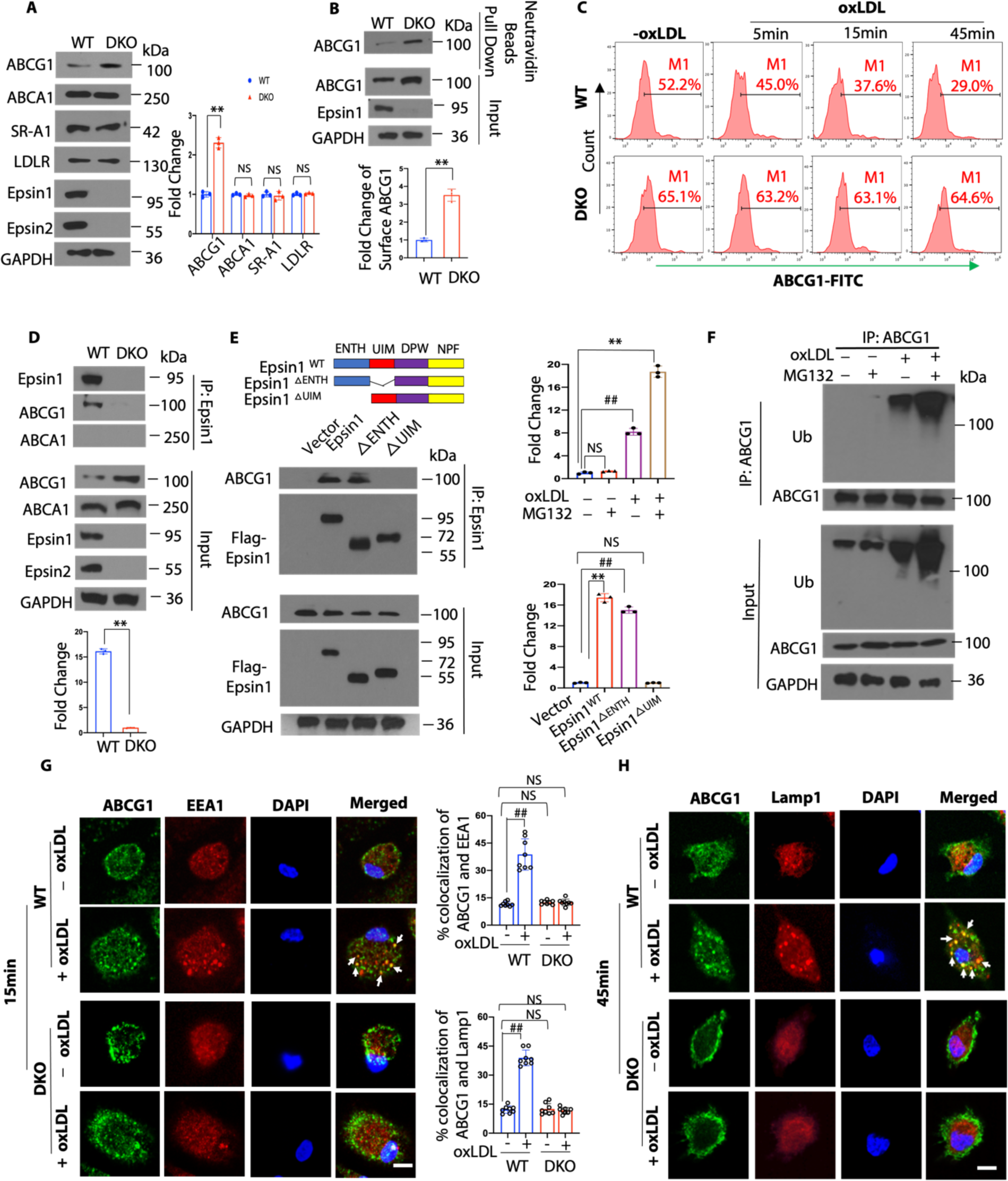
Epsins bind to ABCG1 and facilitate the internalization and degradation of ABCG1 via lysosomes. (**A**) Macrophages from WT (n=3) and DKO (n=3) mice were lysed for WB analysis (**WT vs DKO, P<0.01). (**B**) Macrophages from WT (n=3) and DKO (n=3) mice were pretreated with a liver X receptor (LXR) activator and followed by a cell surface biotinylation assay to evaluate the cell surface ABCG1 levels (**WT vs DKO, P<0.01). (**C**) LXR activated WT and DKO macrophages were incubated in lipid-deficient medium for 24h and treated with or without 100μg/mL oxLDL for 5, 15 and 45 min followed by staining with anti-ABCG1 and analyzed by flow cytometry. (**D**) LXR pretreated macrophages isolated from WT and DKO mice were treated with 100μg/mL oxLDL for 1h followed by IP and WB for Epsin1 and ABCG1 or ABCA1 (n=3, **WT vs DKO, P<0.01). (**E**) ABCG1 plasmids and full length (FLAG-Epsin1^WT^) or domain- deletion constructs (FLAG-Epsin1^△ENTH^ or FLAG-Epsin1^△UIM^) in the pcDNA3 vector were transfected into HEK 293T cells for 24 h in the presence of LXR agonist. Cells were then treated with 100μg/mL oxLDL for 1h, followed by IP and WB analysis using antibodies against FLAG tags and ABCG1 (n=3, **Epsin1 ^WT^ vs vector group, P<0.01; ^##^ Epsin1^△ENTH^ vs vector group, P<0.01). (**F**) Peritoneal macrophages isolated from WT mice were cultured in serum-free media for 24h followed by treatment with 5 μM MG132 for 3h. Cells were then treated with 100μg/mL oxLDL for 1h followed by IP and WB for ubiquitin and ABCG1. (**G-H**) WT and DKO macrophages were incubated in lipid-deficient medium for 24h followed by incubation with or without 100μg/mL oxLDL for 15min (G) or 45min (H) at 37^0^C. Macrophages were stained with ABCG1 (green), early endosome marker EEA1 (red) or lysosome marker Lamp1 (red) and DAPI (blue), then assessed by confocal microscopy. White arrows indicate the endocytic vesicles, scale bar=5μm, ^##^-oxLDL vs +oxLDL group (n=8), P<0.01. Data from A-H are presented as mean ± SD and were analyzed using either an unpaired Student’s t-test (two groups) or one-way ANOVA (multiple comparisons).

We next determined how Epsins cause increases in total and surface protein levels of ABCG1 in macrophages from DKO mice. When peritoneal- or bone marrow-derived macrophages from WT and DKO mice were lysed and processed for IP with anti-Epsin1 antibodies, we observed ABCG1 in WT macrophages, but not in DKO macrophages (Figure 6D). This suggested that Epsins bind to ABCG1. Accordingly, we showed that Epsin interacted with ABCG1 through the Epsin UIM domain, which was confirmed by transfection of Flag-tagged mammalian Epsin wild type full length, ENTH or UIM deletion constructs as well as ABCG1 plasmid to HEK 293T cells (Figure 6E). We showed that ABCG1 is ubiquitinated in response to oxLDL, allowing Epsins to interact with ABCG1 through their UIM, which promotes ABCG1 lysosomal degradation (Figure 6F). In addition, we observed oxLDL-induced ABCG1 trafficking to early endosomes (Figure 6G) and lysosomes (Figure 6H) in WT macrophages, but not DKO cells.

To confirm that elevated cholesterol efflux and RCT were a result of increased total and surface levels of ABCG1 in ApoE^-/-^/LysM-DKO macrophages, we created ApoE^-/-^/LysM-DKO-ABCG1^fl/+^ mice by crossing the ApoE^-/-^/LysM-DKO mouse strain with ABCG1 (flox/+) heterozygous mice to generate a novel mouse model (ApoE^-/-^/LysM-DKO/ABCG1^myeloid-het^) with genetically-reduced ABCG1 levels in Epsin-deficient macrophages (Figure 5J). We observed reduced gene (Figure 5K) and protein (Figure 5L) expression of ABCG1 in ApoE^-/-^/LysM-DKO-ABCG1^fl/+^ macrophages. The LysM-DKO-ABCG1^fl/+^ macrophages showed less cholesterol efflux to HDL and RCT compared to ApoE^-/-^/LysM-DKO macrophages (Figure 5A, 5B and 5D-5I). These studies demonstrated that Epsin deficiency in macrophages specifically enhanced ABCG1-mediated cholesterol efflux to HDL and RCT, which is inversely related to the risk of atherosclerotic cardiovascular disease^34^. In accordance, immunostaining of human patient aortic arch (Figure S11A) and mouse aortic root (Figure S11B) sections revealed that ABCG1 expression is inversely associated with atherosclerotic plaque severity, *i.e.,* there is considerable reduction in ABCG1 expression in more severe atheroma compared to that in less severe atherosclerotic plaques in both human and mouse atheromas.

### S2PNP-siEpsin1/2 hinders progression of early and advanced stages of atherosclerosis

Since we found that Epsins facilitate CD36-mediated lipid uptake and promote ABCG1 degradation to inhibit ABCG1-mediated cholesterol efflux, we sought to determine whether targeting Epsins could prevent the progression of atherosclerosis. We used a targeted NP platform^54^ modified with a lesional macrophage-specific targeting peptide (S2P) for systemic delivery of Epsins 1 and 2 siRNA combinations (S2PNP-siEpsin1/2) (Figure S12A).

Isolated macrophages from WT mice were treated with S2PNP-siControl (S2PNP-siCtrl) or S2PNP-siEpsin1/2. Then, RNA and protein were isolated to check the *in vitro* silencing efficacy of the NPs (Figure S12B and S12C). To assess the effect of NPs on *in vitro* peritoneal macrophage lipid loading, cells isolated from ApoE^-/-^ mice were treated with S2PNP-siCtrl or S2PNP- siEpsin1/2 for 24h followed by treatment with oxLDL or plasma isolated from ApoE^-/-^ mice fed a WD for 8 weeks. The treatment of oxLDL or plasma demonstrated decreased lipid uptake by S2PNP-siEpsin1/2-treated macrophages compared to S2PNP-siCtrl treated cells (Figure S13A and S13B).

ApoE^-/-^ mice fed a WD for 8 weeks (early stage, Figure S14) or 17 weeks (advanced stage, Figure 7) were divided into three groups including baseline (before NP injection), S2PNP-siCtrl, and S2PNP-siEpsin1/2. The baseline group was sacrificed before the injection of NPs.

**Figure 7.**
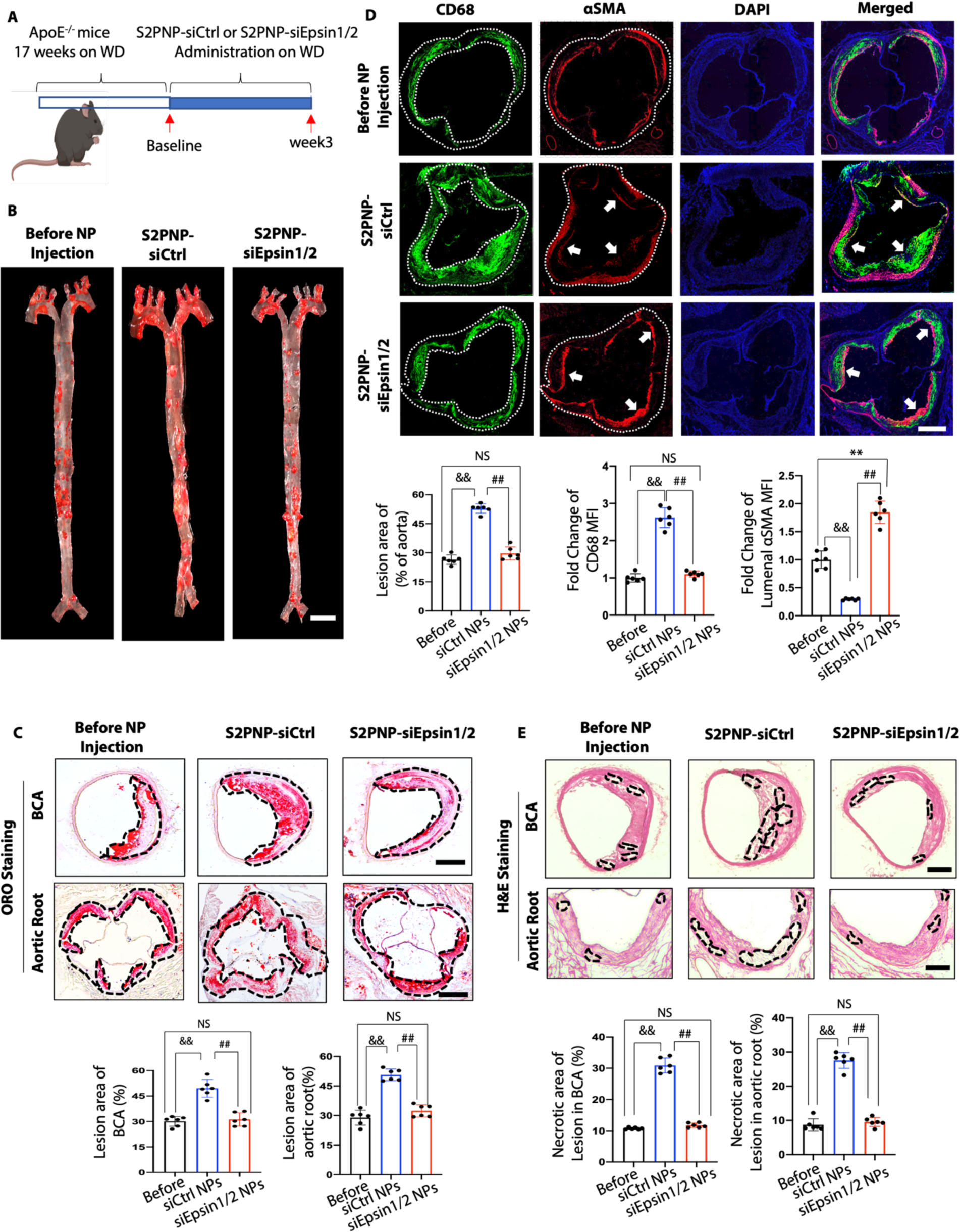
S2PNP-siEpsin1/2 delivery inhibits atheroma progression, decreases necrotic core content, and increases smooth muscle cell content in advanced stage of atherosclerosis. (**A**) ApoE^-/-^ mice fed a Western Diet (WD) for 17 weeks followed by treatment of S2PNP-siCtrl or S2PNP-siEpsin1/2 for 3 weeks (2 doses per week). (**B**) *En face* ORO staining of aortas from baseline, control siRNA NP-treated ApoE^-/-^ or Epsin1/2 siRNA NP treated ApoE^-/-^ mice fed a WD. lesion areas were analyzed with NIH ImageJ, scale bar=5mm. (**C**) ORO staining of brachiocephalic artery (BCA) and aortic root sections of above three groups. Lesional area (black dash line outlined) were analyzed by NIH ImageJ. (**D**) Aortic roots from S2PNP-siCtrl-treated ApoE^-/-^ or S2PNP-siEpsin1/2 treated ApoE^-/-^ mice were stained with the macrophage marker CD68 (dashed white line outlined) and αSMA (white arrow). (**E**) H&E staining of BCA and aortic root sections of the above three groups. Necrotic areas (black dash line outlined) were analyzed by NIH ImageJ. Data from B-E (n=6) are presented as mean ± SD and were analyzed using one-way ANOVA. ^&&^S2PNP-siCtrl vs baseline, P<0.01; ^##^S2PNP-siCtrl vs S2PNP-siEpsin1/2 group, P<0.01, scale bar: B=5mm; C, D, E=500μm.

S2PNP-siCtrl or S2PNP-siEpsin1/2 was intravenously injected twice a week for 3 weeks (Figure S14A and 7A). Digested lesions from S2PNP-siCtrl or S2PNP-siEpsin1/2 treated mice were used for WB to check *in vivo* silencing of Epsins 1 and 2 (Figure S14B). Additionally, the administration of S2PNP-siEpsin1/2 resulted in not only a decrease in lesional macrophage accumulation, but also a significant reduction of Epsins 1 and 2 expressions within the atherosclerotic plaques as showed by diminished CD68 and Epsins 1 and 2 staining in the aortic root sections (Figure S14C and S14D). *En face* ORO staining in aortas, aortic roots, and brachiocephalic artery (BCA) sections (Figure S14E, 7B and 7C) showed reduced lesion size, lesion number, and retarded progression of both early (11 weeks on WD) and advanced (20 weeks on WD) stages atherosclerosis in ApoE^-/-^ mice treated with S2PNP-siEpsin1/2 opposed to the S2PNP-siCtrl group.

Inhibiting the transition of atheroma from stable lesion to vulnerable plaque is paramount in preventing heart attacks and strokes. We observed elevated smooth muscle cell contents and reduced macrophage accumulation (Figure 7D), increased elastic fibers and collagen content (Figure S15), and reduced necrotic core area (Figure 7E) within the atherosclerotic lesions in S2PNP-siEpsin1/2 treated mice compared to S2PNP-siCtrl group. In addition, we observed no changes in plasma total cholesterol, triglycerides, or HDL and non-HDL cholesterol levels between S2PNP-siCtrl and S2PNP-siEpsin1/2 treated mice (Figure S16). These features suggest that the administration of lesion macrophage-specific targeting NPs containing Epsin1/2 siRNA can stabilize the atherosclerotic plaques without affecting the plasma cholesterol levels.

### S2PNP-siEpsin1/2 promotes atheroma resolution by diminishing inflammation and enhancing lesion stability

Given the critical role of macrophage cholesterol efflux and RCT in atheroma resolution, we also wanted to assess the therapeutic effects of S2PNP-siEpsin1/2 on atheroma resolution. We established another mouse model by injecting proprotein convertase subtilisin kexin type 9-adeno- associated virus-8 (AAV8-PCSK9D377Y) into C57BL/6 mice. This generated an atherosclerosis mouse model harboring a gain-of-function PCSK9 mutant ^61^. This model mimics LDLR null mice^26^ and requires no genetic modification ^61^. Another advantage of this mouse line is that a single AAV injection is sufficient to generate atherosclerotic mice by WD feeding compared to long-term crossbreeding with ApoE^-/-^ or LDLR^-/-^ mice.

To test if S2PNP-siEpsin1/2 treatment can promote atheroma resolution, PCSK9-AAV8 mice fed a WD for 16 weeks followed by 4 weeks of a normal diet with S2PNP-siCtrl or S2PNP- siEpsin1/2 treatment (Figure 8A). The latter group showed a significant reduction of lesion size and rapid lesion regression as evidenced in the IF and ORO staining of the whole aorta, aortic root and BCA sections (Figure 8B, 8C, 8D and S17A). Van Gieson’s trichrome staining of BCA sections showed increased collagen content and elastic fibers in the S2PNP-siEpsin1/2 group compared to S2PNP-siCtrl or baseline group (Figure S17B). In addition, the IF co-staining of CD68 and α-SMA showed reduced macrophage accumulation in the fibrous cap, reduced size of atheromatous cores, and increased smooth muscle cell content within the atheroma cap in the S2PNP-siEpsin1/2 group (Figure 8C).

**Figure 8.**
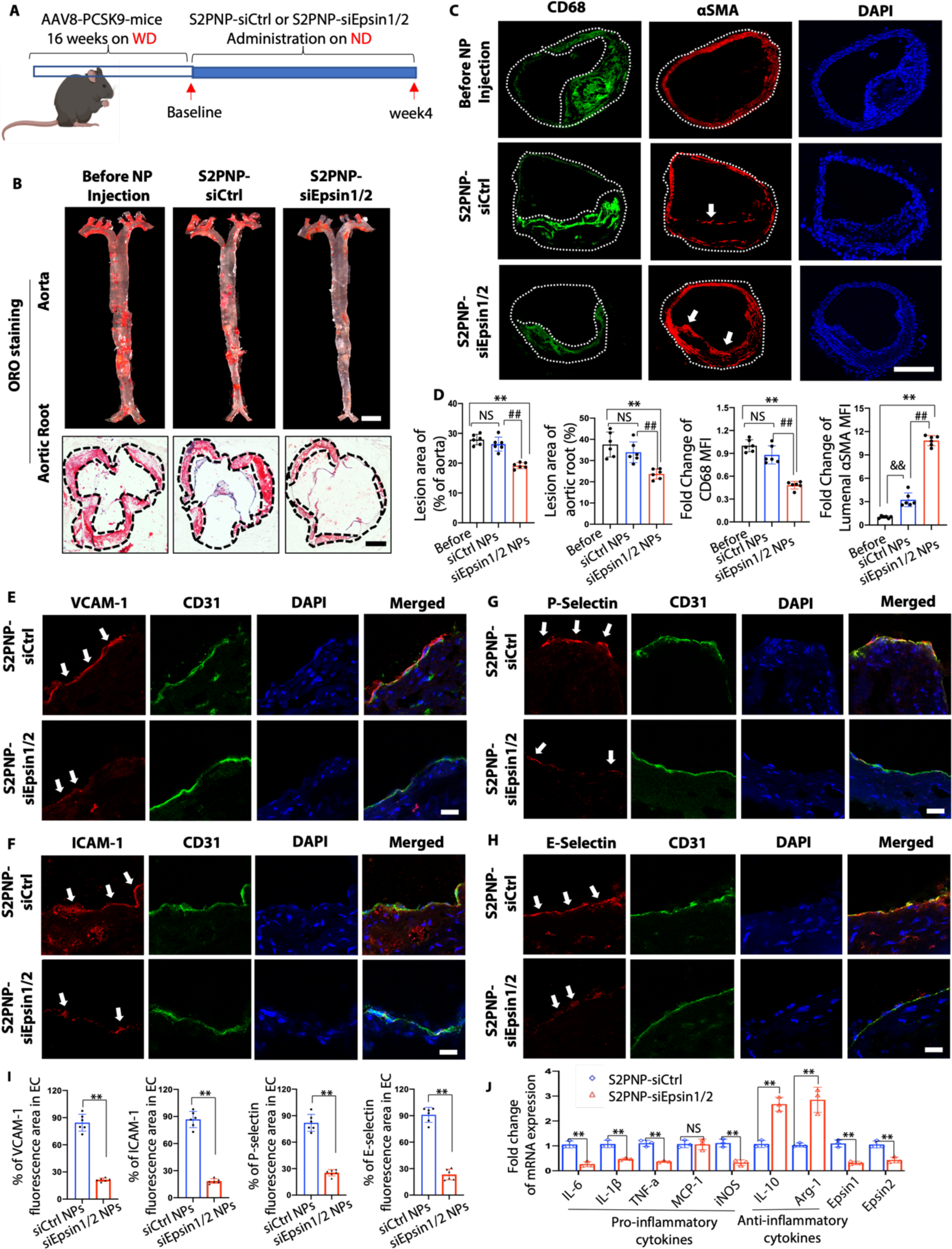
S2PNP-siEpsin1/2 promotes the resolution of atheroma, reduces inflammation, and stabilizes atherosclerotic plaques. (**A**) C57BL/6 WT mice were injected twice with PCSK9- AAV8 (D377Y) virus and fed a WD for 16 weeks and followed by normal diet feeding with the treatment of S2PNP-siCtrl or S2PNP-siEpsin1/2 for 4 weeks (2 doses per week). (**B**) *En face* ORO staining of aortas from above mice. Lesion area was analyzed using NIH ImageJ, scale bar=5mm (**C**) Immunostaining of CD68 (green) and αSMA (red) in the above groups. White arrows indicate the luminal αSMA. For (B) and (C), n=6, scale bar: B (aorta)=5mm, B (aortic root)=500μm, C=500μm. (**D**) Statistics for ORO staining (B) and IF staining (C), **S2PNP-siEpsin1/2 vs baseline, P<0.01, ^##^S2PNP-siCtrl vs S2PNP-siEpsin1/2 group, P<0.01. (**E-H**) Co-staining of CD31 (endothelial cell marker, green) with vascular cell adhesion molecule-1 (VCAM-1, red, E), intercellular adhesion molecule-1 (ICAM-1, red, F), P-selectin (red, G) or E-selectin (red, H) and DAPI on the luminal surface of aortic root sections. White arrows indicate CD31/ICAM-1, CD31/VCAM-1, CD31/E-selectin or CD31/P-selectin colocalization. Scale bars=20μm. (**I**) Statistical analysis of E-H, n=6, **S2PNP-siCtrl vs S2PNP-siEpsin1/2 group, P<0.01. (**J**) Isolated macrophages from PCSK9-mice were treated with S2PNP-siCtrl or S2PNP-siEpsin1/2 for 24h, RNA was isolated, and qRT-PCR was performed (n=3, **S2PNP-siCtrl vs S2PNP-siEpsin1/2 group, P<0.01). B-J are presented as mean ± SD and were analyzed using one-way ANOVA (B and C) and the unpaired Student’s t-test (E-J).

Adhesion molecules play a crucial role in attaching monocytes to activated endothelial cells (EC) as well as recruiting monocytes to the sub-endothelial layer. Co-staining of CD31 with vascular cell adhesion molecule-1 (VCAM-1), intercellular adhesion molecule-1 (ICAM-1), P-selectin or E-selectin (Figure 8E-8I) was performed using aortic root sections. Administration of S2PNP-siEpsin1/2 significantly suppressed these adhesion molecules when compared to S2PNP- siCtrl group. In addition, the expression of the macrophage egressing gene CCR7 was significant increased, while the retention genes Sema3a and Netrin1 were decreased in DKO macrophages (Figure S17C). This is consistent with our RNA-seq analysis that showed the deficiency of macrophage Epsins facilitates immune cell migration. Furthermore, macrophages treated with S2PNP-siEpsin1/2 showed decreased expression of pro-inflammatory genes (iNOS, IL-6, IL-1β, and TNF-α) and increased expression of anti-inflammatory markers (IL-10 and Arg1) compared to the S2P-siCtrl group (Figure 8J). These decreased pro-inflammatory cytokines lead to reduced ICAM-1, VCAM-1, P- and E-selectins expression in ECs (Figure 8E-8H), hindering recruitment of additional macrophages to the lesion. Taken together, these results indicate that specific silencing of Epsins in lesional macrophages may offer therapeutic benefits in preventing or treating advanced atherosclerosis.

## DISCUSSION

In this study, we discovered that Epsins 1 and 2 govern cholesterol transport (Figure S18) in atherosclerotic macrophages by regulating CD36-mediated lipid uptake (Figure 1-3 and S1-S6) and ABCG1-dependent cholesterol efflux (Figure 4-6 and S7-S11). Importantly, we demonstrated that targeting lesional macrophage Epsins using a newly developed nanotechnology-based siRNA-mediated therapy halts inflammation, impedes early and late atherogenesis (Figure 7 and S12-S16), and accelerates atheroma resolution (Figure 8 and S17). By silencing lesional macrophage Epsins, we observed a reduction in necrotic core area in atherosclerotic lesions and marked attenuation in inflammation (Figure 7 and Figure 8). Together, our studies point toward a promising new strategy for the treatment of atherosclerosis.

The advent of next generation therapies targeting PCSK9, including monoclonal anti-PCSK9 antibodies and PCSK9 siRNA, is considered to be game-changing ^62–64^; however, these and other therapies largely ignore the rampant inflammation that occurs in atherosclerotic arteries. Arterial inflammation is a prime contributor to the transition from a stable to vulnerable atheroma ^7–10^. While anti-inflammatory medicines such as anti-Interleukin-1β antibodies potently diminish the risk of recurrent myocardial infarction, these treatments may come with some severe side effects ^65, 66^. In addition, the inability to reduce lipid ingestion while concomitantly reinforce cholesterol efflux in lesional macrophages has impeded the battle against atherosclerosis. Our mechanistic studies show that Epsins bind to CD36 to facilitate lipid uptake by enhancing CD36 endocytosis and recycling. Conversely, Epsins promote ABCG1 degradation via lysosomes and hamper ABCG1-mediated cholesterol efflux and reverse cholesterol transport. Excitingly, by harnessing the latest development in nanomedicine and developing S2P-NPs carrying Epsin1/2 siRNAs (S2PNP-siEpsin1/2), we show a reduction in plaque formation in both early and advanced atherosclerosis (Figure S14 and 7). Furthermore, in a pre-clinical atheroma resolution model, this potent approach reduced lesional macrophage accumulation, decreased pro-inflammatory cytokine and adhesion molecule expression, and accelerated plaque regression (Figure 8). Taken together, our study uncovers a novel and critical role for Epsin in regulating lipid metabolism and cholesterol efflux as well as offers a potential new nanotherapy for treating atherosclerosis by targeting Epsins in macrophages.

As we show that silencing of lesional macrophage Epsins reduces necrotic core size, we reasoned that inhibiting Epsins may reinforce efferocytosis in macrophages, analogous to our observations in DKO macrophages ^26^ given that efferocytosis plays a central role in regulating necrotic core formation ^67–69^. Despite the fact that S2P-NPs have been shown to only negligibly target the endothelium ^54^, whether S2PNP-siEpsin1/2 prohibits endothelial Epsins to synergistically curb atherosclerosis may require further investigation as deletion of endothelial Epsins also reduces inflammation and atherogenesis ^24, 25^. Interestingly, we did not observe significant changes of the lipid profiles in the plasma from S2PNP-siCtrl and S2PNP-siEpsin1/2 treated ApoE^-/-^ mice, suggesting either a minor liver targeting by S2PNP-siEpsin1/2 or that S2PNP- siEpsin1/2-mediated silencing of Epsins in liver could be inconsequential. Nevertheless, whether S2PNP-siEpsin1/2 targets liver epsins warrants further investigation.

While autophagy can eliminate excess lipid droplet-associated cholesterol esters taken up by macrophages ^70–73^, this process is attenuated during atheroma progression, leading to defective lipolysis, impaired efferocytosis, and escalated inflammation, which increases the likelihood of vulnerable lesion rupture ^70–76^. A recent report suggests that macrophage SR-B1 modulates autophagy through the VPS34 complex and PPARα transcription of Tfeb in atherosclerosis ^77^. SR- B1 is a multifunctional membrane receptor, which is not only responsible for selective uptake of HDL-derived cholesteryl esters into cells, but also for transferring HDL-cholesterol efflux from peripheral tissues to back to liver ^78^. Given that Epsins are implicated in autophagy function in *Drosophila* ^79^, the question of whether these proteins suppress autophagic activity by binding to SR-B1 and inhibiting SR-B1-regulated function in macrophages remains unanswered.

Our studies also revealed that LXR-activated macrophage surface expression of ABCG1 was significantly reduced with oxLDL treatment in WT, but not in DKO macrophages owing to their internalization and degradation mediated by the Epsins (Figure 6). It is thought that basal macrophage ABCG1 is largely located at an intracellular location ^80^ and then redistributed to the cell surface following LXR activation ^81^. Our immunostaining and biotinylation studies (Figure 6) are consistent with these observations. More importantly, staining of ABCG1, in both human patient aortic arch and mouse aortic root sections, exhibits a reduction in ABCG1 expression in severe atherosclerotic lesions (Figure S11), suggesting a pivotal role of loss of Epsins in maintaining the abundance of ABCG1 and limiting plaque buildup. Recent clinical trials suggest that enhanced cholesterol efflux capacity (CEC) considerably reduces the risk of repeat heart attacks and severe acute myocardial infarction (MI) ^39–41^. In addition, cholesterol efflux capacity has minimal links to various risk factors and metabolic issues in contrast to HDL cholesterol level^45^. Our results show that targeting Epsins augments cholesterol efflux (Figure 5 and S16) and consequently, may potentially enhance CEC, suggesting that inhibiting Epsins represents a promising new strategy for treating patients suffering from recurrent MI and severe CAD.

In conclusion, our results provide a powerful proof-of-concept for the therapeutic potential of Epsin siRNA-containing NPs to precisely target lesional macrophages to abolish inflammation and resolve atheroma. By illustrating that targeting Epsins expression in macrophages restricts lipid ingestion in foam cells and concomitantly enhances cholesterol efflux, we believe this study provides a foundation for the renewed quest seeking innovative therapies that precisely resolve lesional inflammation in advanced atherosclerosis.

## Nonstandard Abbreviations and Acronyms

AAV8: adeno-associated virus-8
ABCG1: ATP Binding Cassette Subfamily G Member 1
ACAT: acyl-CoA cholesterol acyltransferase
AMI: acute myocardial infarction
BCA: brachiocephalic artery
CAD: coronary artery disease
CEC: cholesterol efflux capacity
DKO: double knockout
ENTH: epsin N-terminal homology
GEM: gel beads-in-emulsion
GO: gene ontology
GSEA: gene set enrichment analysis
HDL: high density lipoprotein
ICAM-1: intercellular adhesion molecule-1
IF: immunofluorescence staining
IP: immunoprecipitation
LXR: liver X receptor
M-CSF: macrophage colony stimulating factor
NP: nanoparticle
ORO: oil red O
oxLDL: oxidized low-density lipoprotein
PCSK9: proprotein convertase subtilisin/kexin type 9 serine protease
RCT: reverse cholesterol transport
scRNA-seq: single-cell RNA sequencing
siRNA: small interfering RNA
SR-A: scavenger receptor-A
S2PNP-siEpsin1/2: S2P-conjugated Epsins 1 and 2 siRNA nanoparticle S2PNP-siCtrl S2P-conjugated Control nanoparticle
TG: thioglycolate
UIM: ubiquitin-interacting motif
VCAM-1: vascular cell adhesion molecule-1
VSMCs: vascular smooth muscle cells
WB: western blot
WD: western diet

## ACKNOWLEDGMENTS

We thank researchers for providing their GEO database information online. We thank the Flow Cytometry Core at Boston Children’s Hospital for the use of the LSRII, the Viral Core at Boston Children’s Hospital for providing viral stocks, and the Biopolymers Facility at Harvard Medical School for quality control analysis of RNA samples.

## AUTHOR CONTRIBUTIONS

K.C. J.S., K.F.C. and H.C. conceived and designed the study. K.C. primarily contributed to the identification of the in vitro molecular mechanism. K. C. and B.W. primarily contributed to the in vivo data in atherosclerosis analysis. X.G., K.C. R.Z., and K.F.C. conducted the RNA-seq data analysis and interpretation. H.W. and Y. D worked on part of the molecular mechanism investigation and provided biochemistry insights. Y.X. and X.J. synthesized, characterized the nanoparticles. M.M. performed the cholesterol and triglyceride analysis. Y.L. and B.Z. provided technical support in RNA-seq experiment. K.C., D.S. and Q.P. analyzed the data and provided comments. S.W. provided technical support with mouse genotyping and colony maintenance. K.C. and H.C. wrote the manuscript. K.C., D.B.C., K.L., S.S., M.F.L., J.S., K.F.C., and H.C. edited the manuscript. All the authors reviewed and provided feedback on the manuscript.

## Online Supplemental Materials

### The supplemental materials include

Data file S1. Differentially expressed genes between cell clusters identified in all the cells in aorta.

Data file S2. Differentially expressed genes between macrophage subclusters. Data file S3. Differentially expressed genes in DKO compared to wild type.

Data file S4. Gene Ontology enrichment analysis for DEGs shared by DKO and CD36KO.

Data file S5. Gene differential expression analysis between ABCG1KO and wild type macrophages.

Data file S6. Gene Ontology enrichment analysis for DEGs in ABCG1KO compared to wild type. Data file S7. Gene Ontology enrichment analysis for pathways reversely regulated by ABCG1KO and DKO.

Data file S8. Uncropped Western Blot.

Data file S9. Primary data of statistical analysis.

## MATERIALS and METHODS

### Isolation of primary mouse macrophage and macrophage culture

The isolation of peritoneal macrophages was performed as described previously ^82^. Briefly, mice were intraperitoneally injected with 1mL of 4% thioglycolate (TG), and 3 days post-injection, mice were sacrificed, and peritoneal cells were harvested with 7mL of sterile PBS by lavage of peritoneal cavity. Cells were spun down (1000xg, 5 minutes), washed with PBS, resuspended, and plated in RPMI (containing 10% FBS and 1% Pen-Strep) at 37 °C in humidified air containing 5% CO_2_ atmosphere. After 3 hours, non-macrophages were washed with PBS. For bone marrow- derived macrophages, mice were sacrificed, and both femurs and tibias were dissected and flushed with sterile 1X PBS, followed by passing through a 70μM cell strainer. Cells were spun down (1000xg, 5 minutes), washed with PBS, and seeded in RPMI (containing 10% FBS and 1% Pen- Strep) with macrophage colony stimulating factor (M-CSF, 10ng/mL) to differentiate into macrophages for 5 days. After which, the macrophages were harvested and used in experiments. Both bone marrow-derived and isolated peritoneal primary macrophages were used to confirm knock out of Epsin1 and Epsin2. Isolated peritoneal macrophages were mainly used for western blots, flow cytometry and immunoprecipitations due to higher yields.

### RNA isolation, quantitative real-time PCR and RNA sequencing

Total RNA was extracted from primary macrophages with Qiagen RNeasy Mini Kit based on manufacturer’s instruction including the optional step to eliminate genomic DNA. The extracted RNA was either used for qRT-PCR or RNA sequencing according to the experimental designs.

For qRT-PCR, mRNA was reverse transcribed to cDNA with the iScript cDNA Synthesis Kit (Bio-Rad Laboratories, Inc., Hercules, CA, United States). 2 μL of the product was subjected to qRT-PCR in StepOnePlus Real-Time PCR System (Applied Biosystems) using SYBR Green PCR Master Mix reagent as the detector. PCR amplification was performed in triplicate on 96-well optical reaction plates and replicated in at least three independent experiments. The ΔΔCt method was used to analyze qPCR data. The Ct of β-actin cDNA was used to normalize all samples. Primers are listed in Supplemental Table S1. For RNA sequencing, extracted RNA from primary macrophages with Qiagen RNeasy Mini Kit based on manufacturer’s instruction were sent to BGI Genomics Company (San Jose, USA) for RNA sequencing.

### RNA sequencing data processing and differential expression analysis

The raw reads of RNA sequencing data were mapped to the mouse genome (version mm10) using STAR (version 2.7.9a) ^83^ or TopHat (version v2.1.1) ^84^. The read count was calculated for each gene by htseq-count (version 0.11.2) ^85^ and further normalized to TPM (transcripts per million) and FPKM (fragments per kilobase of transcript per million fragments mapped). For differential expression analysis of Epsin deficient and wild type macrophage dataset, the read count matrix produced by htseq-count was imported into the R package DESeq2 (version 1.30.1)^86^. For the ABCG1 knock out dataset, we first performed a batch effect removal on the log10 transformed TPM expression matrix using ComBat in the R package sva (version 3.38.0) ^87^. Then, the R package limma was used to do differential expression analysis for the batch effect-removed expression matrix. Genes with fold change larger than 1.2-fold and p-value less than 0.05 were identified as differentially expressed genes (DEGs). Principal component analysis (PCA) was conducted by the plotPCA function of DESeq2. Heatmap of gene expression values was generated using the R package heatmap (version 1.0.12). Gene Ontology (GO) functional enrichment analysis was performed by the R package clusterProfiler (version 3.18.1) ^88^.

For the comparison of DEGs between DKO and CD36 knockout macrophages in mice, we utilized the published bulk RNA-seq data from Chen et. al. ^59^, with the accession number GSE139439, downloaded from NCBI GEO database ^89^. The overlapping DEGs co-regulated in DKO and CD36 knockout macrophages were analyzed by the R package VennDiagram (version 1.6.20) ^90^. The statistical significance of overlapping genes was calculated by Fisher exact test. All the data for bulk RNA-seq are available in Data files (S3-S7).

### Cell culture and plasmids transfection

The HEK 293T cell line (ATCC no. CRL-11268) were used for plasmid transfection to map the binding sites of Epsin to CD36 or ABCG1. Flag-tagged Epsin1^WT^, Epsin1^ΔUIM^, Epsin1^ΔENTH^ truncation constructs, and pcDNA vector were prepared previously in our lab^25^. CD36 (lot:52025) and ABCG1 (lot:53952) plasmids were purchased from AddGene. HEK 293T cells were cultured in DMEM (10% FBS and 1% Pen-Strep) at 37°C in humidified air containing 5% CO_2_ atmosphere and transfected using Lipofectamine 2000 as instructed by the manufacturer. Transfection of Epsin domains to macrophage: Isolated WT and DKO peritoneal macrophages were cultured in RPMI media (containing 10% lipid-depleted serum and 1% Pen-Strep). Epsin1^WT^, Epsin1^ΔUIM^, Epsin1^ΔENTH^ truncation constructs, and pcDNA vector were transfected to macrophages using lipofectamine LTX transfection reagent or using Nucleofector II apparatus (Amaxa, Germany) with mouse macrophage nucleofector kit (Lot: VPA-1009, Lonza) as instructed by the manufacturer.

### Immunoprecipitation (IP) and western blotting (WB)

For total protein levels, primary macrophages were washed with ice cold PBS, lysed in RIPA buffer (50 mM Tris HCl, 150 mM NaCl, 1.0% NP-40, 0.5% Sodium Deoxycholate, 1.0 mM EDTA, 0.1% SDS and 0.01% sodium azide at a pH of 7.4.), added 4X Laemmli buffer (1:3 dilution in lysis buffer) was added and WB was perfomed for the proteins indicated in this study. For IP, cells were washed with ice cold PBS, lysed with lysis buffer (1% Triton X-100, 5mM Na_3_VO_4_, 10mM N-ethylmaleimide, and protease inhibitor cocktail), spun down (12000xg, 5 min at 4°C) to remove the debris. Cell lysates were pre-cleared with appropriate species of IgG and protein A/G Sepharose beads for 1h at 4°C with rotation followed by incubation with A/G Sepharose beads and indicated antibodies for 12 hours at 4°C with rotation. For negative controls, equal concentrations of mouse IgG were added instead of specific antibodies. Precipitated proteins were washed with ice cold lysis buffer for 3 time and eluted from protein A/G beads using 2X Laemmli buffer (1:1 in lysis buffer) followed by WB as described previously ^26^. WB were repeated for at least 3 times with different mice and bands were quantified using NIH ImageJ software. The protein expression levels were normalized to GAPDH levels. For IPs in macrophages involving oxLDL treatment, cells were pre-treated with 1 μM MG132 in serum-free media for 4 hours, followed by the treatment with or without oxLDL (100μg/ml) for 30 minutes at 37°C and processed for immunoprecipitation. For the transfection of Epsin1 constructs and CD36 or ABCG1 constructs, transfected HEK 293T cells were cultured in lipid-depleted DMEM media for 24 hours followed by stimulation with 100μg/mL oxLDL for 30 minutes at 37°C and performed for immunoprecipitation.

### Atherosclerosis analysis

Mice were anesthetized with isoflurane. Blood was collected from the right atrium followed by left ventricle perfusion with cold PBS. Whole aortas, brachiocephalic artery (BCA) and hearts of the mice were isolated. Whole aortas were dissected symmetrically, pinned to parafilm and fixed in 4% PFA to allow the *en face* analysis. Heart and BCA were embedded in OCT mounting medium and immediately frozen. The aortic sinus and BCA in the heart were sectioned at 10 microns (at least 9 sections of each sample were collected). The internal elastic lamina and luminal boundary of the lesion was manually traced and the lesion sizes of the *en face* aortas and aortic roots were quantified by NIH ImageJ software. The methods for Oil Red O (ORO) staining, immunofluorescent (IF) staining, Hematoxylin and Eosin (H&E) staining, and Van Gieson’s staining are described in supplemental materials ^26^.

ORO staining and IF staining of primary macrophages or cryosections were performed as described below. Oil Red O imaging was taken by a Zeiss Axio Scope.A1, AxioCam ICc5, and analyzed by ZEN-Lite 2012 software. Imaging of *en face* aortas was performed using a Nikon SMZ1500 stereomicroscope, SPOT Insight 2Mp Firewire digital camera, and SPOT Software 5.1. Imaging of IF staining was taken by Zeiss confocal microscope and quantification areas were performed by manually tracing the aortas, BCA, and aortic root lesion areas with NIH ImageJ software. Statistical analysis of samples including Oil Red O, Van Gieson’s, H&E, and IF staining were performed by blinding in which each animal was assigned a number and data was collected based on the assigned number with genotype and experimental condition unknown to the data collector.

### Oil Red O staining

For cryostat sections: cryostat sections 10 microns were fixed in 4% paraformaldehyde. Slides were washed with PBS (3 times, 5 min each time), and rinsed with 100% propylene glycol followed by staining with freshly prepared 0.5% Oil Red O solution for 10 minutes at 60°C. Slides were then put in 85% propylene glycol for 2 min, followed by 3 washes in water. Slides were next incubated with hematoxylin for 1-2 min, rinsed 3 times in water, and mounted with aqueous mounting medium. For foam cells: coverslips were washed with PBS, fixed in 4% paraformaldehyde for 10 min and stained with freshly prepared 0.5% Oil Red O solution for 10 min at 65°C. Slides were then washed in PBS (3 times, 5min each time), incubated with hematoxylin for 1min, washed with PBS 3 times, and mounted on coverslips with aqueous mounting medium. Imaging was processed with a Zeiss LSM880 confocal microscope and analyzed with ZEN-Lite 2012 software and NIH ImageJ software. Quantification of lesion was performed as described above. Quantification of foam cells was performed as described below.

### Hematoxylin and Eosin staining

Frozen aortic root and BCA sections: slides were fixed in 10% buffered formalin for 15 min and washed in water. Next, slides were stained with 0.1% hematoxylin for 3min followed by ddH_2_O washes, 95% ethyl alcohol and water. Slides were then dipped in 0.5% Eosin for 3 min, quickly rinsed with ddH_2_O, dipped in 95% and 100% ethanol, incubated in 50:50 Xylenes:100% ethanol and incubated in 100% Xylenes. Slides were mounted using Permount with coverslips.

### Van Gieson’s staining

Sections were fixed in 10% buffered formalin for 15 min and washed in water. Slides were stained with hematoxylin for 10 min, washed in ddH_2_O, stained 1-3 min in Van Gieson’s solution, dehydrated in 95% alcohol and 100% alcohol two times. Then, slides were cleared in xylene for two times and mounted with Permount. Staining results were presented as: Elastic fibers and nuclei–Black, Collagen fibers–Red and Other tissue elements–Yellow. Lesion area was traced using NIH ImageJ software. The percentage of necrotic area was determined by necrotic areas within the lesion. Collagen content was determined by the percentage of lesion areas.

### Immunofluorescence staining

Human samples: human healthy and atherosclerotic aorta paraffin sections were deparaffinized in xylene for 15min, immersed in graded ethanol (100%, 100%, 95%, 90%, 80%, and 70%, each for 3 min), washed with running tap water and processed antigen retrieval with 10mM Sodium Citrate, pH 6.0, with 0.05% Tween 20 at 90°C for 10 min. Samples were blocked in PBS with 3% donkey serum, 3% BSA, and 0.3% Triton X-100 and incubated with primary antibodies ABCG1 or CD68 (1:70 dilution) at 4°C overnight. The sections were washed three times and respective secondary antibodies conjugated to fluorescent labels (Alexa Flour 594, 488, or 647; 1:200 to 1:500) were added for 2 h at room temperature. The sections were mounted with mounting medium containing DAPI (1:100).

Mouse aortic root and BCA cryosections: Sections were fixed by 4% paraformaldehyde for 30 min at room temperature and blocked in PBS solution containing 3% donkey and/or goat serum, 3% BSA, and 0.3% Triton X-100 for 1hour. Samples were then incubated with primary antibody at 4°C overnight, followed by incubation with the respective secondary antibodies conjugated to fluorescent labels (Alexa Flour 594, 488, or 647; 1:200 to 1:500) for 2 h at room temperature. The sections were mounted with mounting medium containing DAPI (1:100).

Staining of peritoneal macrophages: macrophages plated on the 18-mm coverslips were washed with PBS, fixed by 4% paraformaldehyde for 15 min at room temperature and blocked in PBS solution containing 3% donkey and/or goat serum, 3% BSA, and 0.3% Triton X-100 for 1hour. Coverslips were then incubated with primary antibody (CD36, ABCG1, EEA1, Rab11, or Lamp1; 1:100-1:300) at 4°C overnight, followed by incubation with the respective secondary antibodies conjugated to fluorescent labels (Alexa Flour 594, 488, or 647; 1:200 to 1:500) for 1 hour at room temperature. The sections were mounted with mounting medium containing DAPI (1:100). BODIPY^TM^ 493/503 staining of macrophages was performed following F4/80-fluorescent conjugated antibody incubation for 2 hours at room temperature ^91^. Slides were washed with PBS, stained with DAPI and mounted. Immunofluorescent images were captured using a Zeiss LSM880 confocal microscope and analyzed with ZEN-Lite 2012 software and HIH ImageJ software.

Samples stained without the primary antibody were obtained using the same settings as negative controls. Mean fluorescence intensity (MFI) of antibody staining was determined using NIH ImageJ software with n=3 or more.

### Flow cytometry assay

Flow cytometry of elicited primary macrophages: peritoneal macrophages from WT and DKO mice were isolated as described above and plated in 6 well plates in lipid-deficient medium for 24 hours followed by the treatment with or without 100μg/mL oxLDL in the presence or absence of clathrin siRNA at 37^0^C for different time based on the experiment designs. Macrophages were washed with 1XPBS, dissociated with 1mL non-enzymatic cell dissociation buffer, centrifuged (300x*g*, 5 minutes), and resuspended in 100μL FACS buffer (1X PBS, 2% FBS, 2mM EDTA) containing the following: FcR Blocking Reagent, fluorochrome conjugated anti-F4/80, primary antibodies against CD36 or ABCG1. Cells were incubated with the primary antibodies (1:100) on ice for 30min, washed with 100μL FACS buffer, spun down and resuspended with 100μL FACS buffer containing the fluorescent secondary antibodies (1:100). After 30 min, cells were washed with FACS buffer, fixed with 4% paraformaldehyde (PFA), and resuspended in FACS buffer for analysis. Single color and no color controls were prepared using elicited macrophages, which were treated the same as experimental groups. Expression of cell markers was analyzed using a FlowJo version 10 software. Gating strategies were performed as described in Supplemental Figures S4 and S10. Flow cytometry of DiI-oxLDL treated macrophages: peritoneal macrophages elicited from WT and DKO mice were incubated in lipid-deficient medium for 24h followed by the treatment of DiI-oxLDL for 2h at 37^0^C and macrophages were washed with 1XPBS, dissociated with 1mL non-enzymatic cell dissociation buffer, centrifuged (300xg, 5 minutes), resuspended in 100μL FACS buffer (1X PBS, 2% FBS, 2mM EDTA) containing the following: FcR Blocking

Reagent, fluorochrome conjugated anti-F4/80 antibody staining, fixed and assessed the uptake of lipoproteins by flow cytometry as described above.

### Foam cell formation

TG (4%) induced peritoneal macrophages were isolated and plated on 18mm glass coverslips. Cells were cultured in RPM media (containing 10% lipid-depleted serum and 1% PennStrep) for 24 hours and then treated with 10-100 μg/mL oxLDL for 24 hours ^26^. Cells were fixed in 4% PFA for 10 minutes at room temperature and washed with PBS. For Oil Red O staining, coverslips were stained with Oil Red O, washed with PBS, counterstained with hematoxylin, washed with PBS, and then mounted on slides. For Bodipy staining, coverslips were immunofluorescently stained BodipyTM 493/503 and phalloidin-iFluor 555 reagent for 1 hour at 37°C, counterstaining with DAPI, and mounting on slides. Negative controls were not treated with oxLDL. Foam cells were determined as the number of lipid positive cells (Oil Red O positive or Bodipy positive) as a percentage of total cells. At least 6 fields per cover slip and 6 mice per genotype were used for quantification.

### Cell surface biotinylation

Cell surface biotinylation was performed as described previously ^26^. Isolated peritoneal macrophages in the plate were washed with cold PBS, suspended at a concentration of 25×10^6^ cells/mL and treated with 2mM EZ-Link Sulfo-NHS-LC-Biotin reagent on ice for 30 minutes followed by 3 washes with 100mM Glycine to remove excess biotin and then 3 washes with cold PBS (5min each time). Cells were lysed and pulled down by streptavidin bead: cell lysates were incubated with neutravidin beads for at least 12 hours at 4°C with rotation, and proteins were eluted from beads using 4X Laemmli buffer diluted 1:3 in lysis buffer. Cell surface biotinylated proteins were analyzed by western blotting and quantified using NIH Image J software.

### Plasma collection, triglyceride, and cholesterol analysis

For each mouse, 1mL syringe were rinsed with 1mL 0.5M EDTA to coat the inside of the syringe with EDTA to prevent clotting during blood collection. Blood was collected from the right atrium of the mouse heart after sacrifice with isoflurane and added to each 1.7mL tube containing 50μL 0.5M EDTA. Blood was centrifuged at 2000xg for 10 minutes at 4°C. Plasma was transferred to a new tube and stored at -20°C. Plasma cholesterol and triglyceride levels were determined as described below.

Total cholesterol, high-density lipoprotein (HDL) cholesterol and triglycerides in mouse serum were determined on Ace Axcel® Clinical Chemistry System (Alfa Wassermann, West Caldwell, NJ). Non-HDL cholesterol is calculated as Total Cholesterol – HDL and gives a measure of the cholesterol carried by all of the atherogenic lipoproteins.

Quantification of total cholesterol, HDL and triglycerides in macrophages. Lipids were extracted from macrophages with hexane:isopropanol (3:2, v:v) as described in Robinet et al ^92^. Solvents were removed by drying with nitrogen gas. Extracts were resolubilized in 5% bovine serum solution by bath sonication (10 minutes at 37°C), freeze/thaw treatment (1 hour at -80°C) and followed by probe sonication for 10 seconds. Reconstituted samples were analyzed on Ace Axcel® Clinical Chemistry System for total cholesterol, HDL and triglycerides as described above for measurements in mouse serum. Obtained values were normalized to total protein measured in the cell lysates by Bradford Protein Assay (Bio-Rad).

### Macrophage cholesterol efflux assay

Thioglycolate induced peritoneal macrophages from WT, DKO and DKO/ABCG1^fl/+^ mice were plated in RPMI medium for 2-4 hours. Non-adherent cells were removed by washing with PBS and cells were incubated with radiolabeled medium supplemented with 4 μCi/mL of [^3^H]- cholesterol (Perkin-Elmer, Waltham, MA, USA), 5% FBS,1% P/S and 50μg/mL acetyl-LDL for 24 hours. Macrophages were washed twice with warm PBS and incubated with serum-free RPMI 1640 medium supplemented with 0.2% BSA, 2μg/mL acyl-CoA cholesterol acyltransferase (ACAT) inhibitor and 4μmol/L LXR agonist T0901317 for 18 hours equilibration ^81^. After this equilibration period, cells were washed twice with warm PBS and incubated in serum free RPMI medium supplemented with or without cholesterol acceptors (10 μg/mL ApoA-1 or 25 μg/mL HDL) and 2μg/mL ACAT inhibitor for 4 hours ^93^. At the end of the incubation, the efflux media was collected and filtered through a 0.45-μm filter to remove the detached cells. Then, transfer the efflux medium was transferred to a scintillation vial. The macrophages in the plates were added to 500 μL of 0.2N NaOH and incubated on shaker at 4 °C overnight. Cell extract from each well was transferred to a scintillation vial. 4 mL of scintillation liquid were added to each scintillation vial and radioactivity was measured by liquid-scintillation counting ^94^. The cholesterol efflux to acceptors was expressed as a percentage of total cholesterol using the following formula: % cholesterol efflux = (medium [^3^H]-radioactivity [cpm]) / [(medium [^3^H]-radioactivity [cpm] + [^3^H]-radioactivity from cell extract [cpm])] x 100, where cpm = counts per minute.

### Reverse cholesterol transport (RCT) assay

In vivo RCT experiment is based on the method detailed by Joan Carles Escolà-Gil et al ^94^. Briefly, macrophages were radiolabeled with [^3^H]-cholesterol (5 μCi/mL) in 10% lipoprotein-depleted serum, 1% P/S and 50μg/mL acetyl-LDL RPMI media for 48h. Foam cells were washed with serum-free media supplemented with 0.2% BSA and equilibrated for 4h (37 °C, 5 % CO_2_). Then, [^3^H]-cholesterol-labeled macrophages were detached, spun down and resuspended in PBS. The injection dose was 0.5 mL per mouse (4×10^6^ cpm/mL) administered intraperitoneally into C57BL/6 WT mice fed on normal diet as indicated in Figure 5C. Mice were then individually housed in metabolic cages and feces in the cage floor were collected for 2 days. At 48h, mice were sacrificed and blood, liver, intestinal contents were collected in Figure S8. Serum [^3^H]-cholesterol was measured by liquid scintillation counting. [^3^H]-HDL cholesterol was determined after precipitation of ApoB-containing lipoproteins with 0.44mM phosphotungstic acid and 20mM MgCl_2_. Liver, fecal and intestinal lipids were extracted with hexane-isopropanol (3:2, v:v) and partitioned against Na_2_SO_4_. The lipid layer was collected and dried for 48h using nitrogen gas in a fume hood, and [^3^H]-cholesterol radioactivity was measured by liquid scintillation counting (4mL scintillation fluid for 4 min). The [^3^H]-radioactivity observed in fecal biliary acids was determined in the remaining aqueous phase of fecal material extracts. The amount of [^3^H]-radioactivity was expressed as a percentage of the total injected dose, which was taken as 100%.

## SUPPLEMENTAL FIGURES AND LEGENDS

**Figure S1.**
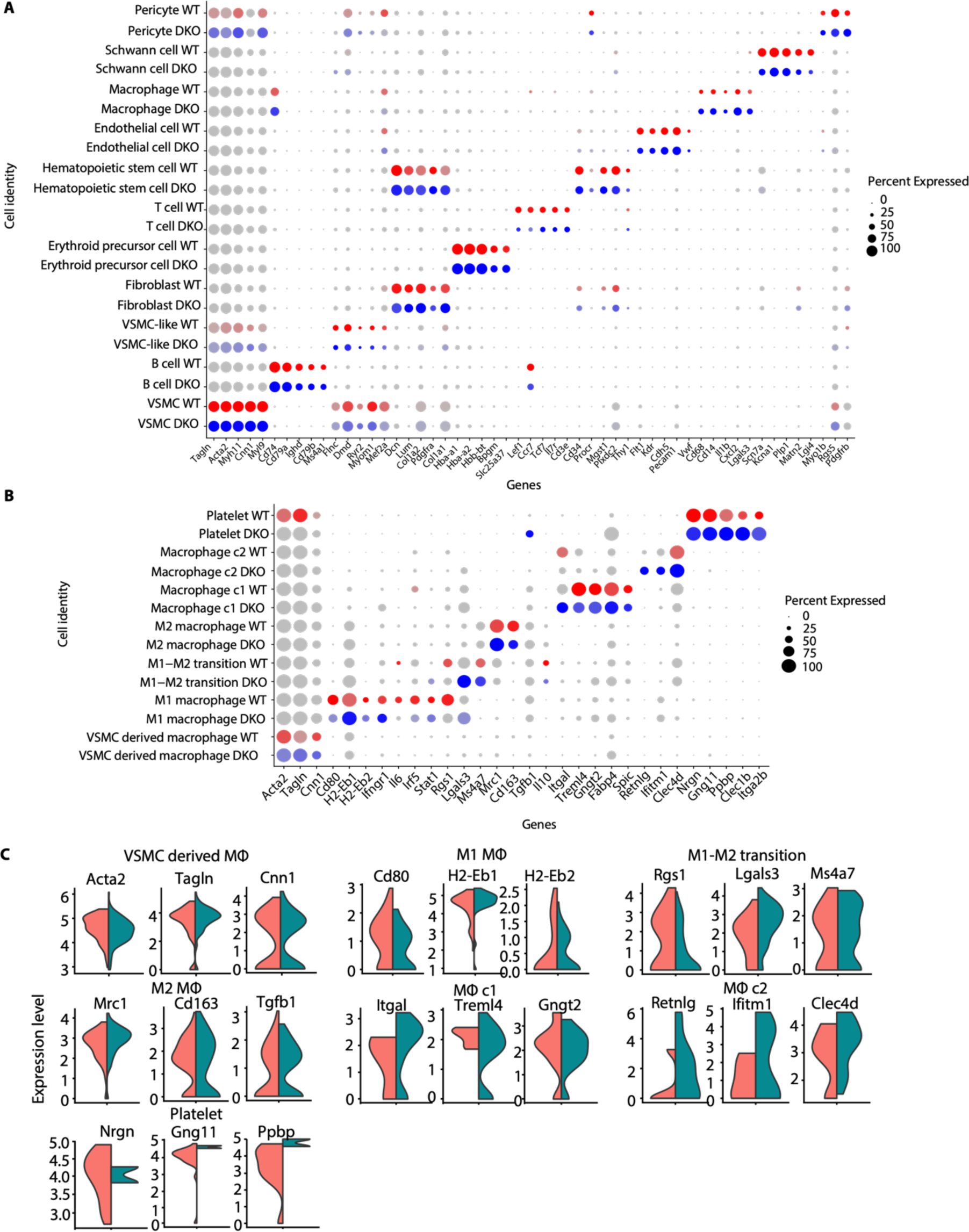
Cell populations were annotated based on marker gene expression. (**A**) Dot plot of known marker genes for each cell cluster in Figure 1A. Marker gene expression are colored by red and blue for WT and DKO, respectively. Size of nodes represent percentage of cells expressing a certain gene. (**B**) Dot plot of known marker genes for each macrophage subclusters. (**C**) Violin plots showing the representative marker gene expression in each macrophage subclusters in WT and DKO.

**Figure S2.**
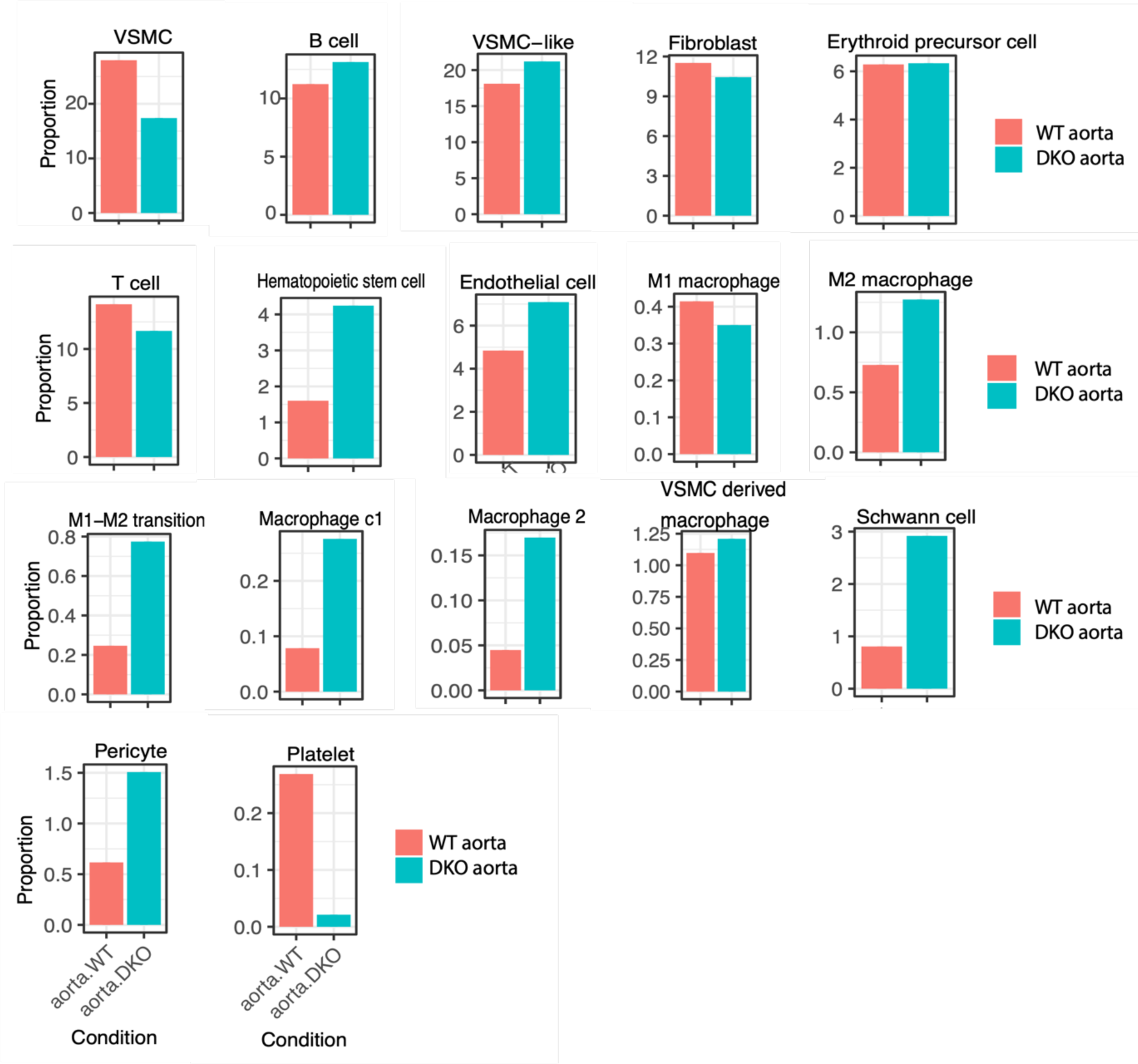
Cell proportion changes for each cell type in WT and DKO. The bar plots showing the cell proportion of M1 macrophage was decreased, while those of M2 macrophage and M1-M2 transition cells were increased in DKO compared to WT.

**Figure S3.**
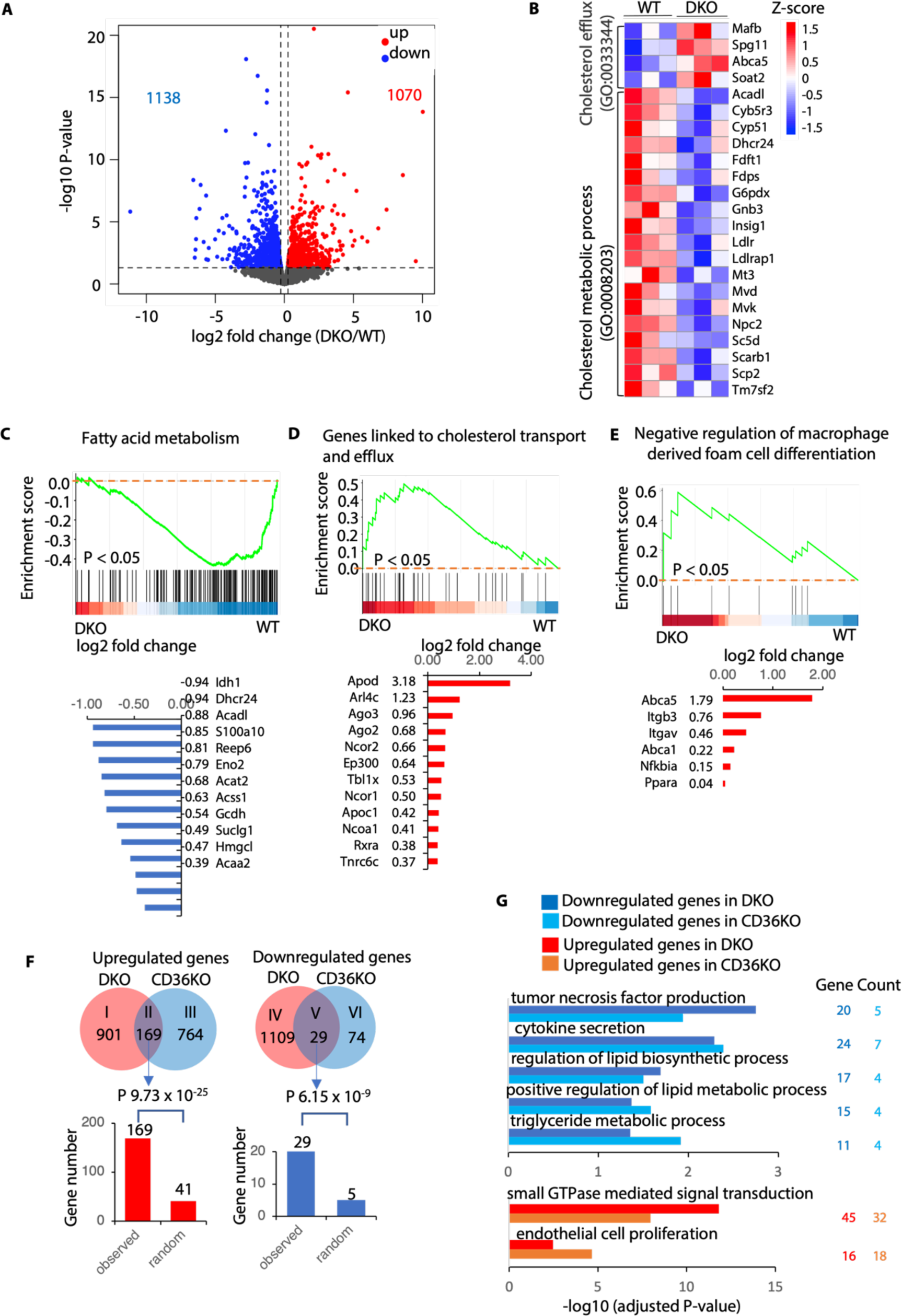
RNA-seq analysis of WT and DKO macrophages reveals that Epsins regulate lipid cholesterol metabolism and efflux pathways. (**A**) A volcano plot showing differential gene expression in Epsin deficient (DKO) and wild type (WT) macrophages. Red and blue indicate up- and down-regulated genes, respectively. (**B**) A heatmap showing the expression of genes involved in the cholesterol metabolic process (GO:0008203) and cholesterol efflux (GO:0033344). (**C-E**) GSEA demonstrated the tendency of individual pathways to be up- or down-regulated in Epsin deficient macrophages compared to wild type (top panels). The bar plots (bottom panels) showing log2 fold change of altered genes in these pathways. (**F**) Venn diagrams (top panels) showing the number of overlapping up- and down-regulated genes in DKO and CD36 knockout (CD36KO) samples. Bar plots (bottom panels) showing the observed numbers of overlapped genes versus numbers expected by random chance. The P-value was calculated using two tail Fisher exact test. (**G**) The Gene Ontology (GO) enrichment analysis for shared down regulated (top) and up regulated genes (bottom) between DKO and CD36KO macrophages. Benjamini corrected P-values are indicated on the x-axis.

**Figure S4.**
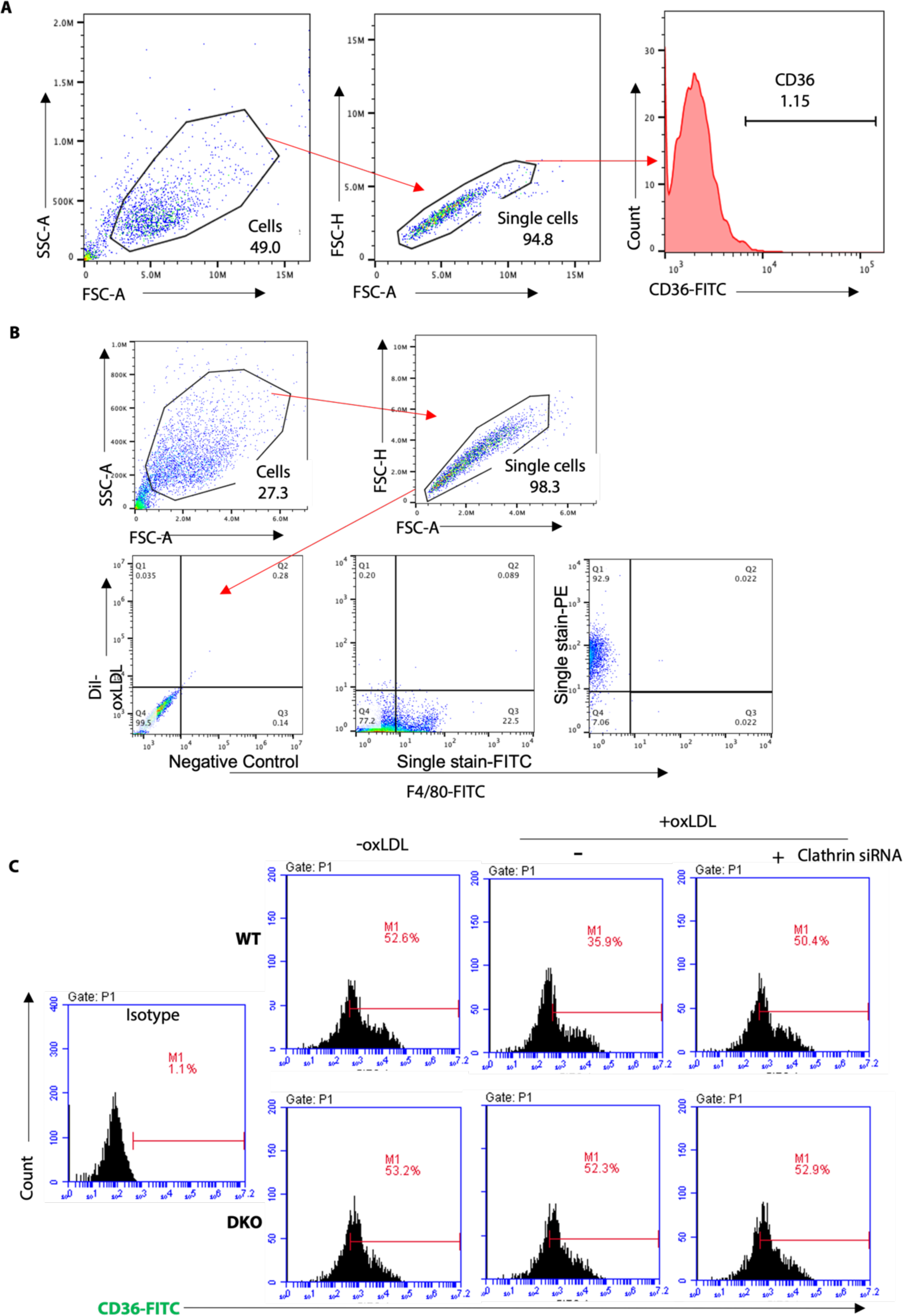
Gating strategy and clathrin-mediated CD36 internalization. (**A**) Macrophages isolated from WT (n=6) and DKO (n=6) mice were incubated in lipid-deficient medium for 24h followed by treatment with oxLDL for 2h, then staining with CD36-Alex488 and cytometric analyses. (**B**) Macrophages isolated from WT (n=6) and DKO (n=6) mice were incubated in lipid-deficient medium for 24h followed by the treatment of 10 μg/mL Dil-oxLDL for 2h at 37^0^C and assessed the uptake of lipoproteins by flow cytometry. The major macrophage population was selected in forward vs side scatter plots and single cell determination was performed by FSC-H vs FSC-A. CD36 (A) or DiI-oxLDL (B) positive macrophages were presented in Figure 2C and 2H, respectively. (**C**) WT and DKO macrophages were incubated in lipid-deficient medium and transfected with clathrin siRNA for 24h followed by treatment with or without 100μg/mL oxLDL for 15 mins at 37^0^C. Flow cytometry for surface level of CD36.

**Figure S5.**
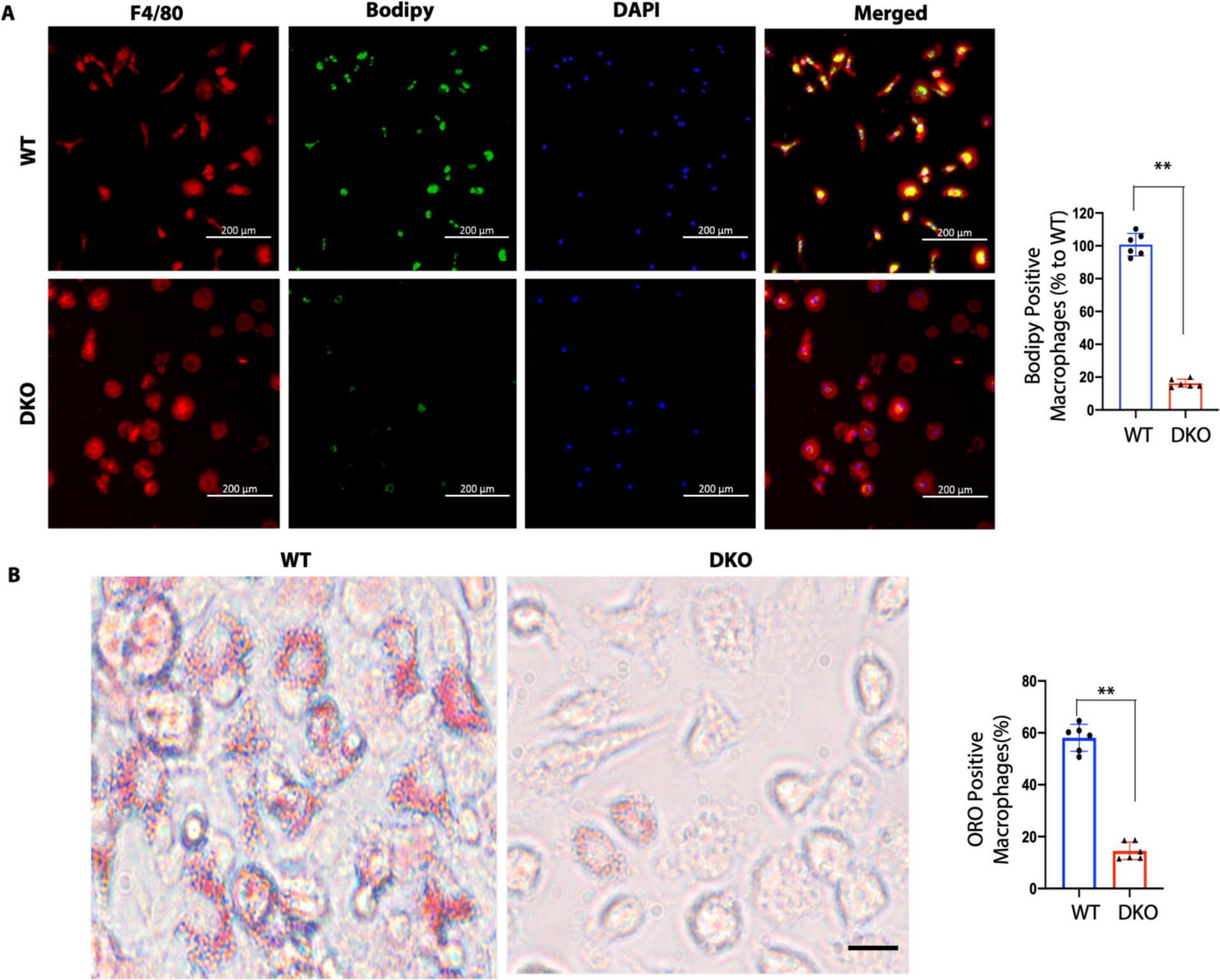
The loss of Epsins reduced oxLDL uptake by macrophages. (**A**) Isolated peritoneal macrophages from WT (n=6) and DKO (n=6) mice were pre-incubated with 25μg/mL oxLDL for 24h in lipid-deficient medium and stained with BODIPY (lipids, green), F4/80 (macrophage, red), and DAPI (blue), **WT vs DKO group, n=6, P<0.01, scale bar=200μm. (**B**) ORO staining of peritoneal macrophages, which were pre-incubated with 25μg/mL oxLDL for 24h in lipid-deficient medium, **WT vs DKO group, n=6, P<0.01, scale bar=20μm. Data are presented as mean ± SD and were analyzed using an unpaired Student’s t-test.

**Figure S6.**
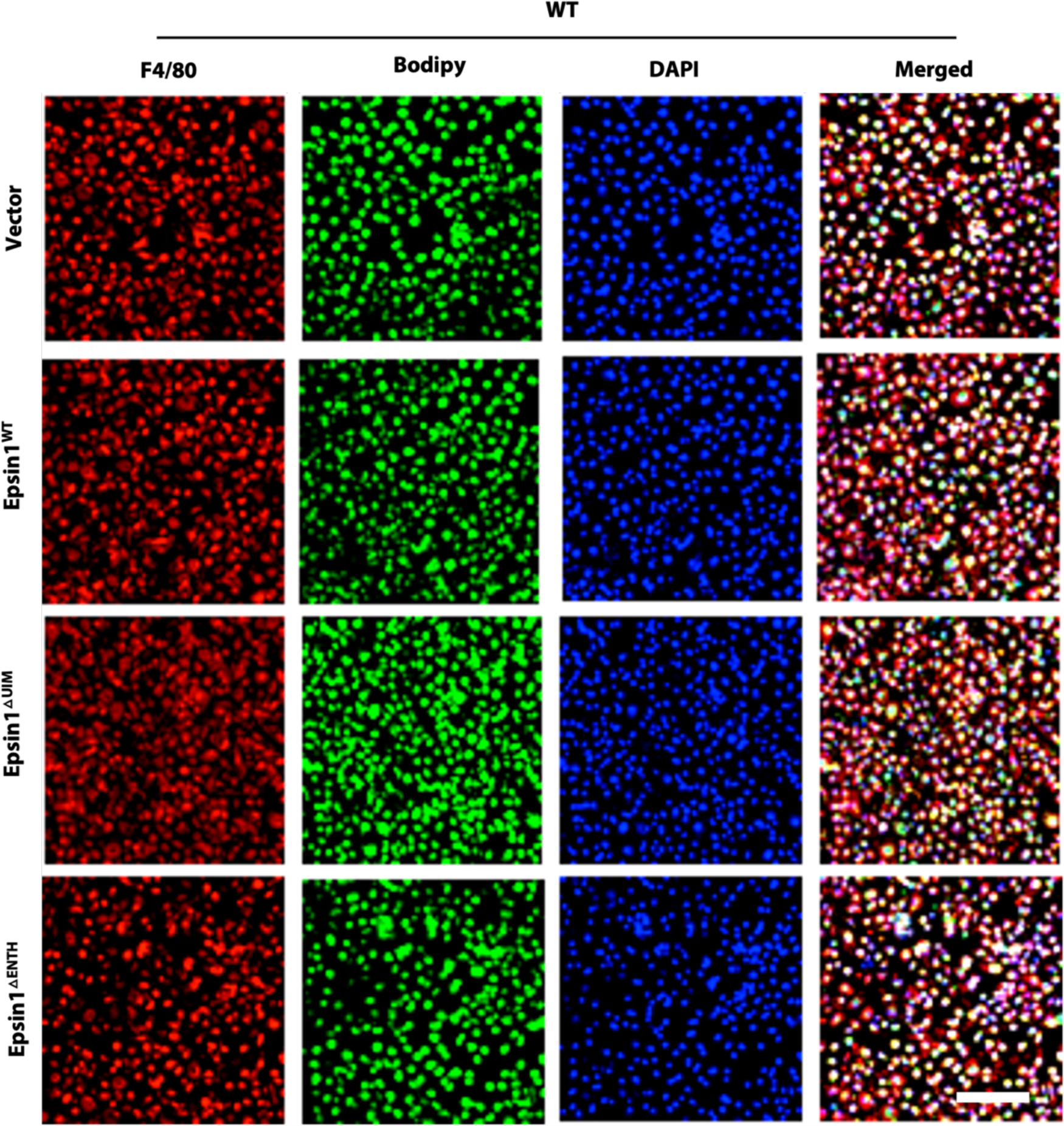
Transfection of full-length Epsin1 and constructs with the ENTH and UIM domains deleted into WT macrophages did not affect lipid uptake. Constructs of FLAG- Epsin1 WT, △ENTH, and △UIM were transfected into ApoE^-/-^/WT macrophages for 48h and treated with 100 μg/mL oxLDL for 1h, followed by staining with F4/80 (red), BODIPY (green) and DAPI (blue). Scale bars=200μm. Statistical analyses are presented in Figure 3E.

**Figure S7.**
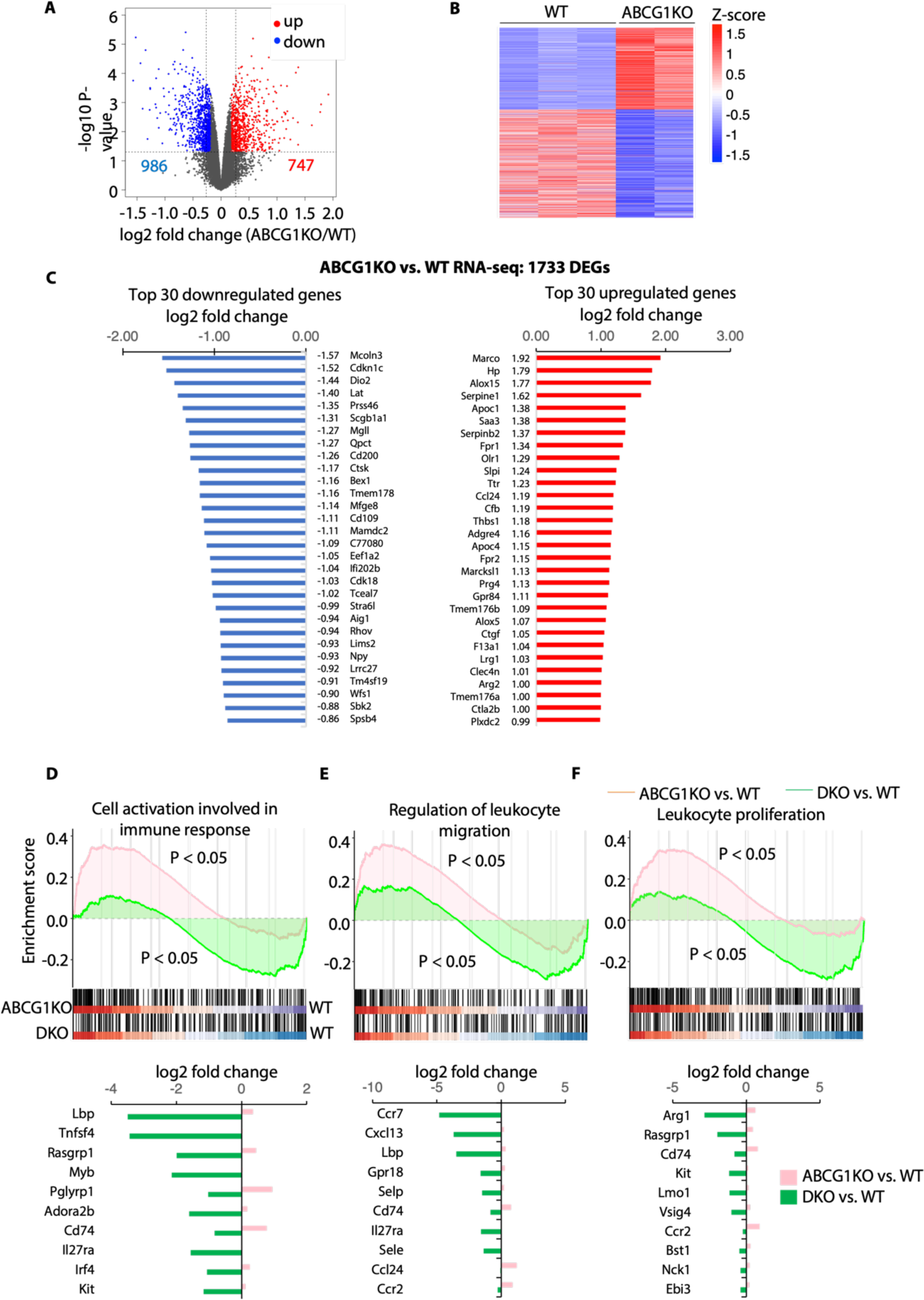
Differentially expressed genes between WT and ABCG1 knockout (ABCG1KO) macrophages. (**A**) A volcano plot showing differential gene expression in ABCG1KO and WT macrophages. Red and blue indicate up- and down-regulated genes, respectively. (**B**) A heatmap exhibiting the expression values of up- and down-regulated genes in each sample. (**C**) Bar plots showing log2 fold changes of the top 30 up- and down-regulated genes between ABCG1KO and WT macrophages. Up- (red) and down-regulated (blue) genes are indicated. (**D**-**F**) GSEA (top panels) indicated the tendency of individual pathways to be up- or down-regulated in response to DKO or ABCG1KO compared to WT cells. Genes associated with cell activation involved in immune response (GO:0002263) (D), regulation of leukocyte migration (GO:0002685) (E), and leukocyte proliferation (GO:0070661) (F) were analyzed. The bar plots (bottom panels) showing log2 fold change of altered genes in these pathways.

**Figure S8.**
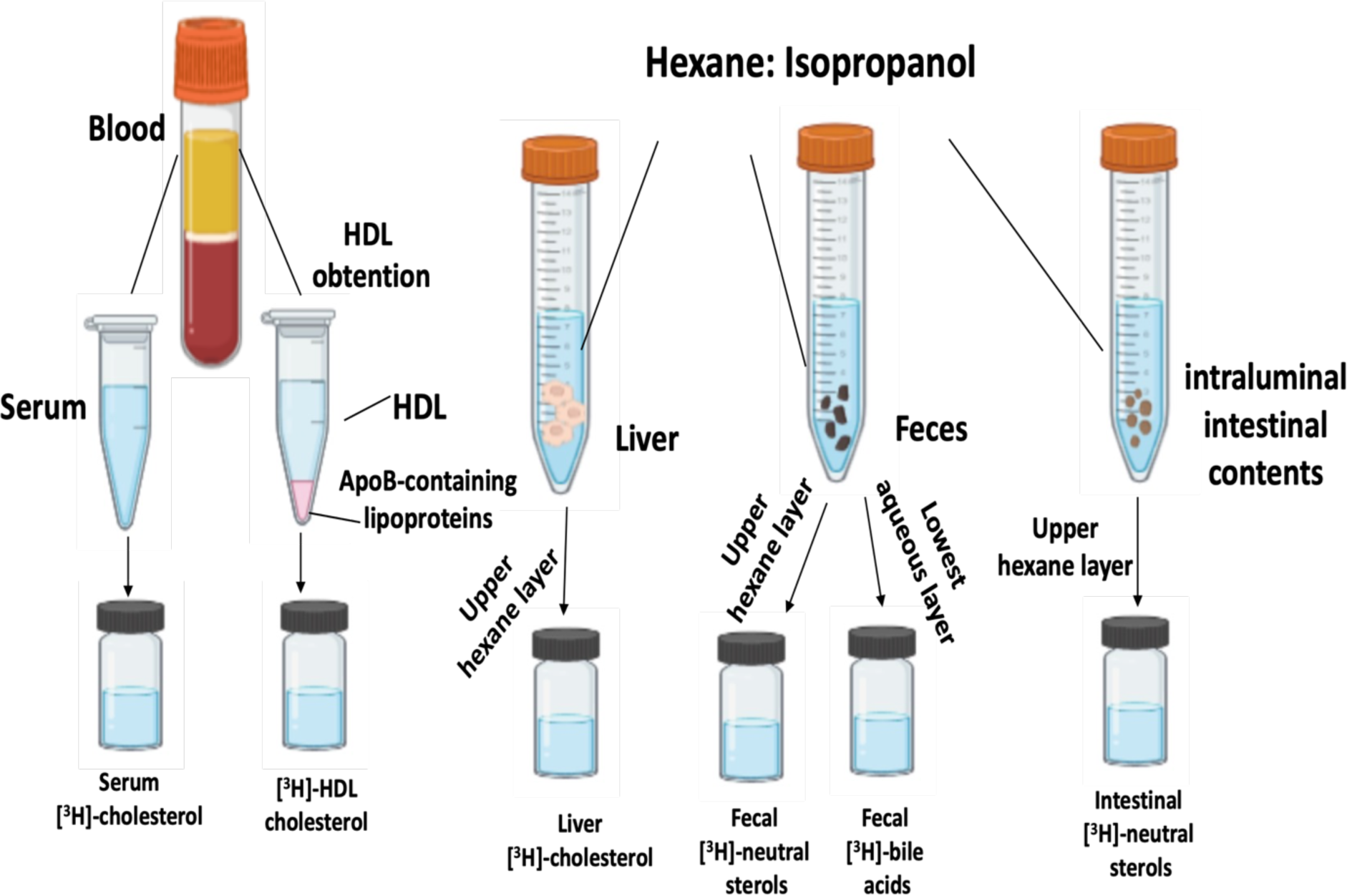
Schematic of the quantification for radiolabeled cholesterol. [^3^H]-cholesterol was measured in serum and HDL after precipitating ApoB-containing lipoproteins. Liver, feces and intestinal lipids were extracted with hexane–isopropanol and partitioned against Na_2_SO_4_. Liver [^3^H]-radioactivity was determined in the upper layer, which contains the [^3^H]-cholesterol. In the feces extract, the amount of [^3^H]-radioactivity was determined in the upper layer (neutral sterols) and the lowest layer (bile acids). In the intestinal contents, [^3^H]-radioactivity was determined in the upper layer.

**Figure S9.**
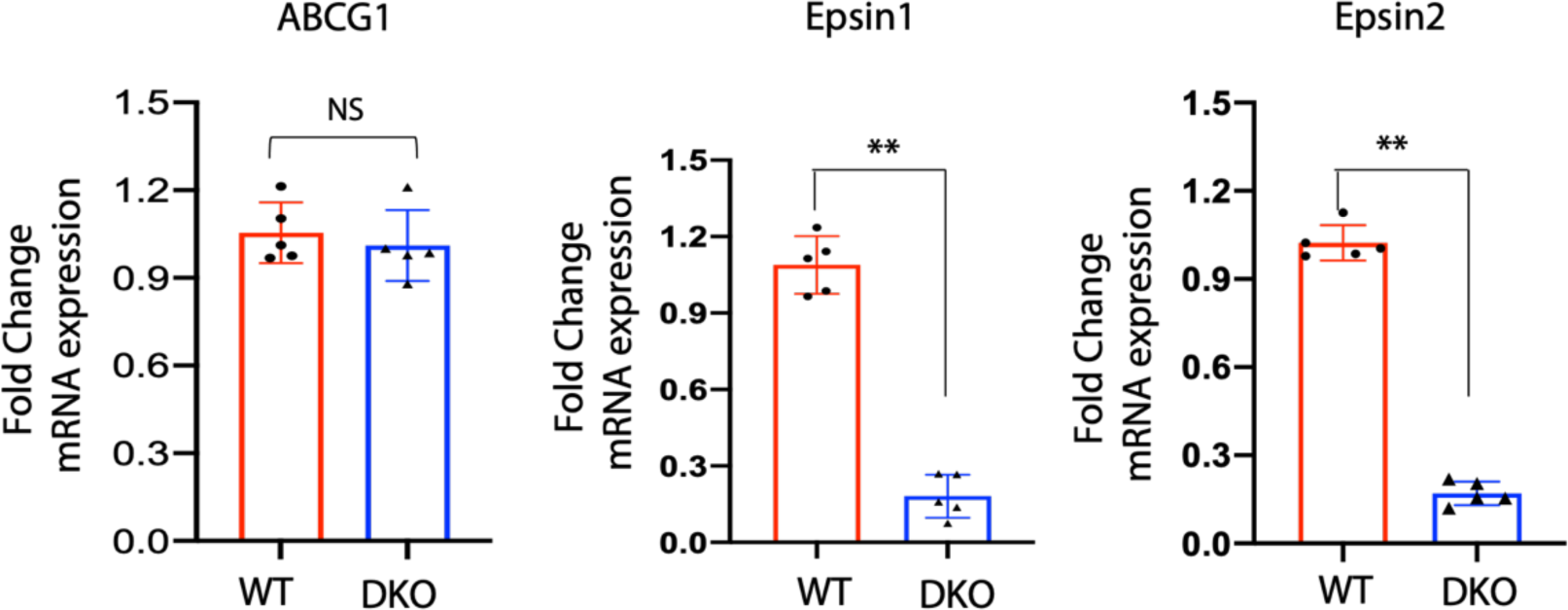
Gene expression levels of indicated genes in WT and DKO macrophage. Peritoneal macrophages were isolated from WT (n=5) and DKO (n=5) mice. Total RNA was extracted and mRNA levels of ABCG1, Epsin1, and Epsin2 were measured (NS=no significant difference, ** WT vs DKO group, n=5, P<0.01). Data are presented as mean ± SD and analyzed using an unpaired Student’s t-test.

**Figure S10.**
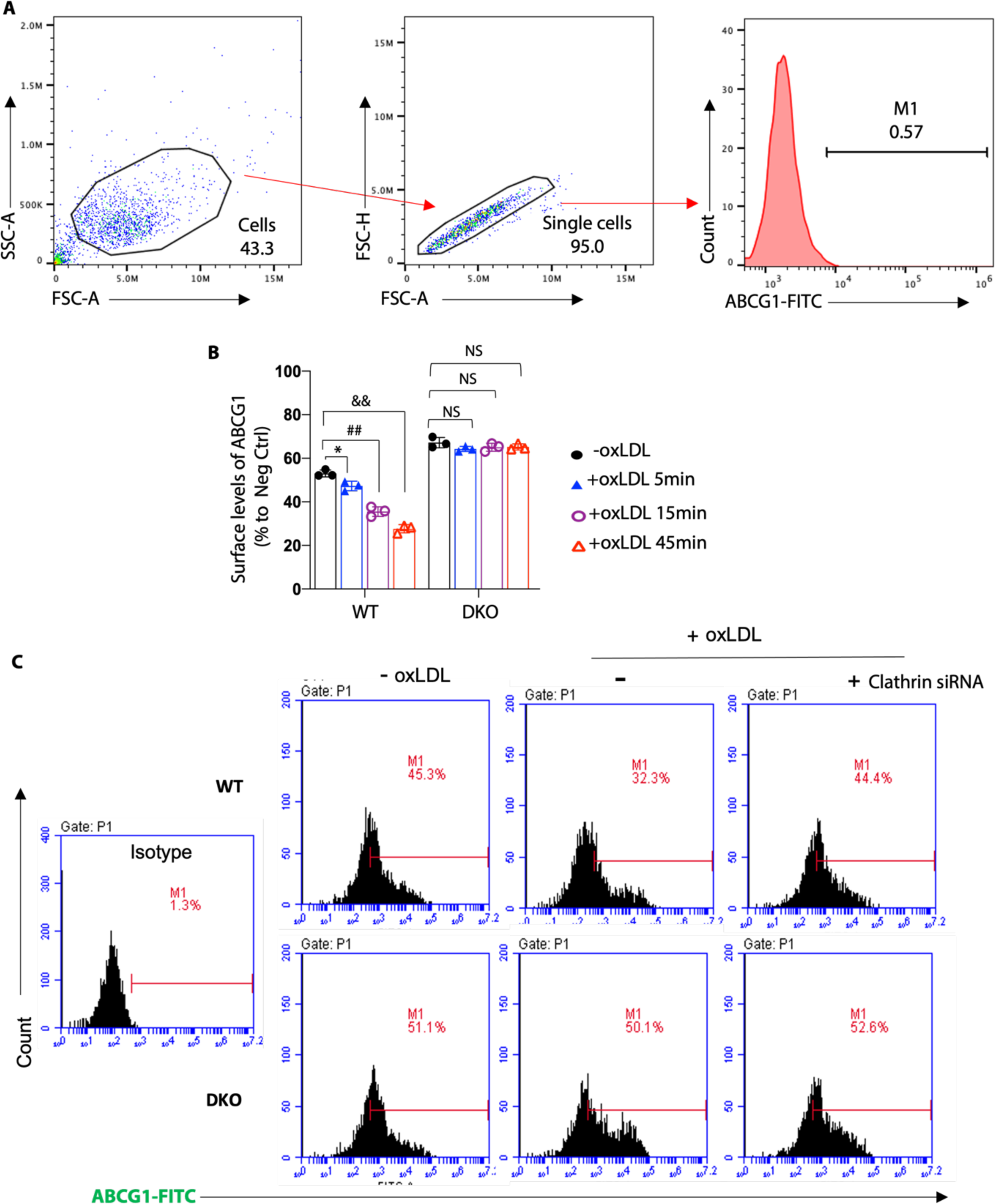
Gating strategy and clathrin mediated-ABCG1 endocytosis. (**A**) Macrophages isolated from WT and DKO mice were incubated in lipid-deficient medium and treated with LXR agonist for 24h followed by treatment with or without 100 μg/mL oxLDL for 5min, 15min, and 45min, and then stained with ABCG1. Surface levels of ABCG1 were assessed by flow cytometry. The major macrophage population was selected in forward versus side scatter plots and single cell determination was performed by FSC-H vs FSC-A. Isotype controls were gated in the histogram as M1 for the negative control of experimental groups for Figure 6C. (**B**) Statistical analysis is presented for Figure 6C. At least three independent experiments were performed for statistical analysis (*-oxLDL vs +oxLDL 5min, P<0.05; ^##^-oxLDL vs +oxLDL 15min, P<0.01; ^&&^-oxLDL vs +oxLDL 45min, P<0.01). Data are presented as mean ± SD and analyzed using one-way ANOVA. **(C)** WT and DKO macrophages were incubated in lipid-deficient medium and transfected with clathrin siRNA for 24h followed by treatment with or without 100μg/mL oxLDL for 15 mins at 37^0^C. Flow cytometry for surface level of ABCG1.

**Figure S11.**
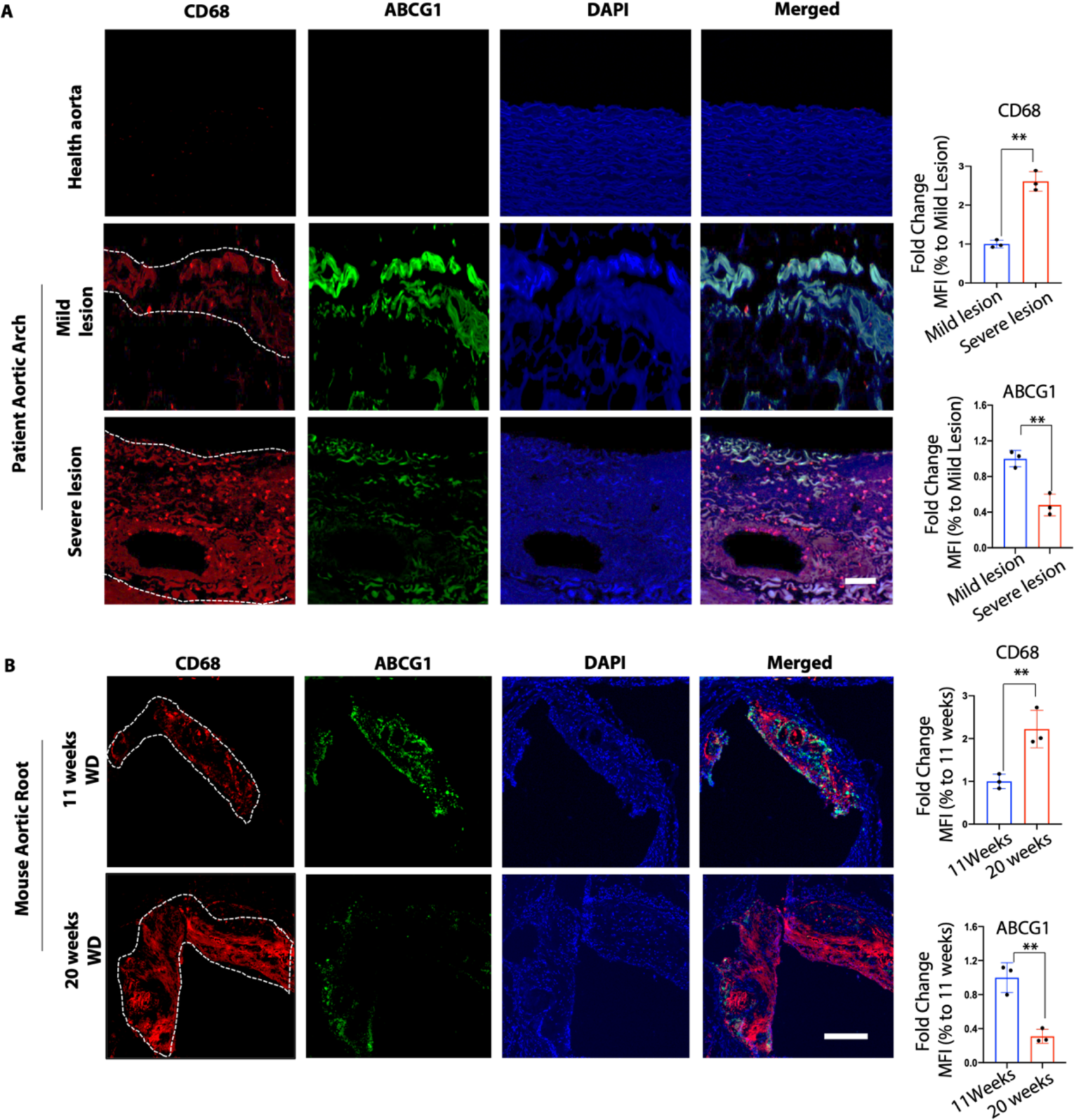
The expression of ABCG1 decreased with progression of atherosclerotic lesions in humans and mice. (**A-B**) Immunostaining of CD68 (red), ABCG1(green) and DAPI (blue) of human patient aortic arch sections (A, n=3) and mouse aortic root sections (B, n=3) in early and advanced stage of atherosclerosis (white dashed line outlined in CD68). Mean fluorescence intensity (MFI), **Mild vs Severe lesion, P<0.01, scale bar, A=50μm, B=200μm. Data are presented as mean ± SD and were analyzed using an unpaired Student’s t-test.

**Figure S12.**
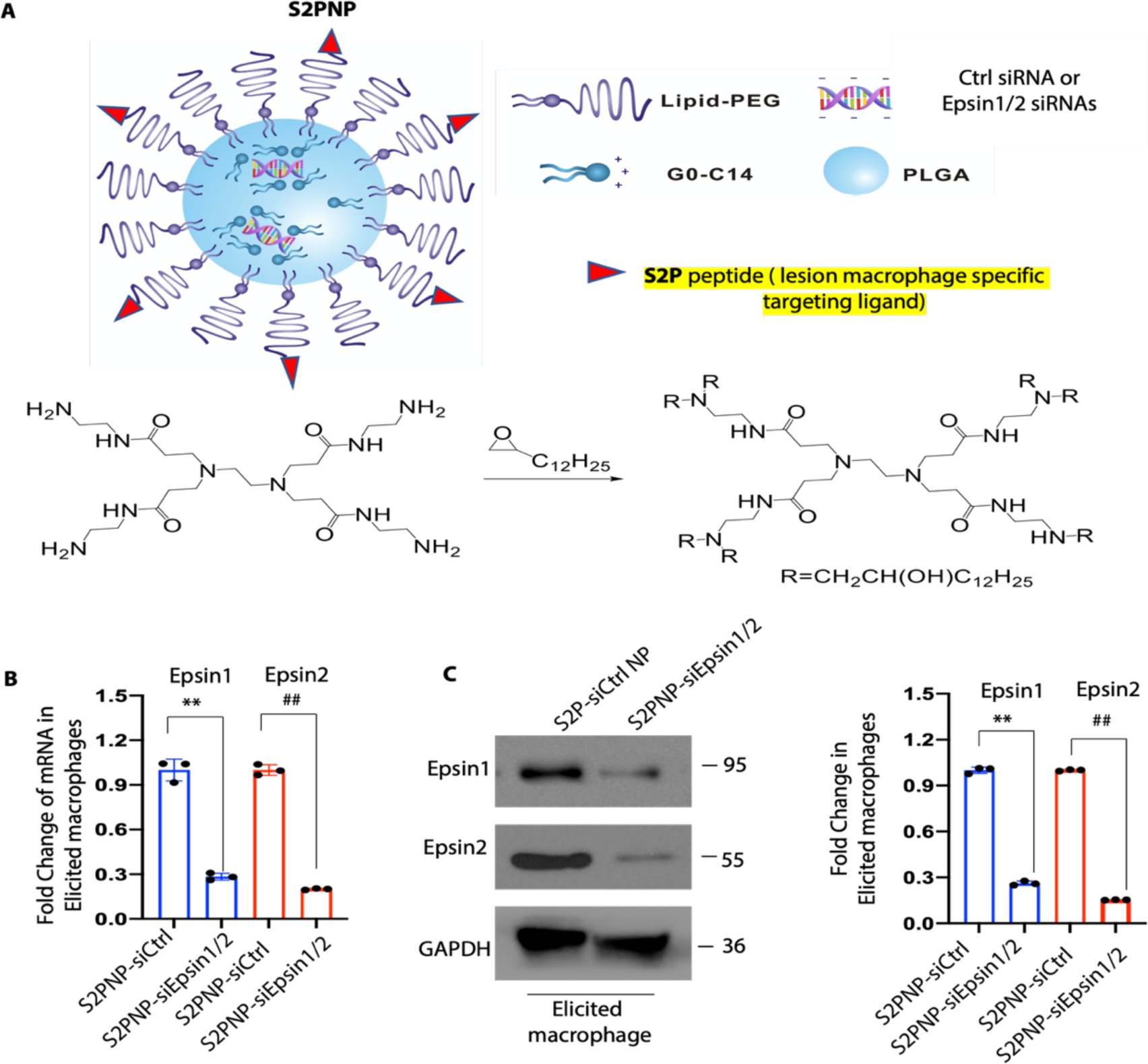
Characterization and silencing efficacy of S2P-conjugated siEpsin1/2 NPs. (**A**) Schematic of the targeted hybrid siRNA NP platform composed of a lipid-PEG shell with a lesion macrophage specific targeting ligand, S2P peptide, and a PLGA core. (**B-C**) Macrophages from WT mice were treated with S2PNP-siCtrl or S2PNP-siEpsin1/2 for 24h, RNA and proteins were isolated, qRT-PCR (B) and western blot (C) was performed to check the expression of Epsin 1 and 2 levels (n=3), **S2PNP-siCtrl vs S2PNP-siEpsin1/2 for Epsin1 expression, P<0.01; ^##^S2PNP- siCtrl vs S2PNP-siEpsin1/2 for Epsin2 expression, P<0.01. Data are presented as mean ± SD and analyzed using an unpaired Student’s t-test.

**Figure S13.**
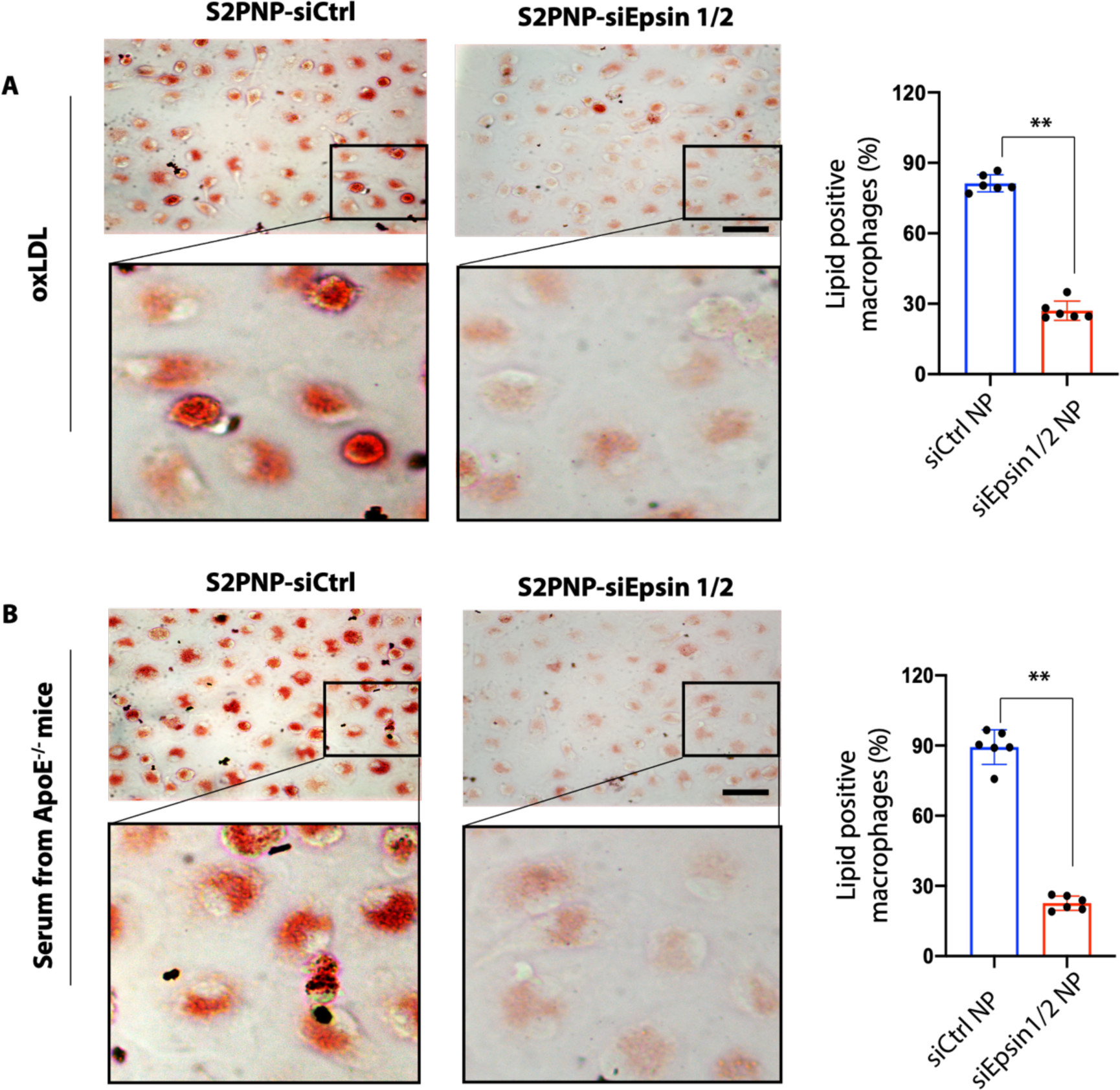
S2PNP-siEpsin1/2 treated macrophages show reduced foam cell formation. (**A- B**) Macrophages isolated from ApoE^-/-^ mice were incubated in lipid-deficient medium and treated with S2PNP-siCtrl or S2PNP-siEpsin1/2 for 48h, follow with treatment of 100 μg/mL oxLDL (A) or serum (B) collected from ApoE^-/-^ mice fed a WD for 8 weeks. ORO staining was performed (** S2PNP-siCtrl vs S2PNP-siEpsin1/2 group, n=6, P<0.01, scale bar=50μm). Data are presented as mean ± SD and analyzed using an unpaired Student’s t-test.

**Figure S14.**
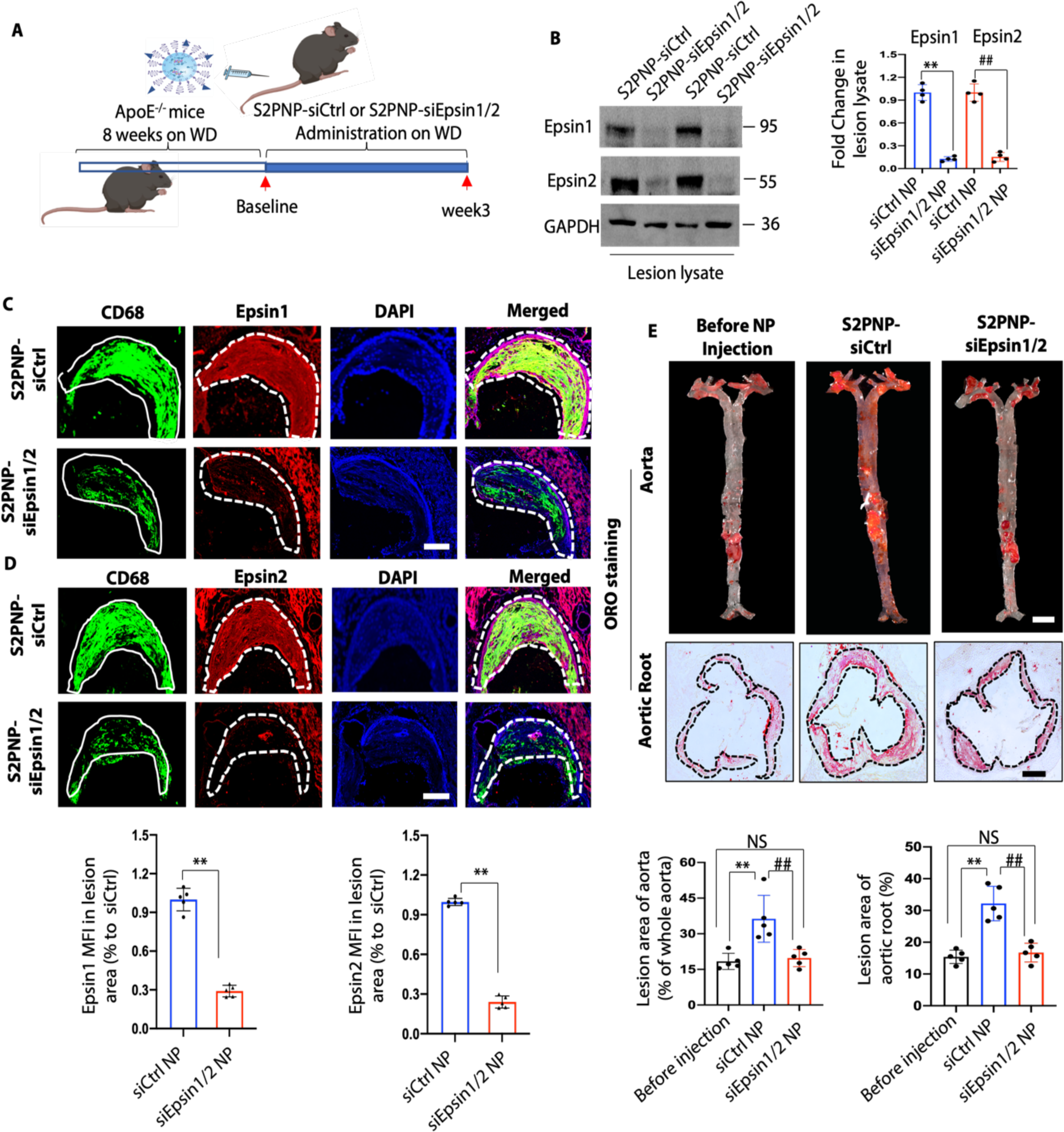
S2PNP-siEpsin1/2 inhibits lesion formation and macrophage accumulation in early stage of atherosclerosis. (**A**) ApoE^-/-^ mice were fed a WD for 8 weeks followed by treatment with S2PNP-siCtrl or S2PNP-siEpsin1/2 for 3 weeks (2 doses per week). (**B**) WB of Epsin 1 and 2 after treatment with S2PNP-siCtrl or S2PNP-siEpsin1/2 NPs using lesional lysates from the aortas (n=4 times, **S2PNP-siCtrl vs S2PNP-siEpsin1/2 for Epsin1, P<0.01; ^##^S2PNP-siCtrl vs S2PNP-siEpsin1/2 for Epsin2, P<0.01). (**C-D**) Aortic roots from S2PNP-siCtrl treated ApoE^-/-^ or S2PNP-siEpsin1/2 siRNA treated ApoE^-/-^ mice were stained with the macrophage marker CD68 (solid white line) and Epsin1 or Epsin2 (dashed white line), Epsin 1 and 2 mean fluorescence intensity (MFI) were analyzed (n=5, **S2PNP-siCtrl vs S2PNP-siEpsin1/2 for Epsin1, P<0.01, scale bars=500 μm). Data from B-D are presented as mean ± SD and analyzed using an unpaired Student’s t-test. (**E**) *En face* ORO staining of aortas (upper panel) and aortic root sections (lower panel) of hearts from baseline, S2PNP-siCtrl or S2PNP-siEpsin1/2 treated ApoE^-/-^ mice fed a WD. Lesion area was analyzed using one-way ANOVA (n=5, **S2PNP-siCtrl vs baseline, P<0.01, ^##^S2PNP-siCtrl vs S2PNP-siEpsin1/2 group P<0.01, Scale bar; aorta=5mm, aortic root=500μm). Data from E are presented as mean ± SD and were analyzed using one-way ANOVA.

**Figure S15.**
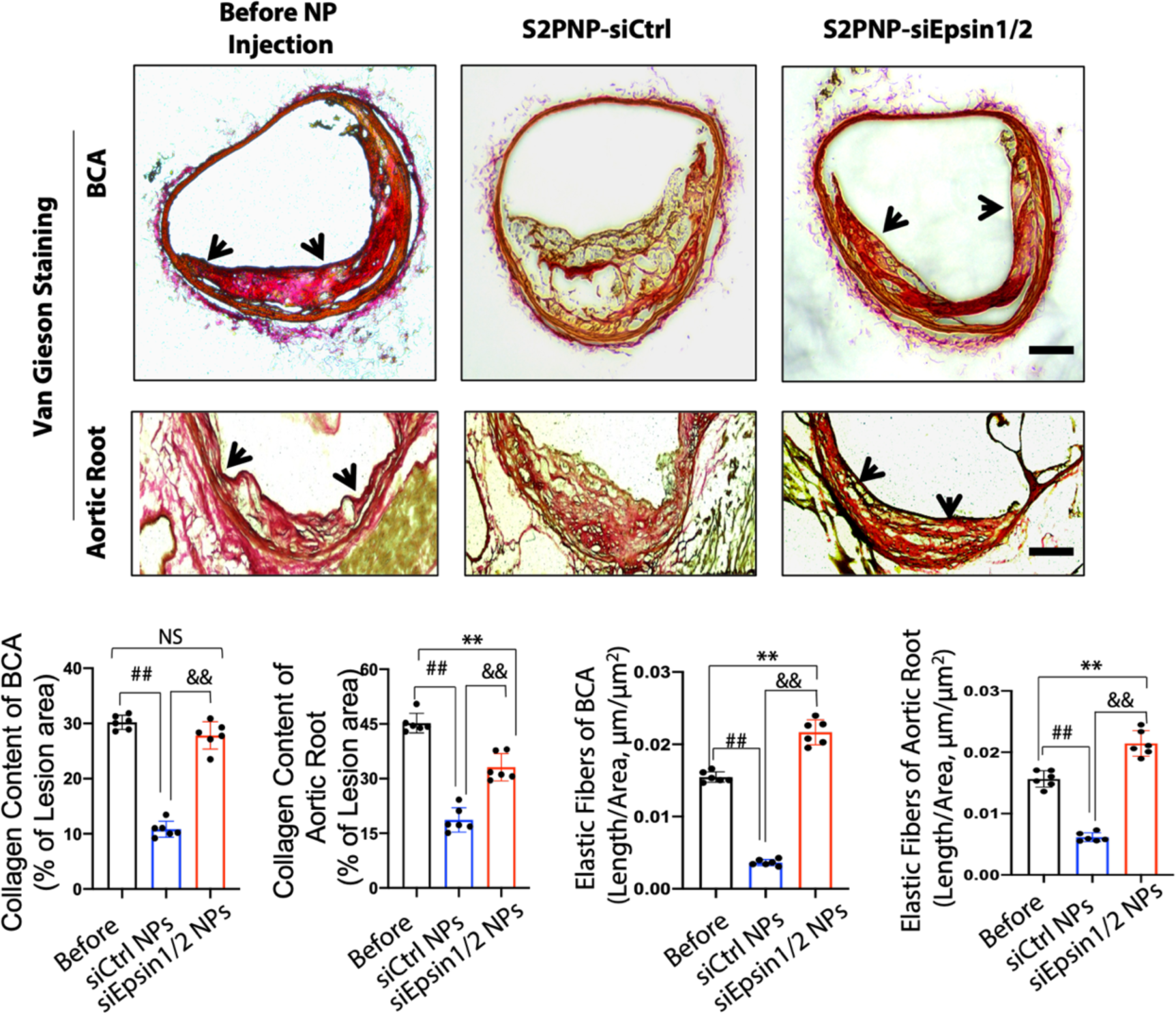
Silencing lesional macrophage Epsin1/2 by S2PNP-siEpsin1/2 NP-treatment stabilized plaques in a progression model of atherosclerosis. ApoE^-/-^ mice fed a WD for 17 weeks followed by treatment with S2PNP-siCtrl or S2PNP-siEpsin1/2 for 3 weeks (two doses per week). Van Gieson’s staining of brachiocephalic artery (BCA) (upper panel) and aortic root (lower panel) sections from baseline, S2PNP-siCtrl-, or S2PNP-siEpsin1/2-treated ApoE^-/-^ mice was performed (arrows indicate the elastic fibers, n=6, **baseline vs S2PNP-siEpsin1/2 group, P<0.01; ^##^baseline vs S2PNP-siCtrl group, P<0.01; ^&&^S2PNP-siCtrl vs S2PNP-siEpsin1/2 group, P<0.01, scale bar=250 μm). Data are presented as mean ± SD and were analyzed using one-way ANOVA.

**Figure S16.**
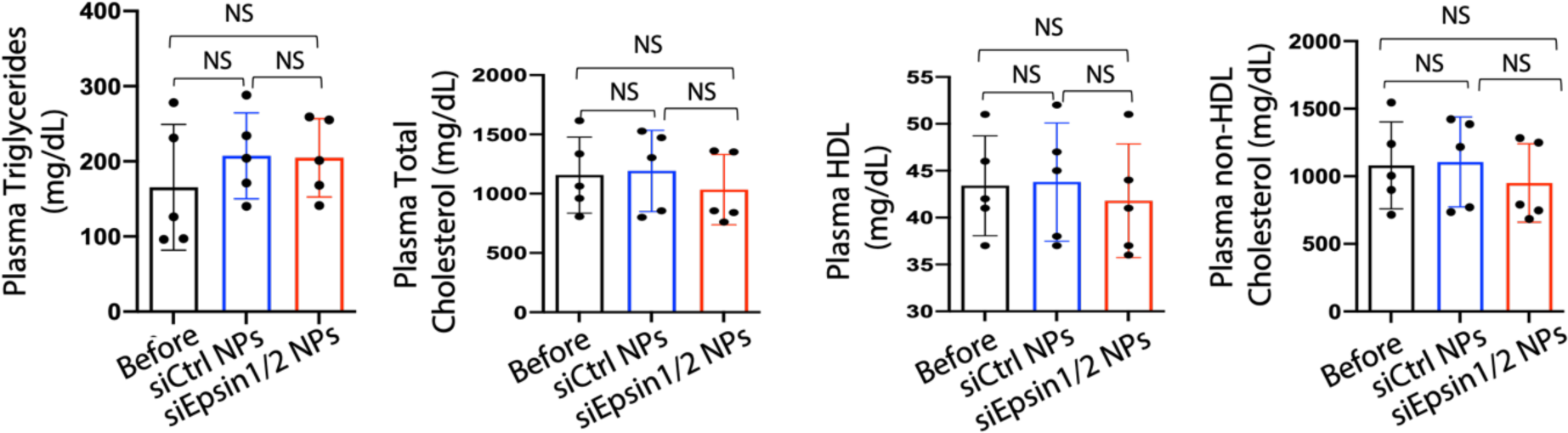
Delivery of S2PNP-siEpsin1/2 does not change cholesterol and triglyceride levels of ApoE^-/-^ mice fed a WD. Plasma from ApoE^-/-^ mice fed a WD for 17 weeks (before NP injection), followed by an additional 3 weeks of treatment with S2PNP-siCtrl or S2PNP-siEspsin1/2, showed no alteration in triglyceride, cholesterol, HDL and non-HDL (LDL/VLDL) cholesterol levels (n=5; NS, no significant difference). Data are presented as mean ± SD and were analyzed using a one-way ANOVA.

**Figure S17.**
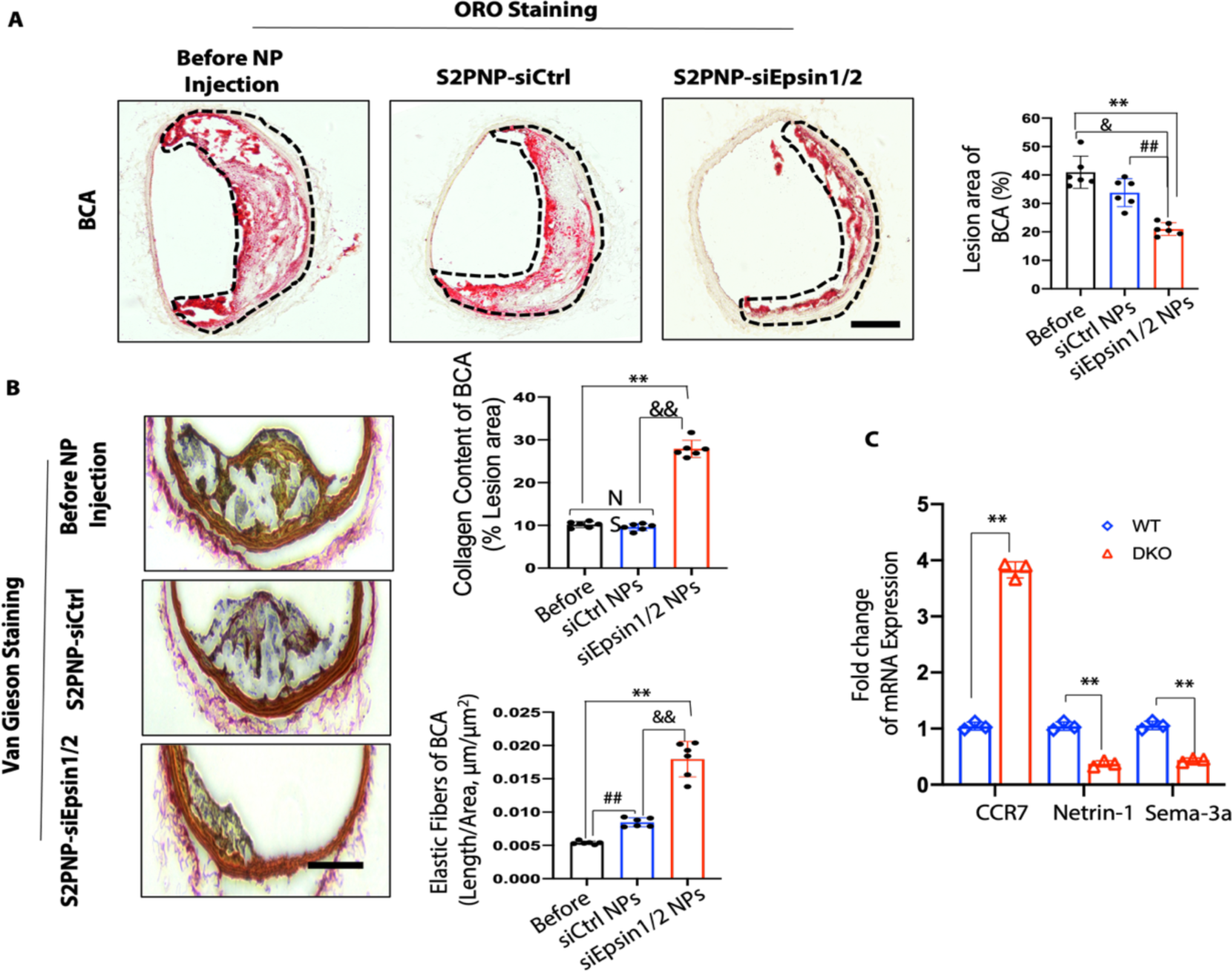
Silencing macrophage Epsin1/2 by S2PNP-siEpsin1/2 NPs reduces lesion size in a regression model of atherosclerosis. (**A**) ORO staining of BCA sections in baseline, S2PNP- siCtrl or S2PNP-siEpsin1/2 treated PCSK9-mice was performed and lesions were indicated (dash lines) (n=6, **Baseline vs S2PNP-siEpsin1/2 group, P<0.01; ^&^Baseline vs S2PNP-siCtrl group, P<0.05; ^##^S2PNP-siEpsin1/2 vs S2PNP-siCtrl group, P<0.01; scale bar=500μm). (**B**) Van Gieson’s staining of BCA sections from above three groups was performed (n=6, **Baseline vs S2PNP- siEpsin1/2 group, P<0.01; ^##^Baseline vs S2PNP-siCtrl group, P<0.01; ^&&^S2PNP-siEpsin1/2 vs S2PNP-siCtrl, P<0.01; scale bar=500μm). Data in A and B are presented as mean ± SD and analyzed using one-way ANOVA. (**C**) qRT-PCR analysis to confirm expression of the indicated genes (n=3, **DKO vs WT group, P<0.01). Data are presented as mean ± SD and analyzed using an unpaired Student’s t-test.

**Figure S18.**
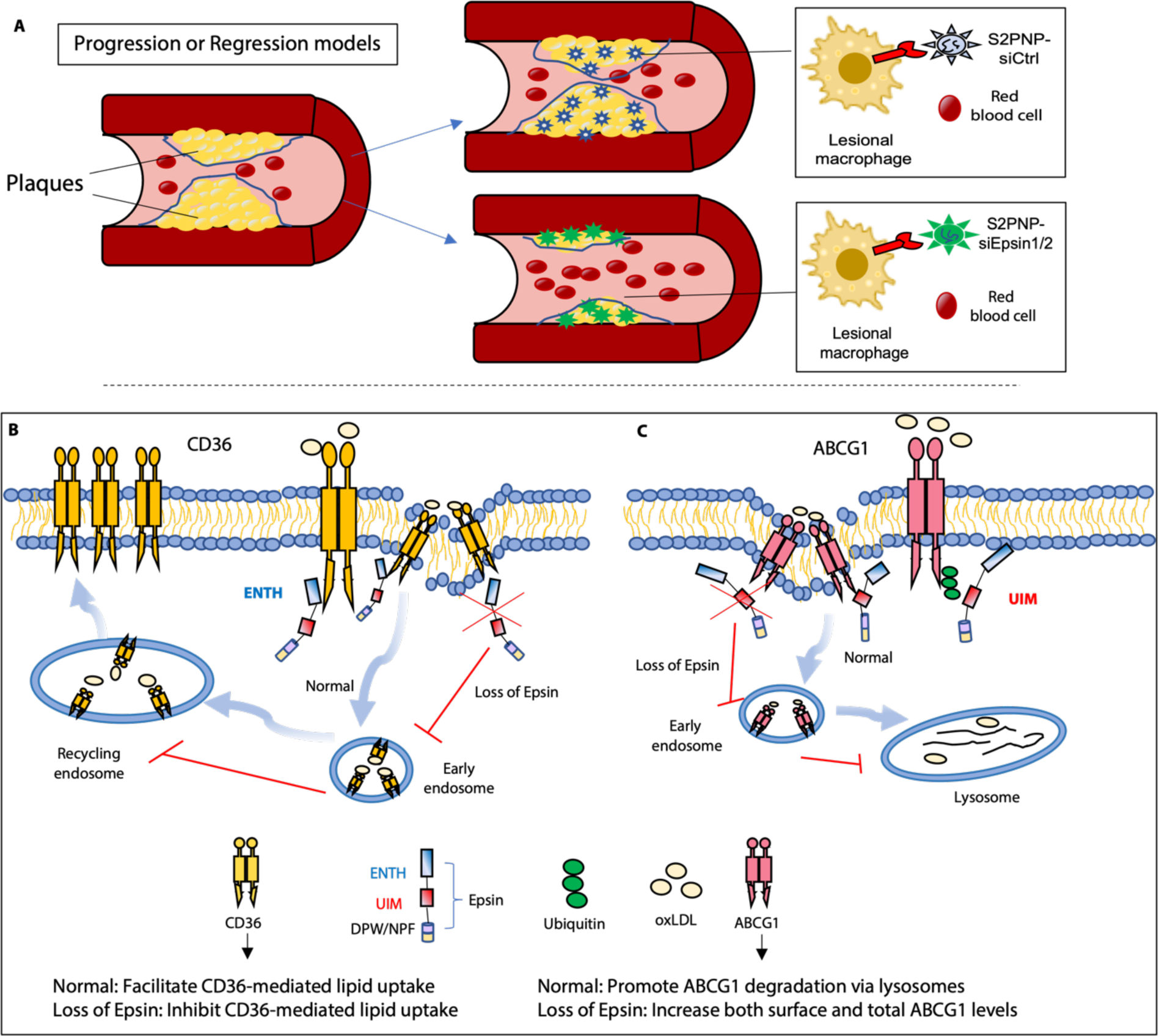
Summary schematic diagram of the study. (**A**) In the progression model of atherosclerosis, the plaque size and lesions increase in the artery on western diet feeding. The delivery of S2PNP-siEpsin1/2 significantly slowed the progression of atherosclerosis compared to S2PNP-siCtrl group. In the regression model of atherosclerosis, plaque size and lesion area were dramatically reduced with the intravenously injection of S2PNP-siEpsin1/2. (**B-C**). Under the stimuli of oxLDL, Epsin binds to CD36 and ABCG1 through Epsin ENTH and Epsin UIM domains, respectively. In B, Epsin facilitates CD36-mideated lipid uptake via recycling endosomes. The loss of Epsin impairs the internalization of CD36, which results in reduced lipid uptake. While in C, Epsin promotes endocytic degradation of ABCG1via lysosomes, which leads to reduced total and surface level of ABCG1.

**Table S1.**
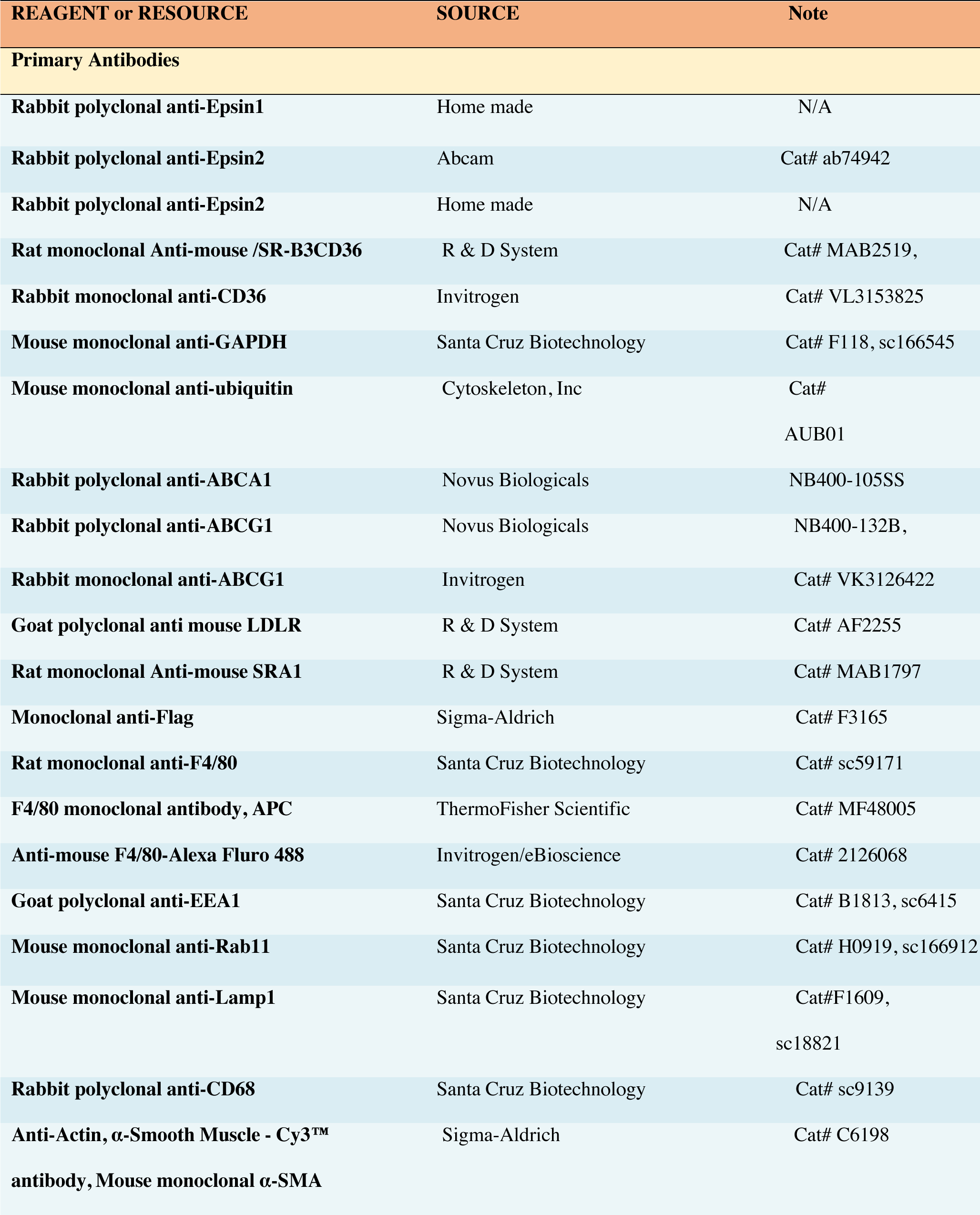
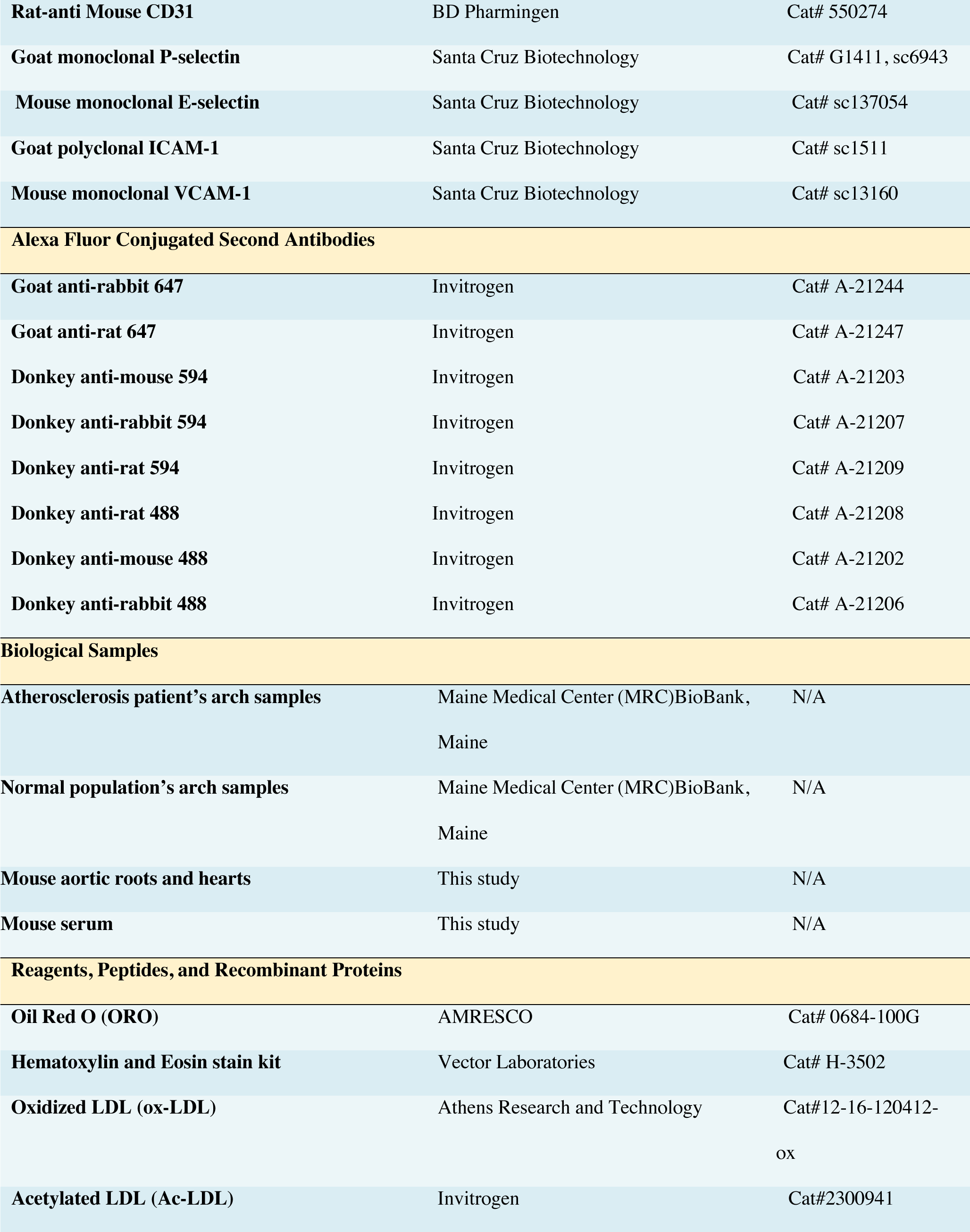
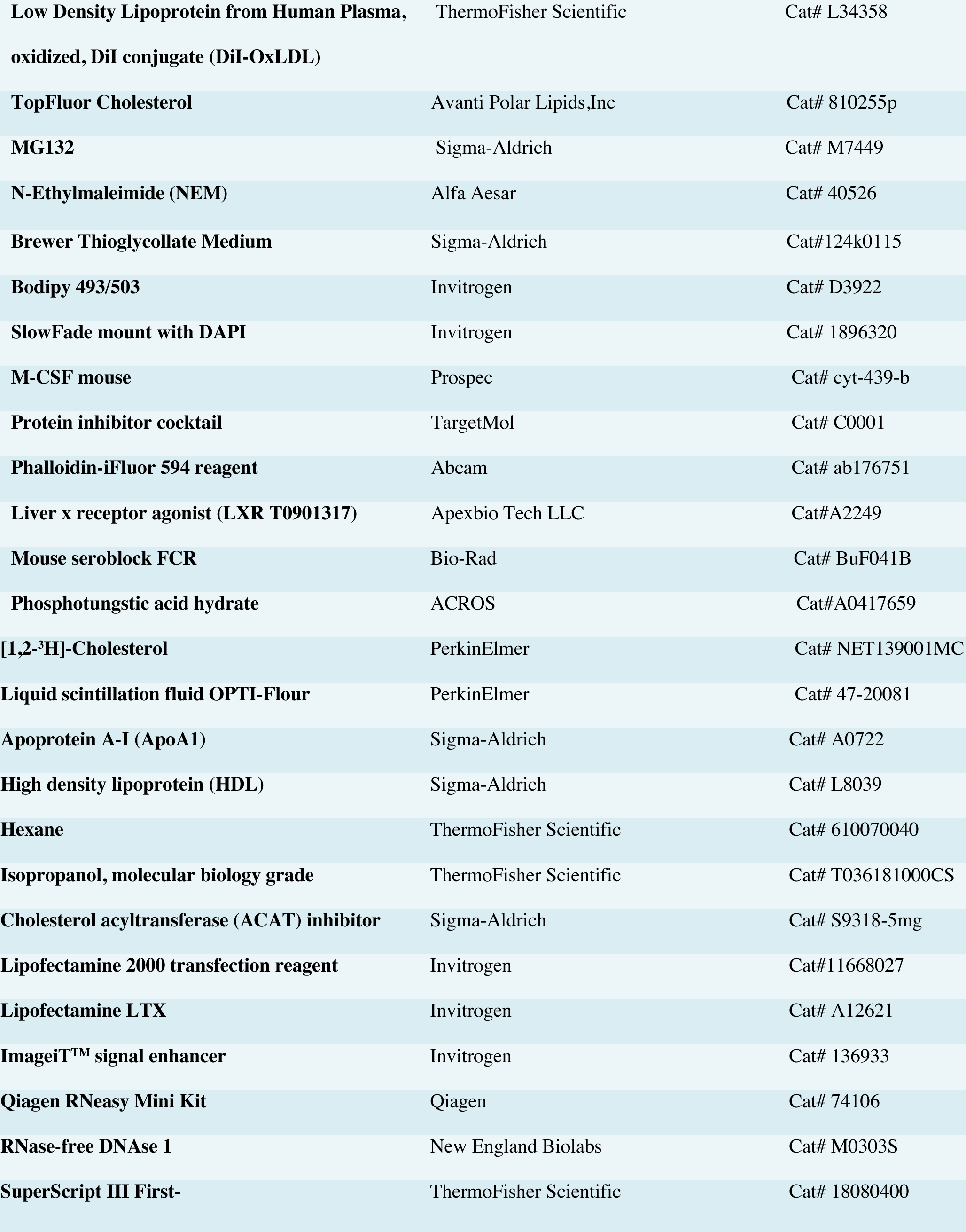
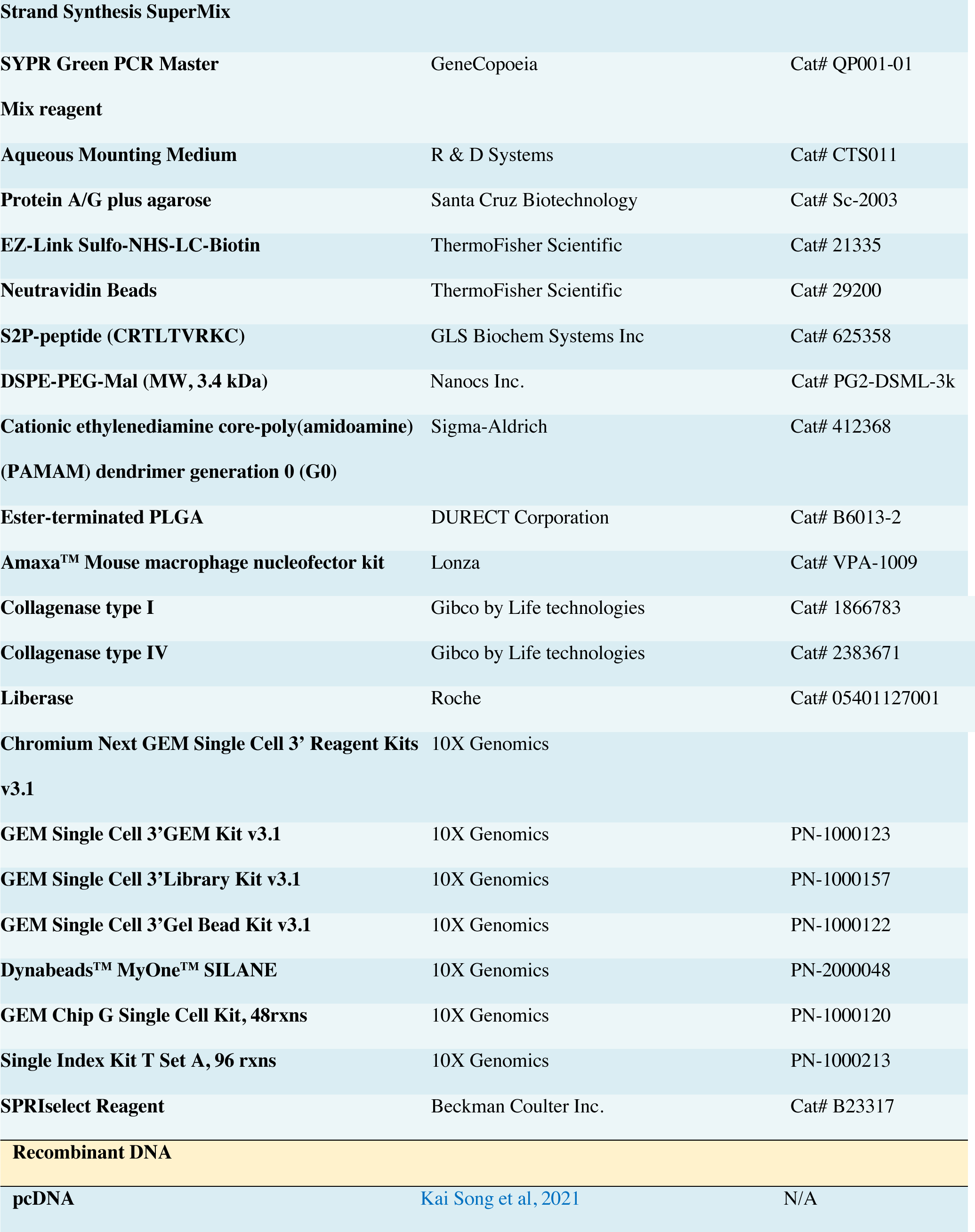
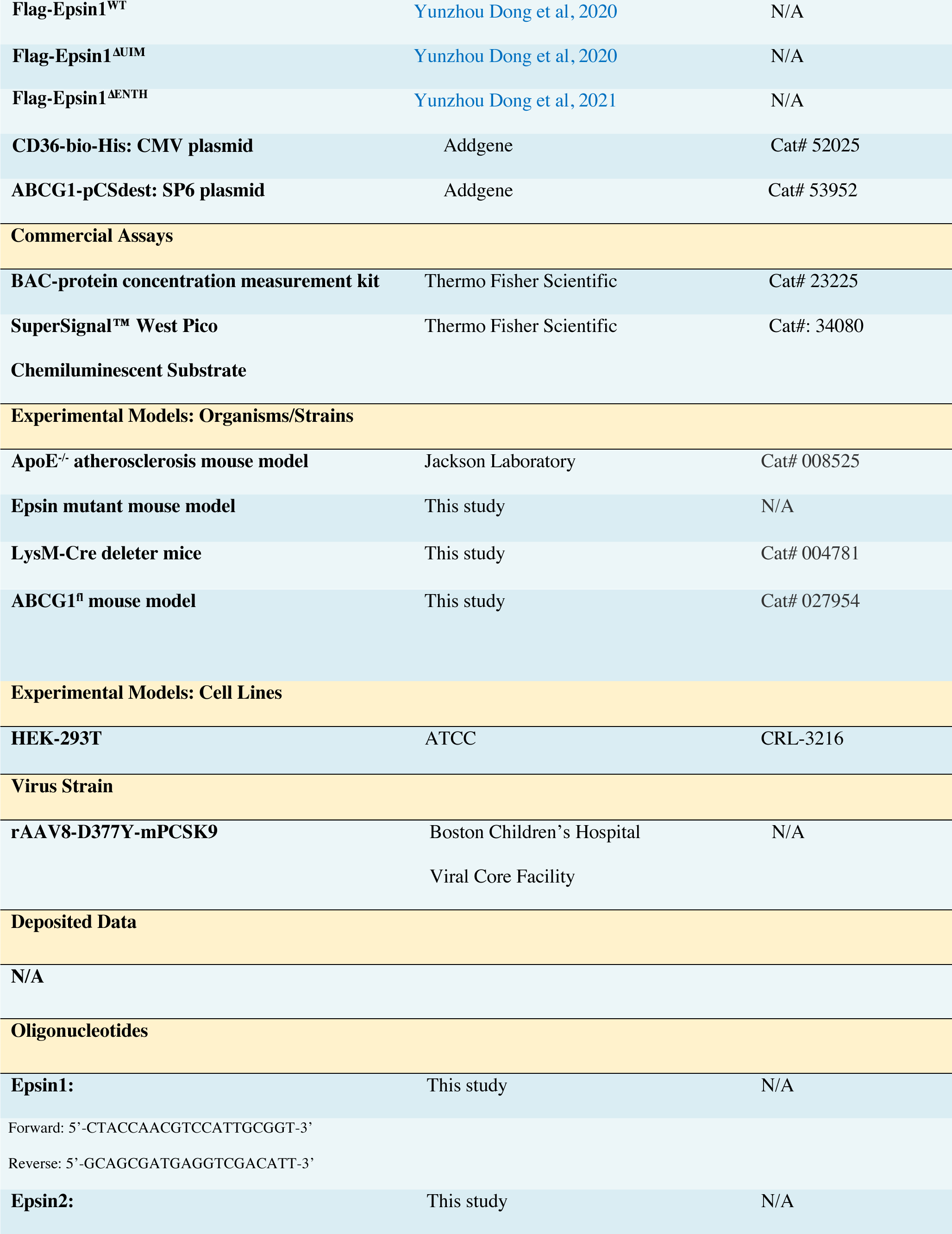
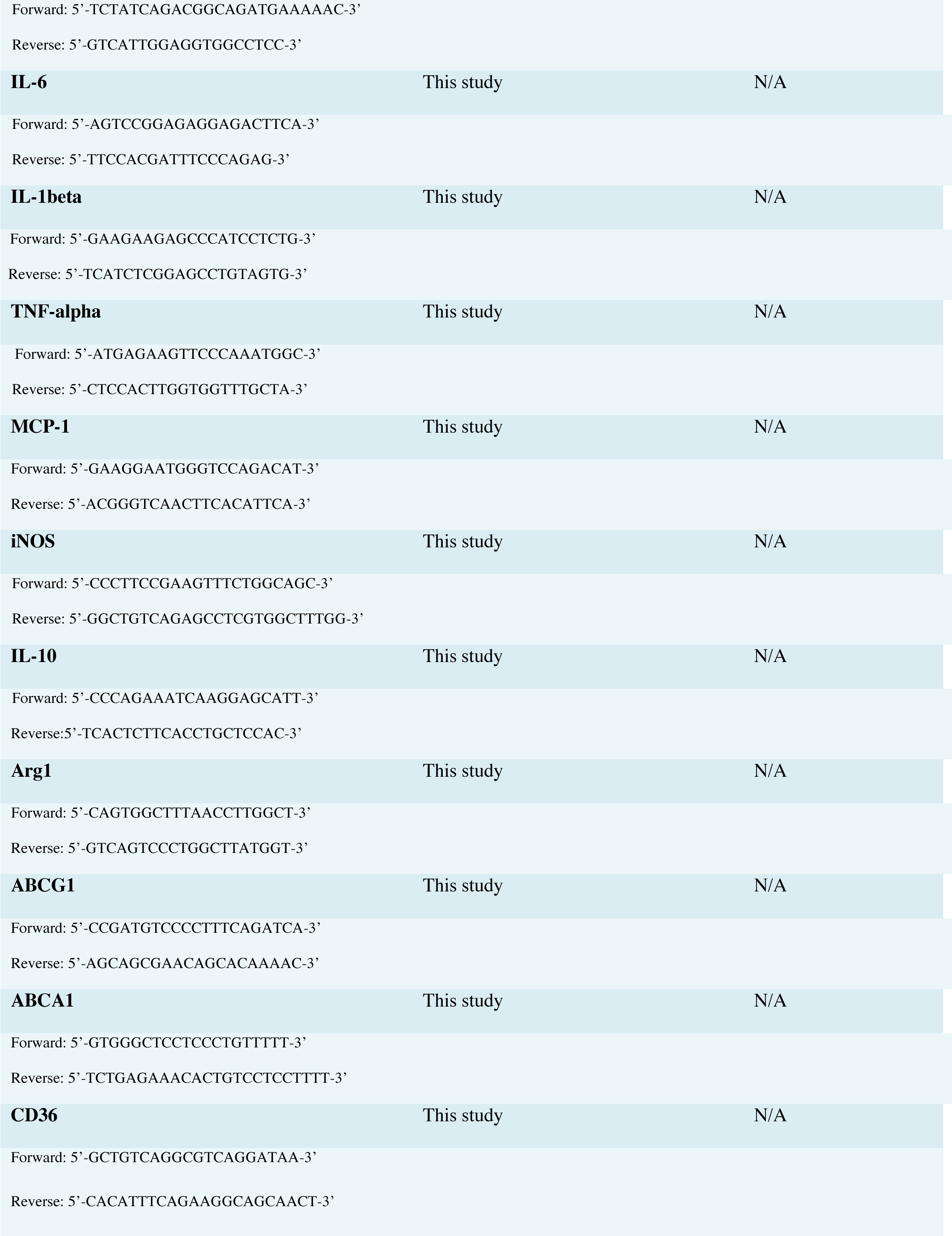
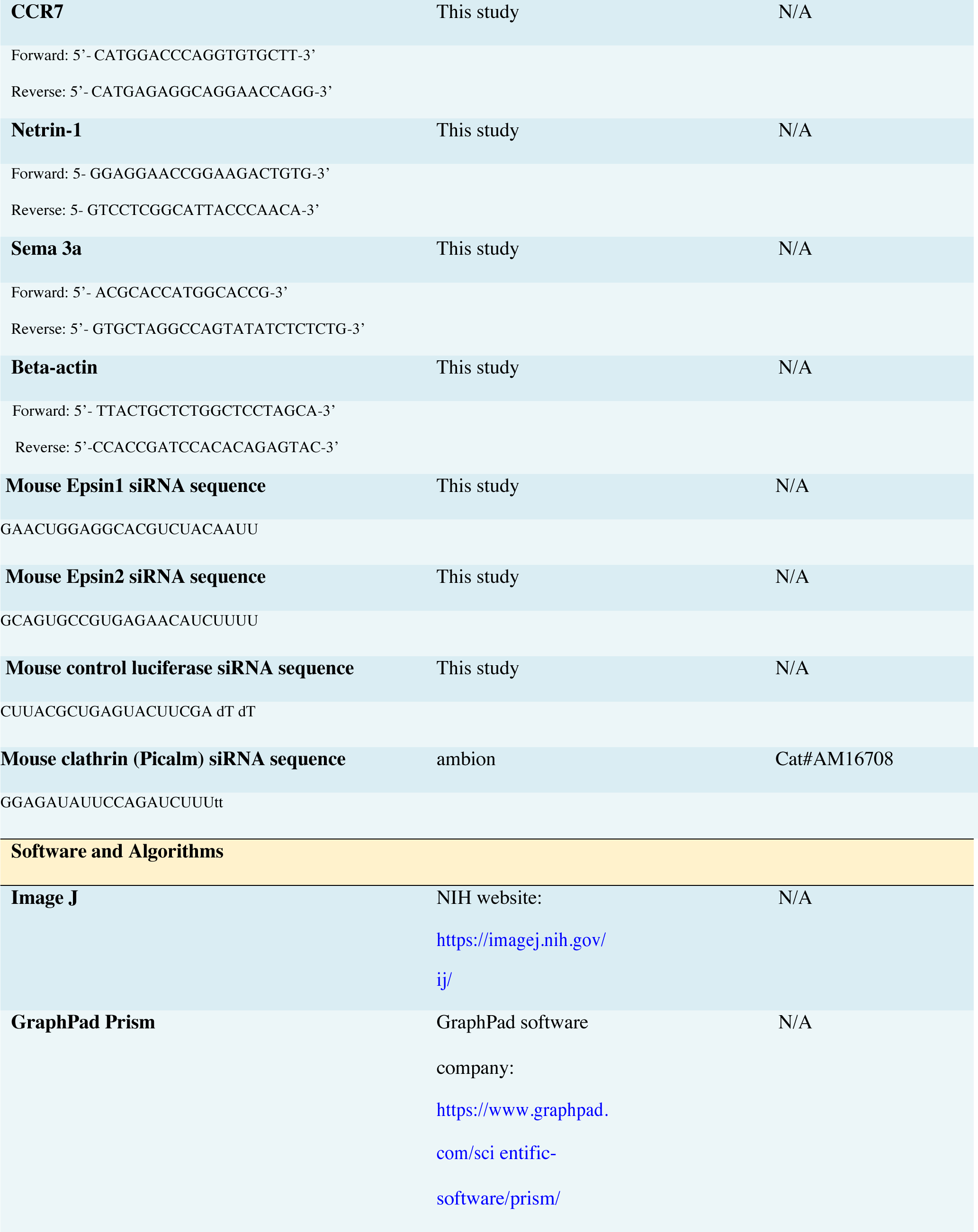
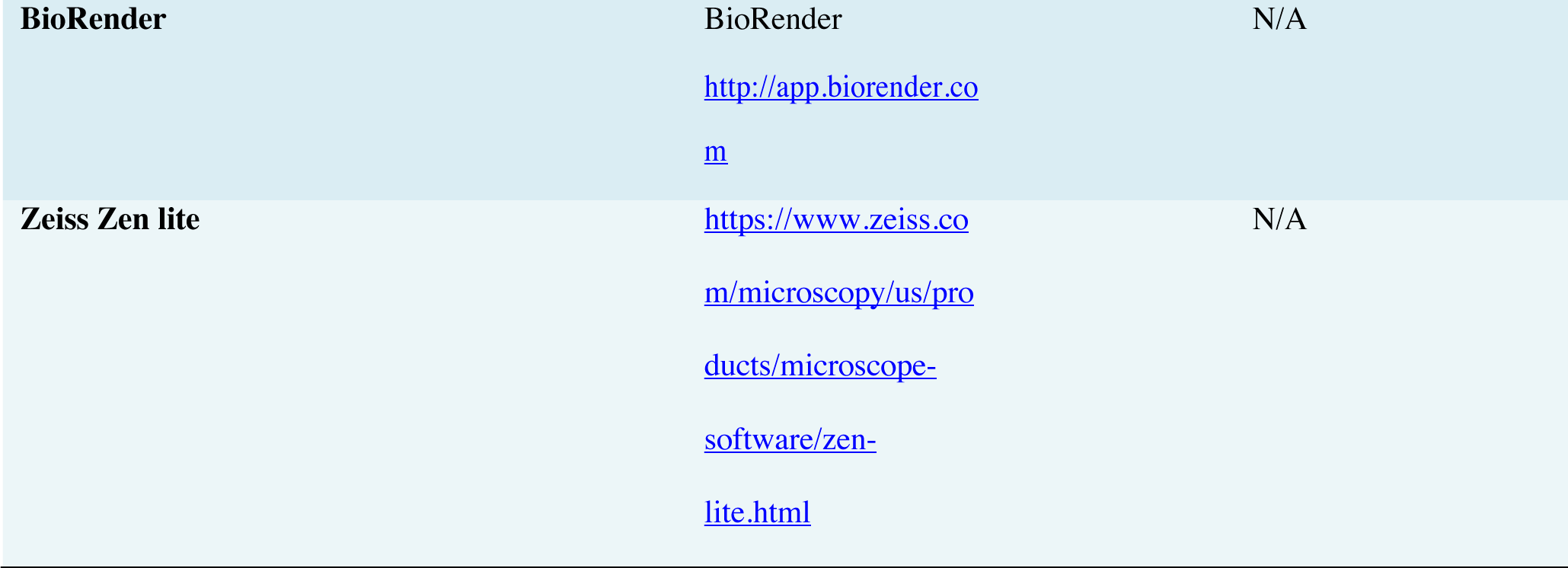
List of reagents and animal models.

